# Genomic time-series data show that gene flow maintains high genetic diversity despite substantial genetic drift in a butterfly species

**DOI:** 10.1101/2021.04.21.440845

**Authors:** Zachariah Gompert, Amy Springer, Megan Brady, Samridhi Chaturvedi, Lauren K. Lucas

**Affiliations:** Department of Biology, Utah State University, Logan, UT, 84322, USA; Ecology Center, Utah State University, Logan, UT, 84322, USA; Department of Organismic & Evolutionary Biology, Harvard University, Cambridge, MA, 02138, USA

**Keywords:** effective population size, population-genetic time series, genetic drift, gene flow, *Lycaeides*

## Abstract

Effective population size affects the efficacy of selection, rate of evolution by drift, and neutral diversity levels. When species are subdivided into multiple populations connected by gene flow, evolutionary processes can depend on global or local effective population sizes. Theory predicts that high levels of diversity might be maintained by gene flow, even very low levels of gene flow, consistent with species long-term effective population size, but tests of this idea are mostly lacking. Here, we show that *Lycaeides* butterfly populations maintain low contemporary (variance) effective population sizes (e.g., ∼200 individuals) and thus evolve rapidly by genetic drift. Contemporary effective sizes were consistent with local census populations sizes. In contrast, populations harbored high levels of genetic diversity consistent with an effective population size several orders of magnitude larger. We hypothesized that the differences in the magnitude and variability of contemporary versus long-term effective population sizes were caused by gene flow of sufficient magnitude to maintain diversity but only subtly affect evolution on generational time scales. Consistent with this hypothesis, we detected low but non-trivial gene flow among populations. Furthermore, using population-genomic time-series data, we documented patterns consistent with predictions from this hypothesis, including a weak but detectable excess of evolutionary change in the direction of the mean (migrant gene pool) allele frequencies across populations, and consistency in the direction of allele frequency change over time. The documented decoupling of diversity levels and short-term change by drift in *Lycaeides* has implications for our understanding of contemporary evolution and the maintenance of genetic variation in the wild.

## Introduction

Patterns of genetic variation within and among populations are observable outcomes of demographic and evolutionary processes. Despite tremendous advances in our ability to model and reconstruct historical processes from population genetic patterns (e.g., Hey & Nielsen, 2004; Gutenkunst *et al*., 2009; Green *et al*., 2010; Li & Durbin, 2011; Speidel *et al*., 2019; Stern *et al*., 2021), connecting pattern to process remains a challenge. Similar patterns can result from multiple processes (Kruuk *et al*., 1999; Sousa *et al*., 2011; Yang *et al*., 2017; Harris *et al*., 2018; Lawson *et al*., 2018). Patterns of allele frequency change over time may provide additional information about demographic and evolutionary processes. Such population-genetic time-series data might be especially useful for understanding contemporary evolutionary processes and eco-evolutionary dynamics (Messer *et al*., 2016). Indeed, time-series data were central to early studies of evolution in natural populations (Fisher & Ford, 1947; Kettlewell, 1958; Ford, 1977; Mueller *et al*., 1985), and play a key role in studies of experimental evolution (Burke *et al*., 2010; Graves Jr *et al*., 2017; R^ego *et al*., 2019; Langmüller & Schlötterer, 2020) and recent attempts to reconstruct human history (reviewed in Pääbo *et al*., 2004; Slatkin & Racimo, 2016). Nonetheless, population-genomic data from natural populations sampled repeatedly through time remain relatively rare (but see, e.g., Bergland *et al*., 2014; Brüniche-Olsen *et al*., 2016; Ryan *et al*., 2018; Bi *et al*., 2019).

Such temporal genomic data are especially informative about effective population size (*N_e_*; e.g., Krimbas & Tsakas, 1971; Waples, 1989; Palstra & Ruzzante, 2008; Gilbert & Whitlock, 2015), which is a factor with substantial effects on the evolutionary process (Wright, 1931). In particular, effective population size determines how fast allele frequencies change by genetic drift, the efficacy of natural selection, and expected genetic diversity levels for neutral loci (Charlesworth, 2009; Lanfear *et al*., 2014; Wang *et al*., 2016). However, various definitions of effective population size exist, with different ones capturing different aspects of random genetic drift and sometimes different spatial or temporal scales (Hill, 1981; Charlesworth, 2009; Walsh & Lynch, 2018). For example, the coalescent effective population size is defined based on the coalescent process and is affected by demographic and evolutionary events from the present back to the common ancestor of set of gene copies (Nordborg & Krone, 2002; Sjodin *et al*., 2005; Wakeley & Sargsyan, 2009). Both the coalescent effective population size and eigenvalue effective population size provide a basis for the expectation that neutral pairwise nucleotide diversity is *π* = 4*N_e_µ*, where *µ* is the mutation rate (Kimura, 1983; Ryman *et al*., 2019). In contrast, the variance effective populations is defined by the rate of random genetic change over one or several generations (Nei & Tajima, 1981; Crow & Denniston, 1988; Jorde & Ryman, 1995; Do *et al*., 2014). It thus represents a contemporary or short-term effective population size, and can be directly inferred from temporal population genetic data (though other methods using samples from a single time point also exist; reviewed in Gilbert & Whitlock, 2015; Wang *et al*., 2016). Several factors, including temporal variation in population size, can cause these effective population sizes to differ.

Population subdivision further complicates the concept of effective population size (Ryman *et al*., 2019). For example, theory indicates that nucleotide diversity (*π*) in each subpopulation or deme should be the same as it would be for a panmictic population of the same total size as long as the subpopulations are connected by gene flow (Nei & Taka-hata, 1993; Lande, 1992). Any level of gene flow is considered sufficient (i.e., any *m >* 0), because with lower rates of gene flow subpopulations diverge more and thus the effect of gene flow when it does occur is increased (Whitlock & Barton, 1997). Thus, with gene flow, the coalescent effective population size for each subpopulation should be equivalent to the species- or metapopulation-level effective population size, and consequently high levels of genetic diversity are expected in each subpopulation. In contrast, low rates of gene flow might have relatively little effect on the short-term rates of evolution by drift, as captured by the (contemporary) variance effective population size (but see Wang & Whitlock, 2003). Consequently, short-term change by drift should reflect local (within deme) effective population size (Serbezov *et al*., 2012; Gilbert & Whitlock, 2015). Thus, theory suggests that loss of diversity and change by drift represent distinct facets of stochastic evolution and might be decoupled in subdivided populations (Ryman *et al*., 2019). But because most studies consider either long-term diversity-based estimates of *N_e_* (e.g., Nei & Graur, 1984; Brown *et al*., 2004; Leffler *et al*., 2012; Feng *et al*., 2017; Capblancq *et al*., 2020) or short-term estimates of contemporary *N_e_* (e.g., Frankham, 1995; Serbezov *et al*., 2012; Gompert & Messina, 2016a; Pazmiño *et al*., 2017; Nunziata & Weisrock, 2018; Rêgo *et al*., 2019) but not both, explicit tests of this prediction are lacking. This knowledge gap has important implications for explaining diversity levels and patterns of evolution in nature, and for informing applied conservation genetics programs.

Here, we combine demographic and population genomic data from ten *Lycaeides idas* butterfly (sub)populations sampled multiple times over a span of five generations (2013–2017) to (i) describe temporal patterns of genome-wide evolutionary change and (ii) determine the relationship between diversity levels (i.e., long-term *N_e_*) and rates of allele frequency change (i.e., short-term *N_e_*). We specifically test the hypothesis that gene flow maintains high diversity levels despite substantial short-term evolution by drift, as predicted by theory (e.g., Ryman *et al*., 2019).

*Lycaeides idas* butterflies occur in northern and western North America (Scott, 1986). These herbivorous insects feed on legumes, especially *Astragalus* and *Lupinus* (Gompert *et al*., 2013). These butterflies have a patchy distribution tied to the distribution of their legume hosts (Gompert *et al*., 2010). Field studies suggest dispersal among host patches is limited (i.e., dispersal rarely exceeds 500 m Knutson *et al*., 1999; U.S. Fish and Wildlife Service, 2003). Despite limited dispersal, genetic differentiation among conspecific populations is low (Gompert *et al*., 2014b), and hybridization with other *Lycaeides* species has been and remains common (Gompert *et al*., 2006; Chaturvedi *et al*., 2020). In fact, many of the populations we study here are ancient hybrids with the majority of their genome from *L. idas* but some segments from *L. melissa* (Gompert *et al*., 2012; Chaturvedi *et al*., 2020). This is unlikely to affect our current analyses and we do not distinguish these populations from putatively non-admixed *L. idas* in this study.

Previous population genomic studies described spatial patterns of genetic variation in this group of butterflies and have used such data to make inferences about past evolutionary processes such as hybridization and host adaptation (e.g., Gompert *et al*., 2012, 2014b; Chaturvedi *et al*., 2018, 2020). However, this prior work did not include repeated sampling across time, and thus lacked the ability to use temporal patterns of change to augment inferences of evolutionary processes. By using such data here, we show that *L. idas* populations evolve rapidly by drift, and consequently have low variance effective population sizes (*Ne* ≈ 200), but nonetheless exhibit little genetic differentiation and harbor substantial genetic diversity consistent with much higher long-term effective population sizes. We find evidence that these populations are connected by modest levels of gene flow, which are sufficient to maintain this diversity, but have low or negligible effects on rates of drift. Nonetheless, gene flow, possibly combined with selection, does have a weak but detectable effect on the direction of evolutionary change across the genome.

## Methods

### Genetic data collection

We sampled 1536 adult *Lycaeides* butterflies from ten localities in each of three or four years between 2013 and 2017 (Table 1, Fig. 1A). Butterflies from within Yellowstone National Park and Grand Teton National Park were collected in accordance with US national park study permits YELL-05924 and GRTE-00285, respectively. Genomic DNA from each butterfly was purified using Qiagen’s DNeasy Blood and Tissue kit in accordance with the manufacturer’s recommendations (Cat. No. 69581; Qiagen Inc., Valencia, CA, USA). We then created reduced complexity, double-digest restriction fragment-based DNA library for each individual following methods outlined in Gompert *et al*. (2014b) (see ‘Preparing the GBS libraries’ in the Online Supplemental Materials [OSM] for details). These DNA libraries were sequenced at the University of Texas Genomic Sequencing and Analysis Facility (Austin, TX, USA). Library preparation and sequencing took place in two batches, with 768 samples sequenced with an Illumina HiSeq 4000 (1×100 base pair [bp] reads) in 2016 (four lanes of sequencing) and 768 samples sequenced on a HiSeq 2500 (1×100 bp reads) in 2018 (four lanes of sequencing) (Table 1). In total, we generated ∼2.5 billion 100 bp DNA sequences. As detailed below and in the OSM, care was taken during data processing and analyses to avoid possible confounding batch effects.

**Figure 1:**
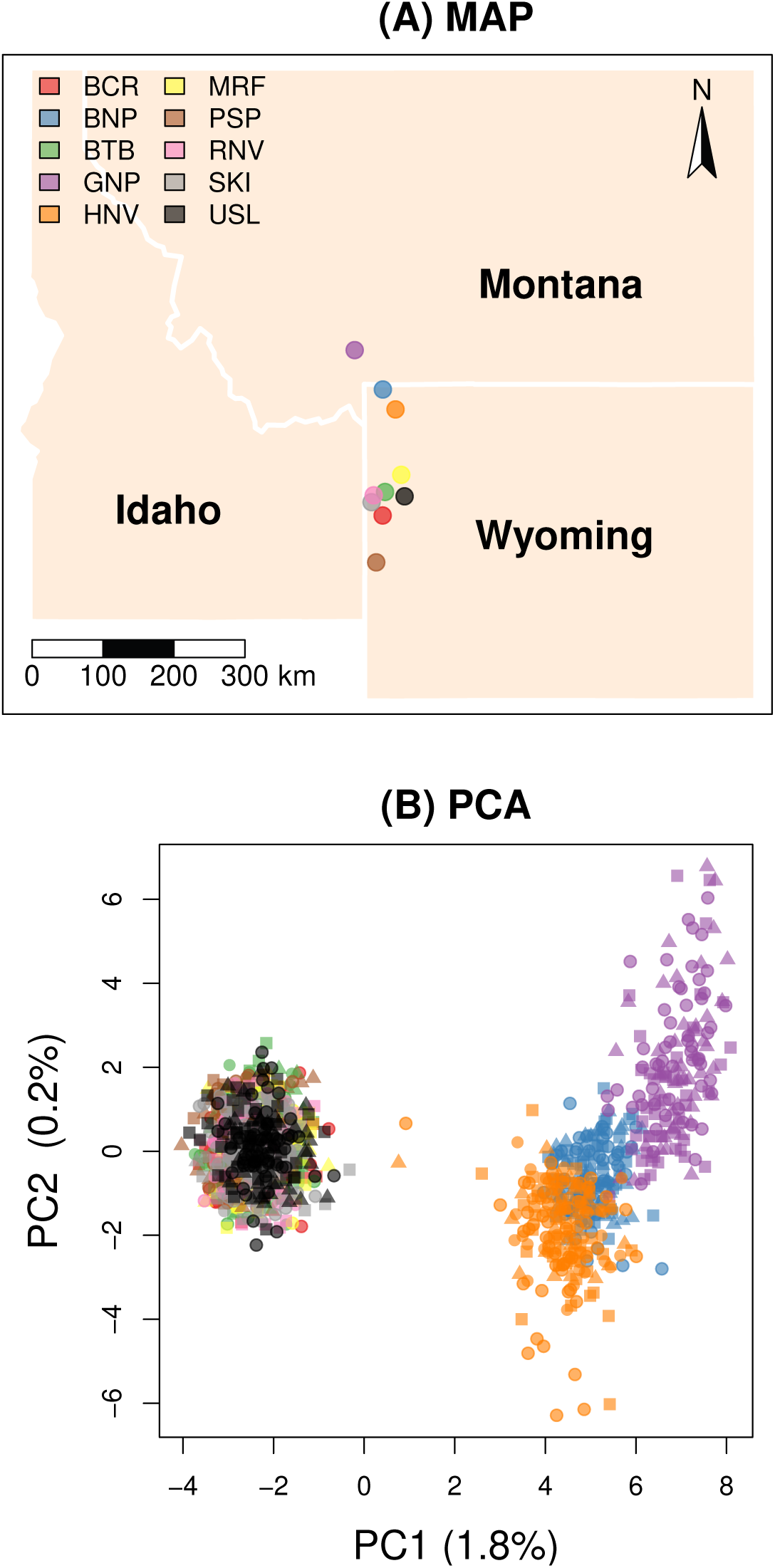
Sample localities (A) and statistical summary of genetic variation in space and time based on a principal component analysis of estimated genotypes (B). (A) The map shows the locations of the populations studied in the Rocky Mountains of the western USA. State names are given. The colored symbols in (B) denote individuals, with colors designating different populations and symbols indicating the year: 2013 = square, 2014 = small circle, 2015 = triangle and 2017 = large circle.

**Table 1:** Population locations and sample sizes (*N*) for population-genomic analyses for each year. Samples were processed in two batches (1 versus 2 below).

| ID | Site name | Lat. ($^{\circ}$N) | Long. ($^{\circ}$W) | Year | Batch | $N$ |
| --- | --- | --- | --- | --- | --- | --- |
| PSP | Periodic Springs | 42.748 | 110.840 | 2013 | 1 | 49 |
|  |  |  |  | 2015 | 1 | 38 |
|  |  |  |  | 2017 | 2 | 49 |
| BCR | Bull Creek | 43.341 | 110.725 | 2013 | 1 | 49 |
|  |  |  |  | 2015 | 1 | 41 |
|  |  |  |  | 2017 | 2 | 49 |
| BTB | Blacktail Butte | 43.638 | 110.682 | 2013 | 1 | 49 |
|  |  |  |  | 2014 | 2 | 50 |
|  |  |  |  | 2015 | 1 | 51 |
|  |  |  |  | 2017 | 2 | 51 |
| USL | Upper Slide Lake | 43.583 | 110.333 | 2013 | 1 | 49 |
|  |  |  |  | 2015 | 1 | 50 |
|  |  |  |  | 2017 | 2 | 59 |
| SKI | Ski Lake | 43.509 | 110.923 | 2013 | 2 | 49 |
|  |  |  |  | 2014 | 2 | 50 |
|  |  |  |  | 2015 | 2 | 27 |
|  |  |  |  | 2017 | 2 | 53 |
| RNV | Rendezvous Mt. | 43.596 | 110.885 | 2013 | 1 | 49 |
|  |  |  |  | 2015 | 1 | 47 |
|  |  |  |  | 2017 | 2 | 32 |
| MRF | Mt. Randolph | 43.855 | 110.392 | 2013 | 1 | 48 |
|  |  |  |  | 2015 | 1 | 50 |
|  |  |  |  | 2017 | 2 | 22 |
| HNV | Hayden Valley | 44.682 | 110.495 | 2013 | 2 | 48 |
|  |  |  |  | 2014 | 2 | 51 |
|  |  |  |  | 2015 | 2 | 24 |
|  |  |  |  | 2017 | 2 | 49 |
| BNP | Bunsen Peak | 44.934 | 110.721 | 2013 | 1 | 49 |
|  |  |  |  | 2015 | 1 | 49 |
|  |  |  |  | 2017 | 2 | 47 |
| GNP | Garnet Peak | 45.432 | 111.225 | 2013 | 1 | 49 |
|  |  |  |  | 2015 | 1 | 51 |
|  |  |  |  | 2017 | 2 | 56 |

After de-multiplexing, we used bwa (version 0.7.17-r1188) to align DNA sequences to the *Lycaeides melissa* genome (Chaturvedi *et al*., 2020) using the aln and samse algorithms (Li & Durbin, 2009). For alignment, we allowed for no more than four mismatches, no more than two mismatches in a 20 bp seed, and only placed sequences with a unique best match. Sequence alignments were compressed, sorted and indexed with samtools (Li *et al*., 2009). We then identified SNPs separately using both samtools (version 1.5) combined with bcftools (version 1.6) and GAKT’s HaplotypeCaller and the GenotypeGVCFs module (version 3.5) (McKenna *et al*., 2010). We retained only the subset of SNPs called by both variant callers and that showed similar allele frequencies in both sequencing batches. See the ‘Variant calling and filtering’ in the OSM for additional details regarding variant calling and filtering (this includes a number of filtering criteria not listed here). We retained 12,886 SNPs for downstream analysis.

### Describing patterns of genetic variation in space and time

We estimated genotypes and allele frequencies for each SNP locus to describe patterns of genetic variation in space and time. Genotypes were inferred using entropy (version 1.2; Gompert *et al*., 2014b; Shastry *et al*., 2021). This program estimates genotypes while accounting for uncertainty caused by limited coverage and sequencing error (as captured by the genotype likelihoods). The model assumes that the allele copies at each SNP locus are drawn from unknown, hypothetical source populations with each individual having a genome with ancestry from some mixture of the source populations. We estimated genotypes assuming two or three source populations, and using the genotype likelihoods from bcftools as input. Estimates were obtained via Markov chain Monte Carlo (MCMC) with three chains, each with 10,000 iterations and a 5000 iteration burn-in. We set the thinning interval to 5. Point estimates of genotypes were obtained as the posterior mean estimate of the number of non-reference alleles, with the posterior summarized across chains and numbers of source populations. We then visualized patterns of genetic variation using a principal component analysis (PCA) via the prcomp function in R, with the centered but not scaled genotype estimates as input (i.e., the covariance matrix). A distance-based redundancy analysis (RDA) was then used to quantify the extent that genetic variation was partitioned by population and year (see, e.g., Driscoe *et al*., 2019). The RDA was conducted in R with the adonis function from vegan (version 2.5-7) and using Euclidean distances (Oksanen *et al*., 2020). Statistical significance of predictors was assessed based on 100 permutations, with permutations across generations constrained to be within populations.

Next, we used estpEM (version 0.1) to estimate allele frequencies for each population and year (Soria-Carrasco *et al*., 2014). This program uses an expectation-maximization algorithm to obtain maximum likelihood estimates of allele frequencies while accounting for uncertainty in genotypes as captured by the genotype likelihoods (Li, 2011; Soria-Carrasco *et al*., 2014). We set the maximum number of iterations for this algorithm to 50 and set the convergence tolerance to 0.001. We assumed females contributed only one copy of any Z-linked loci when estimating allele frequencies, which is expected as females are the heterogametic sex in butterflies (ZW). We then quantified genetic differentiation among populations and years based on F_ST_. This was computed as 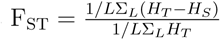, where *H_T_* and *H_S_* are the expected heterozygosities for the combined and individual population or generation pairs and the sums are over the *L* SNPs. Additionally, we quantified change in time for individual SNPs by calculating the allele frequency differences between years. These calculations were done in R.

### Estimating census population size, contemporary *N_e_* and diversity levels

We estimated census population sizes, contemporary variance effective populations sizes, and genetic diversity levels (a proxy for long-term effective population size) to determine how these distinct metrics of population size differed. Our hypothesis that gene flow maintains diversity despite substantial drift predicts that census and contemporary effective population sizes will vary among populations and will be relatively low, whereas diversity levels will vary less among populations and be consistent with a larger effective population size.

Census population sizes were estimated annually in subsets of populations from 2013 to 2018 with a distance sampling approach (25 estimates total) (Table S1). Distance sampling involves counting individuals and recording their distance from a transect line or point (Buckland *et al*., 2001). This distance information is used to estimate a detection function that accounts for imperfect detection away from the transect line. We estimated population densities (adult butterflies per square kilometer) using the distsamp function in the unmarked R package (version 1.0.1; Fiske & Chandler, 2011). We binned the detection distances of butterflies into 1 meter bins prior to analysis (e.g., 0 to 1 m, 1 to 2 m, etc.). We used a half-normal detection function and estimated the detection function and density model parameters using maximum likelihood (Royle *et al*., 2004). This model assumes the latent transect-level abundance distribution is Poisson and that the detection process is multinomial with a different detection probability for each distance class or bin. We then estimated population size by first multiplying density by the area of habitat (km^2^) and then by three because adult *L. idas* live for about a week and the bulk of the adult flight occurs over a ∼3-week period (personal observation). A mark-release-recapture study was conducted at one site in 2018 (BTB) as an independent estimate of census population size. See ‘Estimating census population sizes’ in the OSM for additional details.

We estimated the contemporary variance effective population size at each site based on the magnitude of allele frequency change between 2013 and 2017 (our first and last samples). We did this using a Bayesian bootstrap method, as described in Gompert (2016) and Gompert & Messina (2016a) (also see Jorde & Ryman, 2007; Foll *et al*., 2015). We focused on change over the largest time interval, as this is less sensitive to uncertainty associated with sampling error (i.e., change by drift compounds across generations, whereas sampling error is unique to each sample). This approach assumes evolution occurred solely by genetic drift, but selection on a modest number of SNPs should not have much of an effect on the estimates of *N_e_*. Estimates of contemporary variance *N_e_* were obtained using varne (version 0.1) with 1000 bootstrap replicates (Gompert & Messina, 2016b).

Next, we estimated genetic diversity levels in each population and generation. We did this in the context of the neutral theory expectation that *θ* = 4*N_e_µ*, where *N_e_* is the (long-term) effective population size and *µ* is the mutation rate, and where *θ* = *π* (i.e., nucleotide diversity). We estimated diversity levels using ANGSD (version 0.933-71-g604e1a4), which uses bam alignment files as input and accounts for uncertainty in genotypes and in whether individual nucleotides are variable (Korneliussen *et al*., 2014). We ran this analysis for each sample (population by generation combination) with genotype likelihoods computed in the same way as samtools or GATK (-GL set to 1 or 2). We only used reads with minimum mapping quality of 30 and bases with minimum quality of 20, and computations were based on the folded site-frequency spectrum. Females (the heterogametic sex) were excluded when estimating diversity for the Z sex chromosome.

### Estimating gene flow

We next estimated levels of gene flow among the sampled populations to test the hypothesis that gene flow was sufficient to explain the observed differences between contemporary effective population sizes and long-term effective population sizes as captured by diversity levels. Ideally, we would fit a single model with all populations (including unsampled populations) that accounts for the (likely) complex history of divergence and various rates of gene flow among different populations, however, this is not practical. Instead, we used two simpler, complementary approaches with a main aim of obtaining estimates of gene flow that are of the correct order of magnitude at least.

First, we used the diffusion approximation approach implemented in *δ*a*δ*i to estimate rates of gene flow for pairs of populations assuming an isolation with migration model (Hey & Nielsen, 2004; Gutenkunst *et al*., 2009). This method is computationally efficient and thus can be applied to many pairs of populations, and also counts for the fact that populations might not be at drift-migration equilibrium. With this approach, we estimated an ancestral, mutation-scaled effective population size (*θ* = 4*N_anc_µ*), the population split time (*T_split_*), population growth parameters (*ν*_1_ and *ν*_2_), and migration rates *M*_12_ and *M*_21_ from the joint site frequency spectrum for each pair of populations (see Fig. S1). Joint frequency spectra for each pair of populations were first estimated using ANGSD (version 0.933-71-g604e1a4) using the same approach and settings described above for estimating single-population site frequency spectra (Korneliussen *et al*., 2014). For this, we based our inferences only on the 2017 samples (which were processed in a single batch for all populations) and only used the samtools method for calculating genotype likelihoods. We then used *δ*a*δ*i to estimate migration rates. Site frequency spectra were down-sampled to 50% to account for missing data. Fifteen rounds of numerical optimization were attempted for each population pair, each involving 50 iterations. See ‘Demographic inference with *δ*a*δ*i’ in the OSM for additional details.

Second, we estimated gene flow among the six southernmost populations by fitting a Bayesian F-model (e.g., Gaggiotti & Foll, 2010) to the allele frequency data from 2017. This statistical model can be used to approximate various demographic processes, including migration-drift equilibrium in an island model (Balding & Nichols, 1995; Rannala & Hartigan, 1996; Nicholson *et al*., 2002; Falush *et al*., 2003; Gompert *et al*., 2012, ; see ‘Quantitative basis for assuming drift-migration equilibrium’ in the OSM and Fig. S4 for evidence in support of this assumption). Thus, this approach makes equilibrium assumptions, but also allows us to fit a single model for a set of populations. We focus on the six southernmost populations because we find little to no evidence for isolation-by-distance across this set of populations (i.e., patterns of differentiation are consistent with an island model; see Results for details). We fit the Bayesian F-model using Hamiltonian Monte Carlo via the rstan interface with stan (Stan Development Team, 2021, 2019). We assumed a single migration rate for all populations and SNP loci. We placed a Cauchy prior on the number of migrants per generation (location parameter = 0, scale parameter = 10, truncated at 0 and 50) and a beta prior on the migrant allele frequency (*α* and *β* both set to 0.5, which corresponds with Jeffrey’s prior). Parameter estimates were based on samples from four chains, each consisting of 1000 iterations as a burn-in and 2000 sampling iterations.

### Using simulations to test the effects of gene flow

We next used simulations to verify that gene flow of the order detected here could indeed cause the persistence of high diversity levels (suggestive of a high long-term *N_e_*) despite much lower contemporary variance effective population sizes for each population and thus high rates of short-term evolution by genetic drift. This was at least in part expected, as past theory suggests that any non-zero level of gene flow should (in the long-term) maintain as much variation as would be expected for a single, large panmictic population (Whitlock & Barton, 1997). But diversity could vary over time in a way not captured by analytical theory (i.e., diversity could decline for long periods of time between very rare gene flow events), hence our desire to examine this issue with simulations.

To this end, we simulated evolution forward in time under a Wright-Fisher model with SLiM3 (Haller & Messer, 2019). We considered a spatial matrix of 36 populations arranged in a 6×6 grid, each with a variance *N_e_* of 173 (the mean for *L. idas*). We chose this number of populations as it provided a substantial contrast between demic and total population size, but was also computationally tractable. We assumed that these were descended from a single large, panmictic population of 6228 individuals (i.e., 173×36). Migration occurred between neighboring demes with *m* = 0.001 (low migration) or *m* = 0.01 (high migration); we also simulated the case of no migration (*m* = 0). Our high migration rate (*m* = 0.01) corresponds approximately with our migration rate estimate from the Bayesian F-model, and we consider the low migration rate (*m* = 0.001) as a reasonable lower bound on the minimum rate of gene flow in this system. Simulations were run for an initial 100,000 generations before the population split, followed by 300,000 simulations after the split. See ‘Simulations with SLiM3’ in the OSM for additional details.

### Testing for short-term effects of gene flow

Having shown that gene flow can maintain high diversity levels despite low contemporary *N_e_* (see Results), we next asked whether gene flow had detectable effects on short-term patterns of genome-wide allele frequency change. We focused on specific contrasts where predictions can be made based on the hypothesis of contemporary effects of gene flow and where batch sequencing effects could be avoided. In some cases, polygenic selection is a viable alternative hypothesis for the predicted patterns. We make note of this when introducing these predictions below, and return to this issue in the Discussion.

We first focused on the set of six southernmost populations where the samples from 2013 and 2015 were all sequenced in a single batch (i.e., BCR, BTB, MRF, PSP, RNV, and USL; Table 1, Fig. 1A). Here, we tested the prediction that gene flow among this set of populations should cause change towards the mean allele frequency. In other words, if an allele is less common in a population than it is on average across a set of populations, gene flow should cause the allele to increase in that population. To test this prediction, we determined the correlation (across SNPs) between the sign of the allele frequency difference for each population in 2013 relative to the mean (i.e., sign of 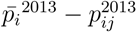) and the sign of change between 2013 and 2015 (i.e., sign of 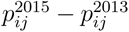) (where *i* denotes a SNP locus and *j* a population). Because sampling error could induce a spurious correlation between these metrics, we determined a null distribution by randomly permuting allele frequencies between years (2013 versus 2015) for each SNP and repeating these calculations (1000 permutations were used to generate a null distribution).

We next focused on the two populations for which every sample was part of the same sequencing batch, HNV and SKI. Here, we predicted that gene flow from other populations would cause consistent changes in allele frequency over multiple generations. This pattern of consistency would also be expected with directional polygenic selection (Buffalo & Coop, 2019). For each population, we used allele frequency estimates from the four temporal samples (2013, 2014, 2015, and 2017) to compute change over each of three successive intervals. We then determined the proportion of SNPs where change was in the same direction over all three intervals. We compared this to null expectations by again randomly permuting allele frequencies across years; this was done 1000 times.

Lastly, we tested the prediction that gene flow should cause an excess of evolutionary change at SNP loci in the same direction as they are being affected by similar immigrant pools. Notably, this pattern would also be consistent with polygenic selection acting similarly across space. We again focused on the six southernmost populations with 2013 and 2015 samples from a single sequencing batch for this, and considered all 15 pairwise comparisons between pairs of these populations (BCR, BTB, MRF, PSP, RNV, and USL; see Table 1 and Fig. 1A). For each pair, we determined the proportion of SNPs where change between 2013 and 2015 was in the same direction. We then generated null expectations by randomly permuting allele frequencies between years for each population and re-calculating the proportion with change in the same direction. This was again repeated 1000 times.

## Results

### Patterns of genetic variation in space and time

Genetic differentiation among butterfly populations was low, but still exceeded genetic differentiation through time. Specifically, in a PCA ordination of genetic variation at the 12,886 SNPs, butterflies clustered by population, with some genetic differentiation between the more southern populations (PSP, BCR, BTB, USL, SKI, RNV, and MRF) and more northern populations (HNV, BNP, and GNP) (Fig. 1). No temporal structure was apparent in the PCA analysis. Likewise, with RDA, population explained 2.5% of the total variation, whereas year explained only 0.2% (both *P <* 0.01). Using the data from 2017 (all one sequencing batch), the average genetic differentiation between pairs of populations was F_ST_ = 0.014 (minimum = 0.006, maximum = 0.026), with similar results in other years (e.g., for 2013 mean F_ST_ = 0.013, minimum = 0.007, maximum = 0.024). Estimates of F_ST_ between years were even lower, with an average of 0.009 (minimum mean across years for a population = 0.007, maximum = 0.010).

Consistent with low levels of temporal genetic structure, the mean allele frequency change between contiguous generations was between 0.018 an 0.024 (mean across all comparisons = 0.020; Figs. 2, S2). With that said, some genomic regions or SNPs showed more substantial allele frequency change. For example, we documented slightly more substantial change for Z-linked SNPs (mean |Δ*p*|, Z = 0.023, autosomes = 0.020, Z *>* autosomes for all 33 comparisons between contiguous temporal samples). Likewise, some SNPs showed much higher rates of change, with averages across all comparisons of 0.068, 0.12, and 0.37 for the 95th, 99th, and 100th percentiles. For example, a single SNP (G versus C polymorphism at position 1,042,561 on chromosome 8) exhibited a decrease in allele frequency of 0.34 between 2013 and 2015 in BTB. Similarly, large negative changes were observed in MRF (0.31), BCR (0.37) and GNP (0.12) (Fig. S3). Interestingly, the allele frequency at this SNP for all of these sites had increased again by 2017.

**Figure 2:**
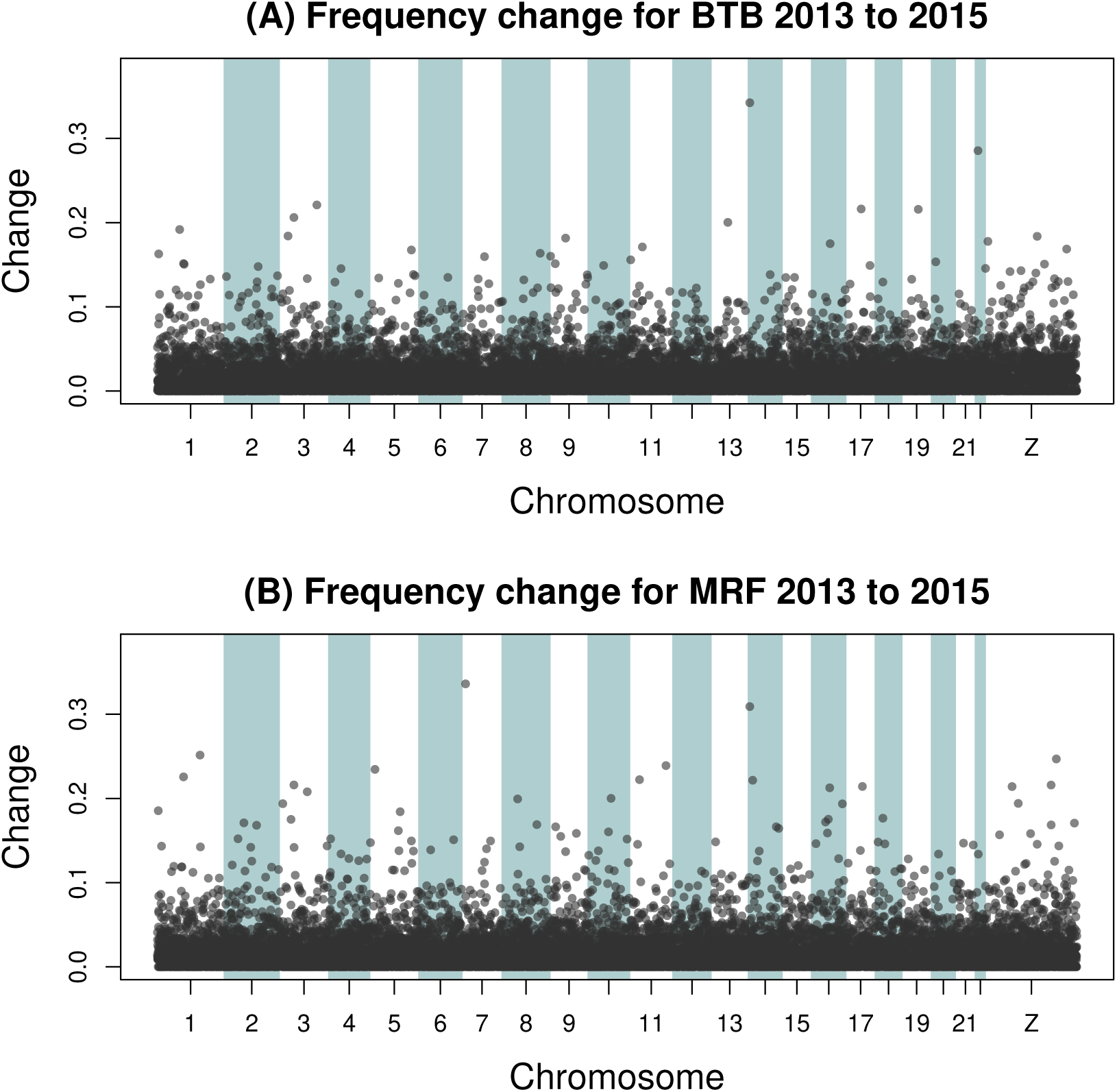
Manhattan plots show patterns of allele frequency change between 2013 and 2015 for two populations–BTB (A) and MRF (B). Points denote the absolute allele frequency difference for each of 12,886 SNPs between 2013 and 2015 along each of the 22 autosomes and Z sex chromosome.

### Census population size, contemporary *N_e_* and diversity levels

Census population size estimates varied from 343 to 5291, with many sites showing evidence of variation in size across years (Table S1, Fig. 3). We failed to detect an effect of host plant abundance on population densities or total census population sizes (linear regression, *P* = 0.68 and *P* = 0.45, respectively), but population size was higher for populations spanning larger geographic areas (linear regression, *β* = 1.5 × 10^−2^, S.E. = 2.2 × 10^−3^, *P* = 8.3 × 10^−7^).

**Figure 3:**
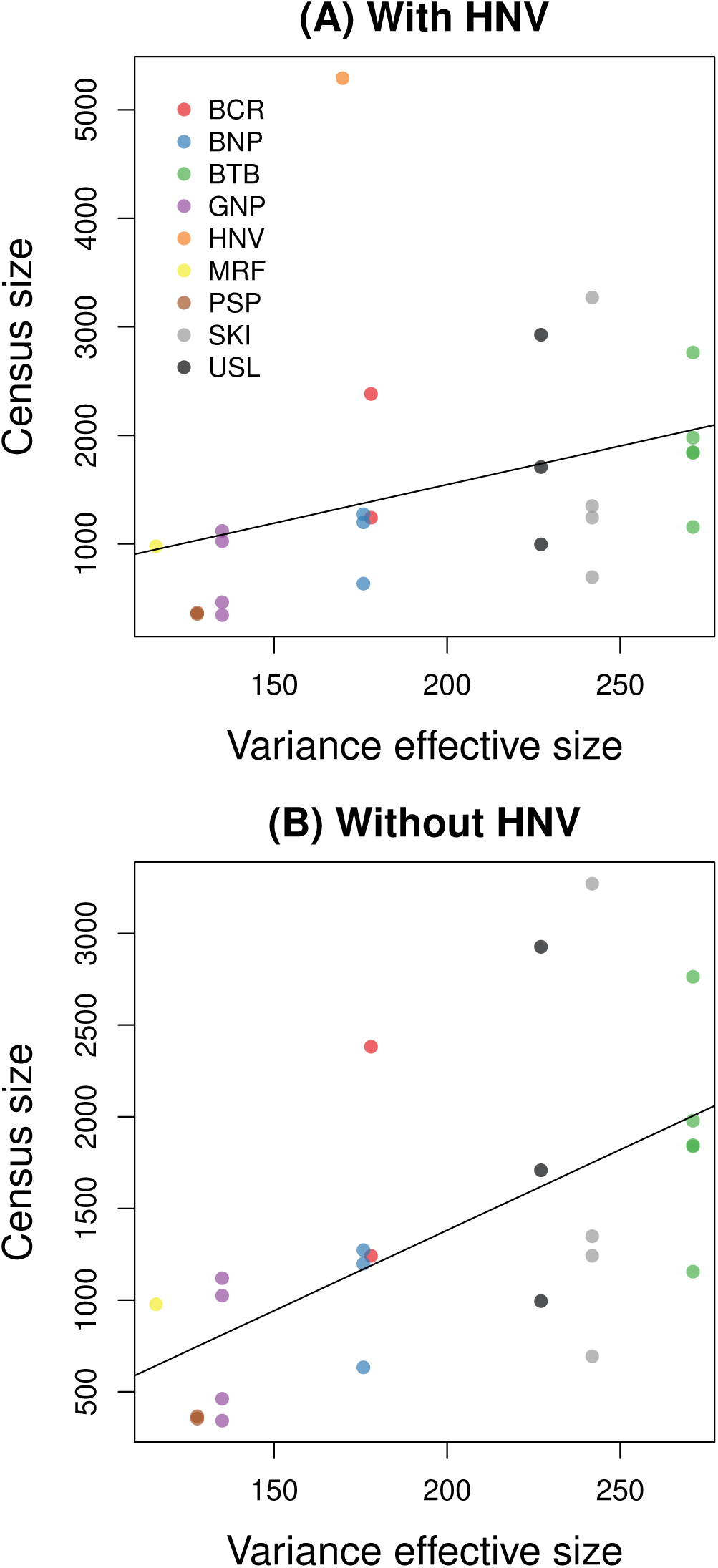
Scatterplots depict the relationship between variance effective population size and census size estimates for nine populations where we obtained reliable census estimates. Results are shown with (A) and without (B) the single, exceptionally large (i.e., outlier) estimate for HNV. Best fit lines from linear regression are shown (with HNV, *r*^2^ = 0.12, *P* = 0.088; without HNV, *r*^2^ = 0.35, *P* = 0.002).

Estimates of variance effective population size based on allele frequency change from 2013 to 2017 were in general lower, with an average of 173 individuals (minimum = 101 for RNV, maximum = 271 for BTB; Fig. 4A). Nonetheless, census and variance effective population sizes were positively correlated (Pearson *r* = 0.35, 95 CIs = -0.05–0.65, *P* = 0.087; excluding a single, exceptionally high estimate of *>*5000 from HNV, *r* = 0.60, *P* = 0.002), and the ratio of census to effective population size was consistent with general expectations from other studies (∼10:1; see Frankham, 1995; Fig. 3).

**Figure 4:**
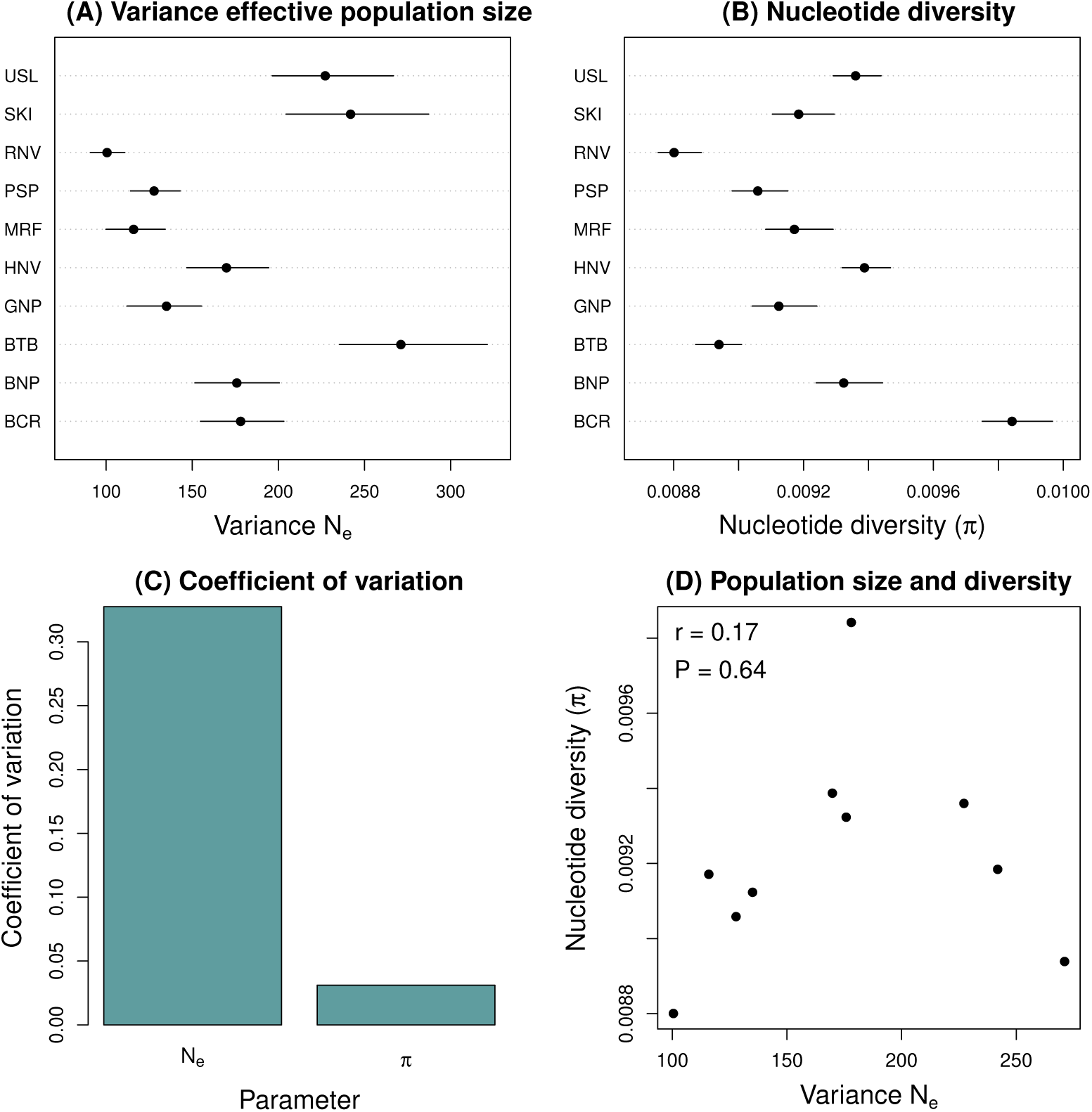
Contemporary variance effective population sizes (*N_e_*) versus nucleotide diversity levels (*π*). Dot plots in panel (A) give Bayesian estimates of variance effective population size based on change from 2013 to 2017. Points denote the median of the posterior and horizontal lines give the 95% equal-tail probability intervals [ETPIs]. Similarly, dot plots in panel (B) report the median and 95% ETPIs for estimates of nucleotide diversity (*π*) based on the 2017 samples. Bars in panel (C) give the coefficient of variation (S.D. relative to the mean) for variance *N_e_* and nucleotide diversity (*π*). The scatterplot in (D) shows the lack of relationship between variance *N_e_* and nucleotide diversity (*π*).

Genetic diversity levels (*π*) were lowest in RNV (0.0088) and highest in BCR (0.0098), with a mean of 0.0092 (Fig. 4B; results reported here are for 2017 samples and samtools genotype likelihood calculations, but other years and methods gave similar results, which are not shown). Given these estimates and assuming *θ* = 4*N_e_µ*, one would have to posit a very high mutation rate of ∼ 1.3 × 10^−5^ to obtain an estimate of effective population size similar to our mean of 173 noted in the preceding paragraph (even assuming a relatively high mutation rate of *µ* = ×10^−8^ yields *N*_e_ = 230,000). Moreover, diversity levels varied much less among populations than variance effective population sizes (coefficient of variation, diversity = 0.031, contemporary *N_e_* = 0.321) (Fig. 4C). We did not detect a relationship between diversity levels and estimates of contemporary variance effective population size (Pearson *r* = 0.17, 95 CIs = -0.52–0.72, *P* = 0.64; Fig. 4D).

### Patterns of gene flow

We obtained reliable estimates of gene flow for 35 out of 45 pairs of populations (78%) under the IM model with *δ*a*δ*i. Specifically, for these pairs model fits met our convergence criteria (see in ‘Demographic inference with *δ*a*δ*i’ the OSM) and joint site-frequency spectra predicted by the fit models were generally consistent with those inferred from the data (Figs. S5–S39). The mean number of migrants per generation between pairs of populations was 0.43 (median = 0.25, minimum ∼0, maximum = 2.30; Fig. 5; see Table S2 for the full set of parameter estimates). As expected, gene flow estimates were higher on average between nearby populations than between farther away populations (Pearson *r* = −0.29). When we instead assumed an equilibrium island model, our estimate of the migration rate was 7.6 migrants per generation (95% equal-tail probability interval [ETPI] = 7.3–8.0). Here, the estimate refers to the total number of migrants from all populations entering each of the six southernmost populations with samples from 2013 and 2015 that were part of the first sequencing batch.

**Figure 5:**
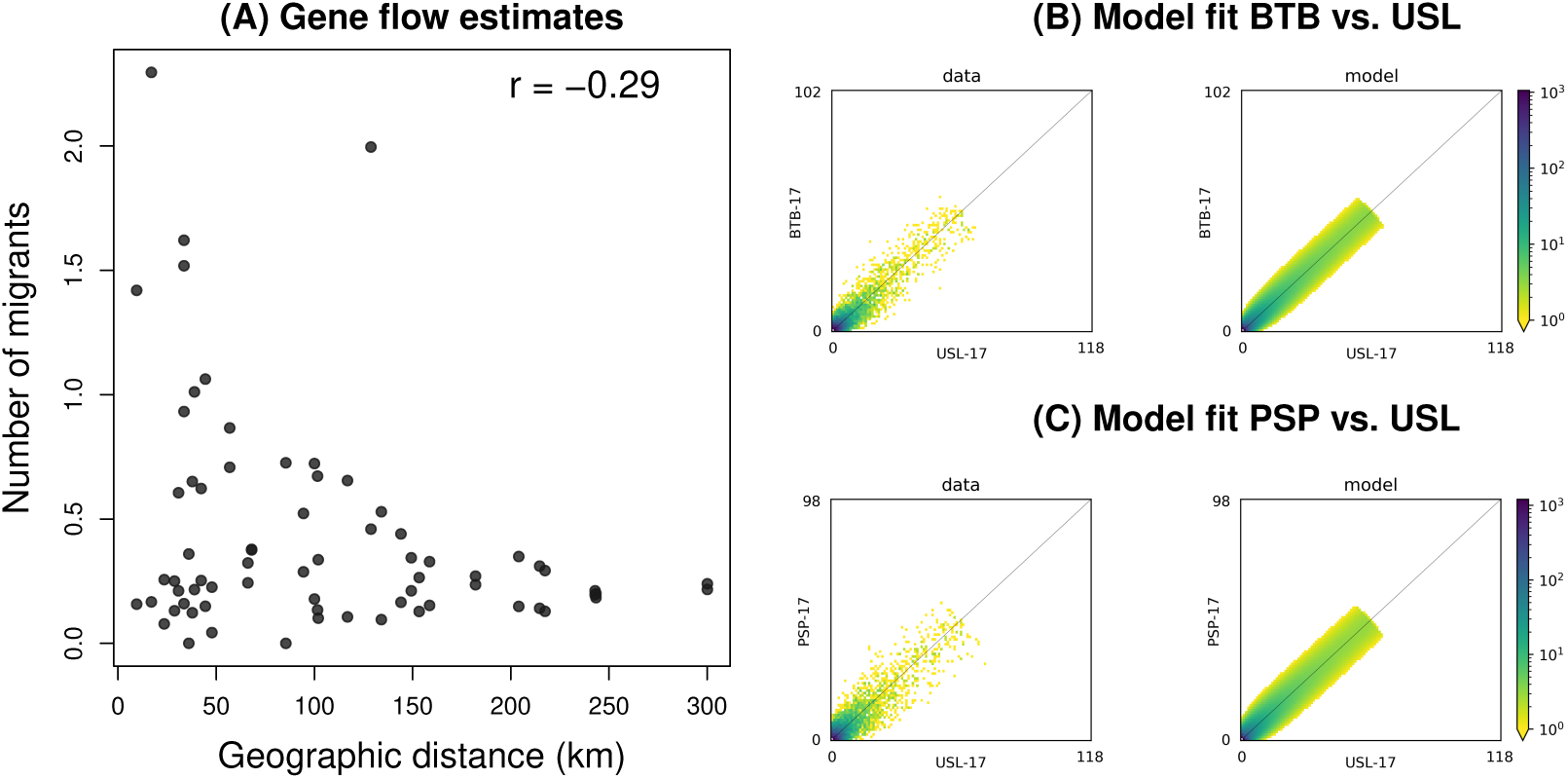
Summary of migration estimates from *δ*a*δ*i. The scatterplot in (A) shows estimates of *Nm* (contemporary number of migrants per generation) between pairs of populations as a function of geographic distance. Results are shown for the 35 out of 45 population pairs that converged (Pearson *r* = -0.29). Panels (B) and (C) show the observed and model-predicted joint site-frequency spectra for two representative pairs of populations (BTB×USL and PSP×USL, respectively). See Figs. S5–S39 for summaries of model fit for additional pairs of populations.

### Effects of gene flow from simulations

SLiM3 simulations showed that with *m* = 0.001 or 0.01 (low or high gene flow), nucleotide diversity (*π*) within populations remained similar to diversity levels for a large, panmictic population (Fig. 6A,B). However, with *m* = 0.001, diversity levels exhibited increased variance over time after the large population split into many isolated populations (e.g. the average S.D. in *π* over time after the populations split was 6.2×10^−4^ for low migration versus 1.9×10^−4^ for high migration; Fig. 6B). In contrast, diversity rapidly declined after the populations split in simulations with no gene flow (Fig. 6C). Despite these differences in diversity levels with versus without ongoing gene flow, estimates of variance effective population size (based on change between generations 150,000 and 150,002, i.e., at a similar temporal scale to our data) were similar to each other and generally in line with the simulated, local (demic) variance effective population sizes (Fig. 6D). Specifically, mean estimate of variance *N_e_* were 206.2 (median = 177.8), 174.7 (median = 164.1) and 170.8 (median = 159.7) for *m* = 0.01, 0.001 and 0, respectively. Thus, in these simulations, gene flow maintained high diversity levels but did not notably affect variance *N_e_* and thus the rate of evolution by drift.

**Figure 6:**
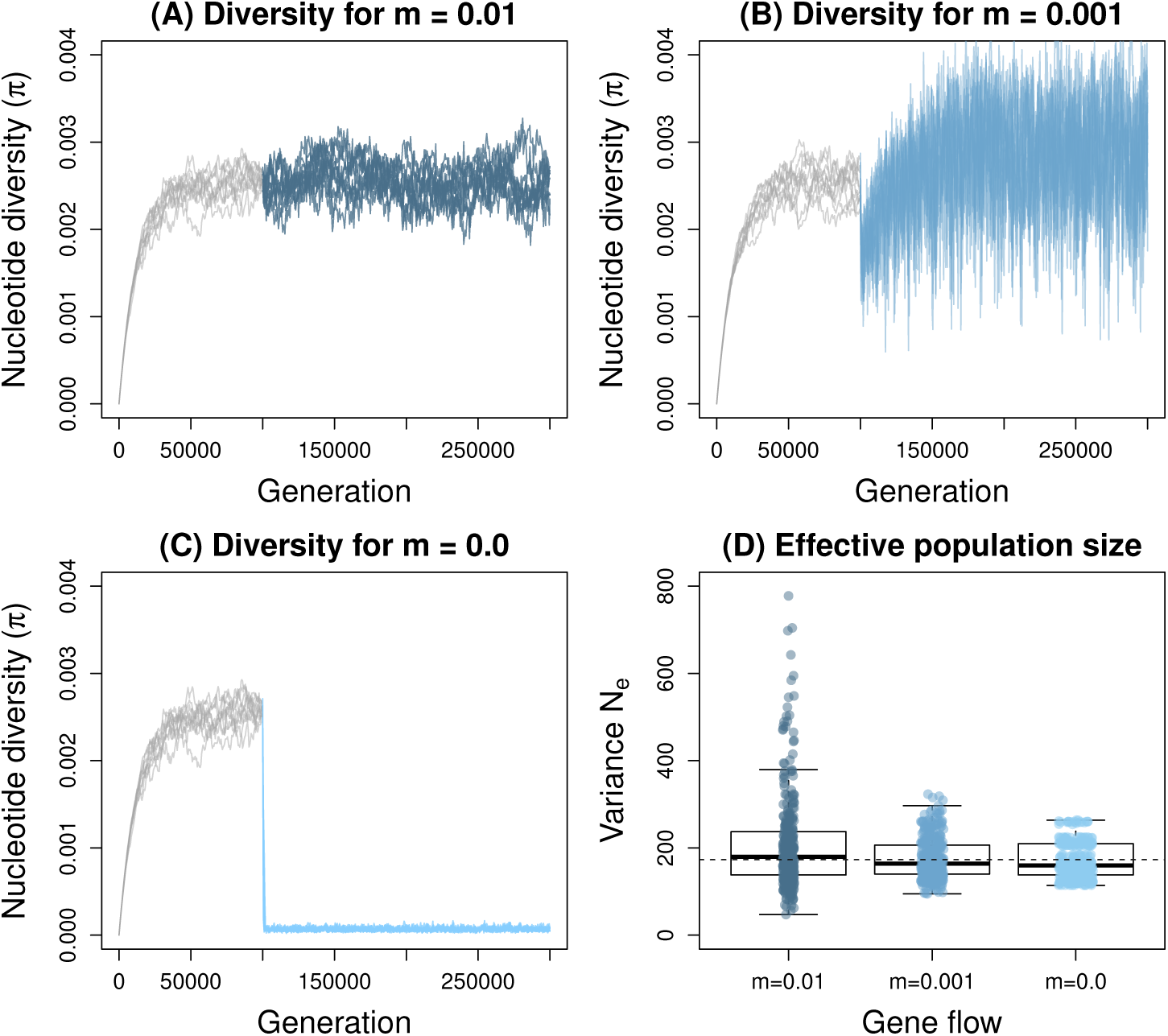
Simulations testing the effect of gene flow on variance *N_e_* (drift) and nucleotide diversity (*π*). Panels (A), (B), and (C) show nucleotide diversity over time across ten replicate simulation with *m* = 0.01 (high migration), 0.001 (low migration), and 0 (no migration), respectively. Colors indicate generations before (gray) versus after (blue) the single large panmictic population split into 36 populations with the same total size (a single non-edge population is shown after the split). Boxplots in panel (D) show estimates of variance effective population size (*N_e_*) based on patterns of allele frequency change between generations 150,000 and 150,002. Boxes denote the 1st and 3rd quartile with the median given by the midline; whiskers extend to the minimum and maximum value or 1.5× the interquartile range. Points show individual estimates for each replicate and population (a few extreme outliers, five with *N_e_ >* 800 and one negative estimate are not shown for the high migration scenario to facilitate visualization). Dashed lines correspond to *N_e_* = 173, which was per-deme population size used for simulations.

### Short-term effects of gene flow

In some cases, populations showed an excess of allele frequency change towards the average allele frequency 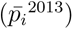 between 2013 and 2015, consistent with predictions from our contemporary effects of gene flow hypothesis (Fig. 7A). In particular, we detected an excess of change towards the mean allele frequency in BTB (*P* = 0.002) and USL (*P <* 0.001). In contrast, PSP, the population farthest to the south, showed an excess of change away from the mean (*P <* 0.001), and change for BCR, MRF, and RNV were consistent with null expectations of change in specific direction. Results were also mixed in terms of our second prediction of consistent change over time for HNV and SKI, with HNV showing a small but statistically significant excess of SNPs with consistent patterns of change (observed = 14.9%, null = 12.9%, *P <* 0.001), but SKI showing no such pattern (observed 11.2%, null

**Figure 7:**
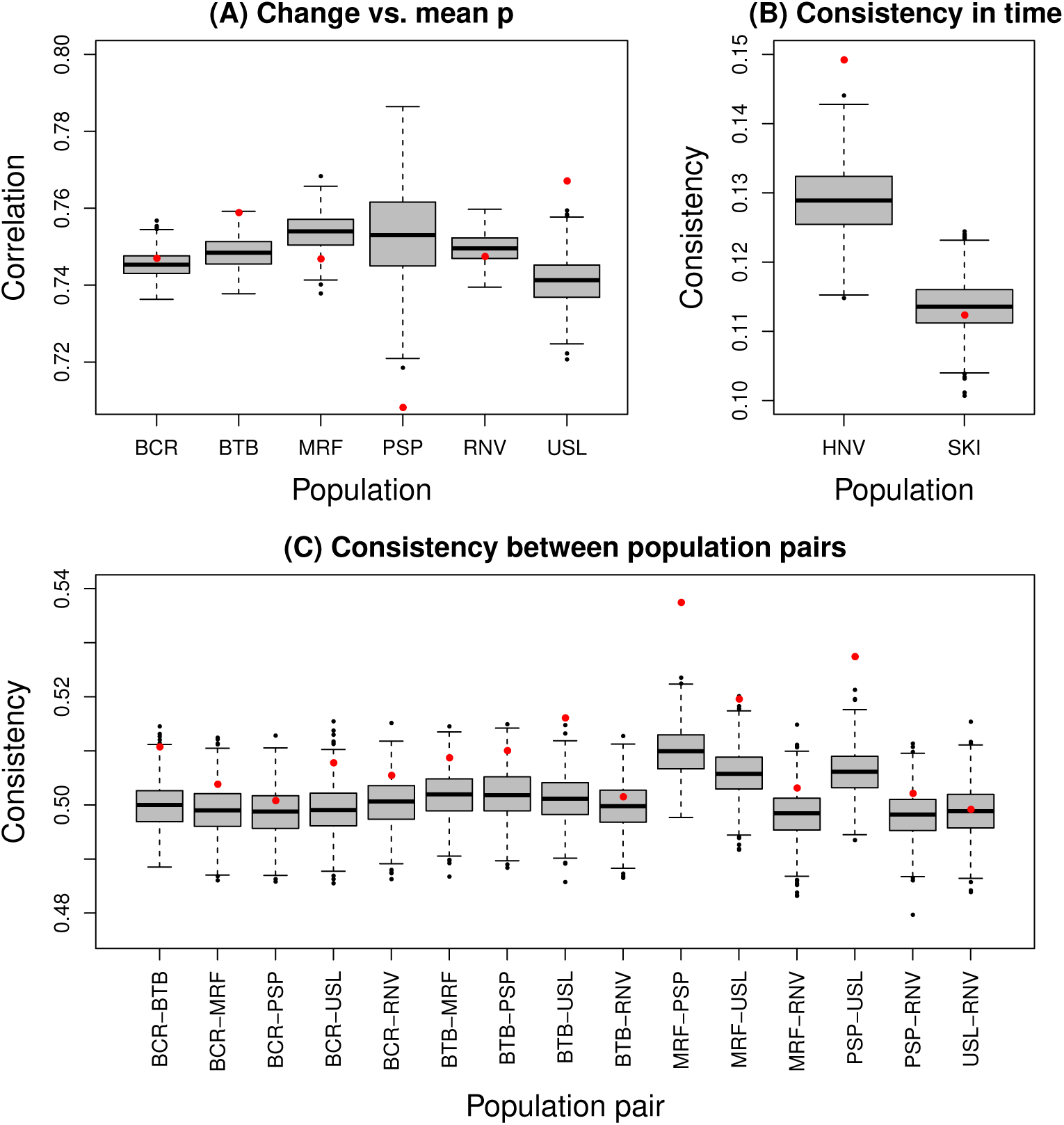
Evidence for contemporary evolutionary change caused by gene flow. In each panel, red circles denote observed values and boxplots show null distributions from 1000 permutations. Boxes denote the 1st and 3rd quartile with the median given by the midline; whiskers extend to the minimum and maximum value or 1.5× the interquartile range with points for more extreme values. Panel (A) shows the observed values and null expectations for correlations between the direction of the allele frequency difference between the mean and population for 2013 and the change from 2013 to 2015. Panel (B) shows the consistency in time (proportion of SNPs with the same direction of change across three time intervals) for change between in HNV and SKI. Panel (C) shows the consistency in change between 2013 and 2015 for pairs of populations (proportion of SNPs with change in the same direction for each pair of populations).

= 11.4%, *P* = 0.593; Fig. 7B). Lastly, we tested whether each of 15 pairs of populations showed an excess of consistency in terms of the direction of allele frequency change between 2013 and 2015 as predicted by our gene flow hypothesis. This was true for seven of the 15 pairs (all *P <* 0.05; Fig. 7C). Moreover, in 14 of the 15 cases change occurred in the same direction more than 50% of the time (51%, binomial probability, *P* = 0.0005) and the observed consistency exceeded the mean of the null in all 15 cases (binomial probability, *P* = 3×10^−5^).

## Discussion

Inferring evolutionary processes from patterns of genetic variation in space or time is a major goal in biology. However, making such connections is difficult because of the lack of a one-to-one mapping between genetic patterns and evolutionary processes (e.g., Wright, 1948). Nonetheless, by combining multiple sources of information, as we have done here, progress towards this goal can be made. Herein, we have shown that the maintenance of genetic diversity and rates of evolution by drift do not reflect the same evolutionary processes, at least not in a manner that can be captured by a single effective population size. Rates of evolution by drift were relatively high, with 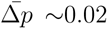 and 5% of SNPs showing change *>*0.05 over just a few generations. Such high rates were consistent with our estimates of small local (demic) variance effective population sizes and modest census sizes. In contrast, populations harbored substantial molecular diversity (i.e., *π* ∼0.009), which our analyses and simulations suggest have been maintained despite drift by periodic gene flow involving ∼1 to several immigrants per generation (year). This decoupling of diversity levels and short-term change by drift is relevant for our understanding of contemporary evolution and patterns of genetic variation in the wild, as we discuss in more detail below.

### Patterns and consequences of short-term evolutionary change

Our results show that non-trivial, genome-wide evolutionary change can occur on short time scales (5% of SNPs changing by *>* 0.05 and 1% changing by *>* 0.1). Similarly rapid genomic change has been documented in the lab (e.g., Turner *et al*., 2011; Gompert & Messina, 2016a; R^ego *et al*., 2019; Hardy *et al*., 2018; Langmüller & Schlötterer, 2020) and in field experiments (e.g., Barrett *et al*., 2008; Anderson *et al*., 2013; Gompert *et al*., 2014a; Egan *et al*., 2015; Marques *et al*., 2018; Exposito-Alonso *et al*., 2019). Population-genomic studies provide some evidence that high rates of genome-wide evolution might be less common in unmanipulated, natural populations (e.g., Pinsky *et al*., 2021). However, our results, combined with other recent studies of genomic time series (e.g., Campbell-Staton *et al*., 2017; Alves *et al*., 2019; Bi *et al*., 2019), show that appreciable genetic change can occur over short time periods in natural populations. Moreover, rates of inferred evolution often depend on the time-scale of measurements, because evolutionary fluctuations result in higher rates on shorter time scales (Hendry & Kinnison, 1999; Gingerich, 2019). Thus, fine temporal resolution, as was possible in our study, might provide better estimates of per-generation change than longer but sparser time series. Still, over slightly longer time periods, gene flow and drift may reverse much of the allele frequency change documented here making the observed change ephemeral. Indeed, our results suggest additional complexity in terms of predicting evolutionary change and patterns of adaptation. For example, local adaptation of demes might be impeded by low local *N_e_* (high rates of drift), but regional adaptation might be facilitated by higher metapopulation or species *N_e_* and the spread of adaptive alleles via gene flow. The balance of these processes might thus affect the geographic scale and extent of adaptation to heterogeneous environments.

Evolutionary change can have ecological consequences, even on short time scales (reviewed in Hendry, 2016). This is of particular interest given the potential for contemporary evolution and adaptation to reduce biodiversity loss caused by maladaptation to human-mediated global change (e.g., Mergeay & Santamaria, 2012; Mills *et al*., 2018; Oziolor *et al*., 2019; Capblancq *et al*., 2020). Our results provide additional evidence for rapid evolutionary change in natural populations, however, whether this change has ecological consequences in *L. idas* is unclear. We have thus far assumed that most of the change documented was caused by random genetic drift and (to a lesser extent) gene flow. However, selection likely contributed to the documented change, at least linked selection and especially for the SNPs with exceptionally high or consistent patterns of change (e.g., the 1% of SNPs with change *>* 10%). Moreover, polygenic selection (rather than gene flow) could explain the weak, genomewide patterns of consistent evolutionary change in time or space that we documented in some populations (see, e.g., Buffalo & Coop, 2019, 2020). Thus, as selection may have contributed to some of the observed change, it is reasonable to hypothesize that some of the change is of ecological relevance. Additional work involving increased temporal and genomic sampling is underway to better parse the contributions of selection and drift to the observed patterns of evolutionary change.

### Gene flow and diversity levels

Our results suggest that even in systems composed of populations occupying isolated habitat patches and organisms with limited dispersal abilities (Gompert *et al*., 2010), gene flow can have a small but detectable effect on patterns of change and a large effect on diversity levels. The potential for gene flow to be a potent evolutionary process, even when rare on an individual basis, has long been recognized. For example, even very low levels of gene flow can be sufficient for the spread of adaptive alleles among populations or species (Morjan & Rieseberg, 2004; Ellstrand, 2014), and theory suggests that any non-zero level of gene flow can maintain diversity levels (Whitlock & Barton, 1997). Our results are consistent with these theoretical predictions. For example, if we assume a mutation rate of 2.9×10^−9^ (from the butterfly *Heliconius melpomene*; Keightley *et al*., 2015) and *π* = 0.0092 (our mean estimate), diversity levels suggest a global, long-term effective population size of ∼793,000. This is ∼4500 times greater than the average variance effective population size we estimated, and thus would require ∼4500 populations of this size connected by gene flow. Given the geographic range of this species and the number of populations we have encountered in our own field work (hundreds), this is not entirely unreasonable.

This effect of gene flow on diversity levels might be relevant for resolving one of the great mysteries in evolutionary genetics, namely, Lewontin’s paradox of variation (Lewontin, 1974). Under neutral theory, molecular diversity levels are predicted to scale linearly with population size, at least at drift-mutation equilibrium (Kimura, 1983). However, studies of genetic variation in the wild show that diversity levels only vary by a few orders of magnitude across species, whereas population sizes vary by many more orders of magnitude (Lewontin, 1974; Leffler *et al*., 2012). Various solutions to this paradox have been proposed, most notably that linked selection constrains diversity levels (Gillespie, 2000; Corbett-Detig *et al*., 2015; Buffalo, 2021). However, none appear to be sufficient alone (Coop, 2016; Ellegren & Galtier, 2016; Buffalo, 2021). Another possibility is effective population size varies considerably less among species than does census population size. This could occur with boom-bust (“sweepstakes”) style reproduction (e.g., Nunney, 1993). Or, as one might surmise from our results, geographic isolation and gene flow could constrain long-term *N_e_* relative to global census sizes. In particular, if most species are broken up into numerous discrete populations or demes, there might be geographic limits to the number of populations that are connected by gene flow (i.e., beyond the spatial scale we studied), and thus to the effective population size relevant for diversity levels. This could be true even if rates of gene flow are non-zero, if migration rates are low enough to make the approach to drift-migration equilibrium very slow (as suggested by the high variance in our low migration simulations). The approach to equilibrium at large spatial scales may even be slow relative to the time-frame of speciation and other biogeographic events that result in severe bottlenecks (e.g., Wang *et al*., 2020). We think that future work focused on the combined effects of selection (including fluctuating selection) and the limits of gene flow could be critical for eventually resolving Lewontin’s paradox.

### Conclusions

In conclusion, by combining census population size estimates with spatial and temporal analyses of population genomic patterns, we showed that genome-wide change and the maintenance of diversity are driven largely by different processes, drift versus gene flow, and reflect dramatically different effective population sizes. This adds further complexity to arguments about the use of genetic diversity metrics in conservation biology (e.g., Reed & Frankham, 2003; Brüniche-Olsen *et al*., 2018; Wernberg *et al*., 2018; Scott *et al*., 2020; Teixeira & Huber, 2021). While most of the change documented was likely driven by drift, we showed patterns of change consistent with short-term effects of gene flow (and likely selection) as well. In general, we think our results highlight the potential utility of population-genomic time series data from natural populations for understanding evolutionary biology, especially the causes and consequences of contemporary evolution.

## Acknowledgments

We thank R. Olsen, P. Nelson, C. Gabbitas, and B. Allen for assistance with field work. We thank C. A. Buerkle, J. Fordyce, M. Forister, C. Nice, and P. Nosil for comments on an earlier draft of this manuscript. This research was funded by the National Science Foundation (DEB-1844941 to ZG) and the University of Wyoming NPS research station. The support and resources from the Center for High Performance Computing at the University of Utah are also gratefully acknowledged. The authors declare no conflicts of interest.

## Data Accessibility

The DNA sequence data analyzed in this manuscript have been archived on NCBI’s SRA (accession # pending). Computer scripts, census data, and key downstream data files, such as genotype and allele frequency estimates, will be archived on Dryad (DOI pending).

## Author Contributions

ZG and LKL conceived and designed the study. All authors collected the data. ZG analyzed the data. ZG drafted the initial version of the manuscript. All authors contributed to later versions of the manuscript.

## References

Alves JM, Carneiro M, Cheng JY, et al. (2019) Parallel adaptation of rabbit populations to myxoma virus. Science, 363, 1319–1326.

Anderson JT, Lee CR, Rushworth CA, Colautti RI, Mitchell-Olds T (2013) Genetic trade-offs and conditional neutrality contribute to local adaptation. Molecular Ecology, 22, 699–708.

Auckland JN, Debinski DM, Clark WR (2004) Survival, movement, and resource use of the butterfly parnassius clodius. Ecological Entomology, 29, 139–149.

Balding DJ, Nichols RA (1995) A method for quantifying differentiation between populations at multi-allelic loci and its implications for investigating identity and paternity. Genetica, 96, 3–12.

Barrett RDH, Rogers SM, Schluter D (2008) Natural selection on a major armor gene in threespine stickleback. Science, 322, 255–257.

Bergland AO, Behrman EL, O’Brien KR, Schmidt PS, Petrov DA (2014) Genomic evidence of rapid and stable adaptive oscillations over seasonal time scales in *Drosophila*. PLoS Genetics, 10, e1004775.

Bi K, Linderoth T, Singhal S, et al. (2019) Temporal genomic contrasts reveal rapid evolutionary responses in an alpine mammal during recent climate change. PLoS Genetics, 15, e1008119.

Brown GR, Gill GP, Kuntz RJ, Langley CH, Neale DB (2004) Nucleotide diversity and linkage disequilibrium in loblolly pine. Proceedings of National Academy of Sciences, 101, 15255–15260.

Brüniche-Olsen A, Austin JJ, Jones ME, Holland BR, Burridge CP (2016) Detecting selection on temporal and spatial scales: a genomic time-series assessment of selective responses to devil facial tumor disease. PloS ONE, 11, e0147875.

Brüniche-Olsen A, Kellner KF, Anderson CJ, DeWoody JA (2018) Runs of homozygosity have utility in mammalian conservation and evolutionary studies. Conservation Genetics, 19, 1295–1307.

Buckland ST, Anderson DR, Burnham KP, Laake JL, Borchers DL, Thomas L (2001) Introduction to distance sampling: estimating abundance of biological populations. Oxford University Press, USA.

Buffalo V (2021) Why do species get a thin slice of *π*? revisiting lewontin’s paradox of variation. *bioRxiv* .

Buffalo V, Coop G (2019) The linked selection signature of rapid adaptation in temporal genomic data. Genetics, 213, 1007–1045.

Buffalo V, Coop G (2020) Estimating the genome-wide contribution of selection to temporal allele frequency change. Proceedings of the National Academy of Sciences, 117, 20672– 20680.

Burke MK, Dunham JP, Shahrestani P, Thornton KR, Rose MR, Long AD (2010) Genomewide analysis of a long-term evolution experiment with *Drosophila*. Nature, 467, 587–590.

Campbell-Staton SC, Cheviron ZA, Rochette N, Catchen J, Losos JB, Edwards SV (2017) Winter storms drive rapid phenotypic, regulatory, and genomic shifts in the green anole lizard. Science, 357, 495–498.

Capblancq T, Fitzpatrick MC, Bay RA, Exposito-Alonso M, Keller SR (2020) Genomic prediction of (mal) adaptation across current and future climatic landscapes. Annual Review of Ecology, Evolution, and Systematics, 51, 245–269.

Charlesworth B (2009) Effective population size and patterns of molecular evolution and variation. Nature Reviews Genetics, 10, 95–205.

Chaturvedi S, Lucas LK, Buerkle CA, et al. (2020) Recent hybrids recapitulate ancient hybrid outcomes. Nature Communications, 11, 1–15.

Chaturvedi S, Lucas LK, Nice CC, Fordyce JA, Forister ML, Gompert Z (2018) The predictability of genomic changes underlying a recent host shift in Melissa blue butterflies. Molecular Ecology, 27, 2651–2666.

Coop G (2016) Does linked selection explain the narrow range of genetic diversity across species? BioRxiv, p. 042598.

Corbett-Detig RB, Hartl DL, Sackton TB (2015) Natural selection constrains neutral diversity across a wide range of species. PLoS Biol, 13, e1002112.

Crow JF, Denniston C (1988) Inbreeding and variance effective population numbers. Evolution, 42, 482–495.

Do C, Waples RS, Peel D, Macbeth G, Tillett BJ, Ovenden JR (2014) NeEstimator v2: re-implementation of software for the estimation of contemporary effective population size (Ne) from genetic data. Molecular Ecology Resources, 14, 209–214.

Driscoe AL, Nice CC, Busbee RW, Hood GR, Egan SP, Ott JR (2019) Host plant associations and geography interact to shape diversification in a specialist insect herbivore. Molecular Ecology, 28, 4197–4211.

Egan SP, Ragland GJ, Assour L, et al. (2015) Experimental evidence of genome-wide impact of ecological selection during early stages of speciation-with-gene-flow. Ecology Letters, 18, 817–825.

Ellegren H, Galtier N (2016) Determinants of genetic diversity. Nature Reviews Genetics, 17, 422–433.

Ellstrand NC (2014) Is gene flow the most important evolutionary force in plants? American Journal of Botany, 101, 737–753.

Exposito-Alonso M, Burbano HA, Bossdorf O, Nielsen R, Weigel D (2019) Natural selection on the *Arabidopsis thaliana* genome in present and future climates. Nature, 573, 126–129.

Falush D, Stephens M, Pritchard JK (2003) Inference of population structure using multilocus genotype data: linked loci and correlated allele frequencies. Genetics, 164, 1567–1587.

Feng C, Pettersson M, Lamichhaney S, et al. (2017) Moderate nucleotide diversity in the atlantic herring is associated with a low mutation rate. eLife, 6, e23907.

Fisher RA, Ford EB (1947) The spread of a gene in natural conditions in a colony of the moth *Panaxia dominula* L. Heredity, 1, 143–174.

Fiske I, Chandler R (2011) unmarked: An R package for fitting hierarchical models of wildlife occurrence and abundance. Journal of Statistical Software, 43, 1–23.

Foll M, Shim H, Jensen JD (2015) WFABC: a Wright–Fisher ABC-based approach for inferring effective population sizes and selection coefficients from time-sampled data. Molecular Ecology Resources, 15, 87–98.

Ford EB (1977) Ecological Genetics. Springer.

Frankham R (1995) Effective population size/adult population size ratios in wildlife: a review. Genetics Research, 66, 95–107.

Gaggiotti OE, Foll M (2010) Quantifying population structure using the F-model. Molecular Ecology Resources, 10, 821–830.

Gilbert KJ, Whitlock MC (2015) Evaluating methods for estimating local effective population size with and without migration. Evolution, 69, 2154–2166.

Gillespie JH (2000) Genetic drift in an infinite population: The pseudohitchhiking model. Genetics, 155, 909–919.

Gingerich PD (2019) Rates of Evolution: A Quantitative Synthesis. Cambridge University Press.

Gompert Z (2016) Bayesian inference of selection in a heterogeneous environment from genetic time-series data. Molecular Ecology, 25, 121–134.

Gompert Z, Comeault AA, Farkas TE, et al. (2014a) Experimental evidence for ecological selection on genome variation in the wild. Ecology Letters, 17, 369–379.

Gompert Z, Fordyce JA, Forister ML, Shapiro AM, Nice CC (2006) Homoploid hybrid speciation in an extreme habitat. Science, 314, 1923–1925.

Gompert Z, Lucas LK, Buerkle CA, Forister ML, Fordyce JA, Nice CC (2014b) Admixture and the organization of genetic diversity in a butterfly species complex revealed through common and rare genetic variants. Molecular Ecology, 23, 4555–4573.

Gompert Z, Lucas LK, Fordyce JA, Forister ML, Nice CC (2010) Secondary contact between *Lycaeides idas* and *L. melissa* in the Rocky Mountains: extensive introgression and a patchy hybrid zone. Molecular Ecology, 19, 3171–3192.

Gompert Z, Lucas LK, Nice CC, Fordyce JA, Buerkle CA, Forister ML (2013) Geographically multifarious phenotypic divergence during speciation. Ecology and Evolution, 3, 595–613.

Gompert Z, Lucas LK, Nice CC, Fordyce JA, Forister ML, Buerkle CA (2012) Genomic regions with a history of divergent selection affect fitness of hybrids between two butterfly species. Evolution, 66, 2167–2181.

Gompert Z, Messina FJ (2016a) Genomic evidence that resource-based trade-offs limit host-range expansion in a seed beetle. Evolution, 70, 1249–1264.

Gompert Z, Messina FJ (2016b) Genomic evidence that resource-based trade-offs limit host-range expansion in a seed beetle, Dryad, Dataset. https://doi.org/10.5061/dryad. 1p0rm.

Graves Jr J, Hertweck K, Phillips M, et al. (2017) Genomics of parallel experimental evolution in *Drosophila*. Molecular Biology and Evolution, 34, 831–842.

Green RE, Krause J, Briggs AW, et al. (2010) A draft sequence of the Neandertal genome. Science, 328, 710–722.

Gutenkunst RN, Hernandez RD, Williamson SH, Bustamante CD (2009) Inferring the joint demographic history of multiple populations from multidimensional SNP frequency data. PLoS Genetics, 5, e1000695.

Haller BC, Messer PW (2019) SLiM 3: forward genetic simulations beyond the wright–fisher model. Molecular Biology and Evolution, 36, 632–637.

Hardy CM, Burke MK, Everett LJ, Han MV, Lantz KM, Gibbs AG (2018) Genome-wide analysis of starvation-selected *Drosophila melanogaster* —a genetic model of obesity. Molecular Biology and Evolution, 35, 50–65.

Harris RB, Sackman A, Jensen JD (2018) On the unfounded enthusiasm for soft selective sweeps II: Examining recent evidence from humans, flies, and viruses. PLoS Genetics, 14, e1007859.

Hendry AP (2016) Eco-evolutionary dynamics. Princeton University Press.

Hendry AP, Kinnison MT (1999) Perspective: the pace of modern life: measuring rates of contemporary microevolution. Evolution, pp. 1637–1653.

Hey J, Nielsen R (2004) Multilocus methods for estimating population sizes, migration rates and divergence time, with applications to the divergence of drosophila pseudoobscura and d. persimilis. Genetics, 167, 747–760.

Hill WG (1981) Estimation of effective population size from data on linkage disequilibrium. Genetics Research, 38, 209–216.

Jorde PE, Ryman N (1995) Temporal allele frequency change and estimation of effective size in populations with overlapping generations. Genetics, 139, 1077–1090.

Jorde PE, Ryman N (2007) Unbiased estimator for genetic drift and effective population size. Genetics, 177, 927–935.

Keightley PD, Pinharanda A, Ness RW, et al. (2015) Estimation of the spontaneous mutation rate in *Heliconius melpomene*. Molecular Biology and Evolution, 32, 239–243.

Kettlewell HD (1958) A survey of the frequencies of *Biston betularia* (L.)(Lep.) and its melanic forms in great britain. Heredity, 12, 51–72.

Kimura M (1983) The Neutral Theory of Molecular Evolution. Cambridge University Press.

Knutson RL, Kwilosz JR, Grundel R (1999) Movement patterns and population characteristics of the Karner blue butterfly (*Lycaeides melissa samuelis*) at Indiana Dunes National Lakeshore. Natural Areas Journal, 19, 109–120.

Korneliussen TS, Albrechtsen A, Nielsen R (2014) ANGSD: analysis of next generation sequencing data. BMC Bioinformatics, 15, 1–13.

Krimbas CB, Tsakas S (1971) The genetics of *Dacus oleae*. V. Changes of esterase polymorphism in a natural population following insecticide control-selection or drift? Evolution, pp. 454–460.

Kruuk LEB, Baird SJE, Gale KS, Barton NH (1999) A comparison of multilocus clines maintained by environmental adaptation or by selection against hybrids. Genetics, 153, 1959–1971.

Lande R (1992) Neutral theory of quantitative genetic variance in an island model with local extinction and colonization. Evolution, 46, 381–389.

Lanfear R, Kokko H, Eyre-Walker A (2014) Population size and the rate of evolution. Trends in Ecology & Evolution, 29, 33–41.

Langmüller AM, Schlötterer C (2020) Low concordance of short-term and long-term selection responses in experimental *Drosophila* populations. Molecular ecology, 29, 3466–3475.

Lawson DJ, Van Dorp L, Falush D (2018) A tutorial on how not to over-interpret STRUCTURE and ADMIXTURE bar plots. Nature Communications, 9, 1–11.

Leffler EM, Bullaughey K, Matute DR, et al. (2012) Revisiting an old riddle: What determines genetic diversity levels within species? PLoS Biology, 10, e1001388.

Lewontin R (1974) The genetic basis of evolutionary change. Columbia University Press, New York, NY, USA.

Li H (2011) A statistical framework for SNP calling, mutation discovery, association mapping and population genetical parameter estimation from sequencing data. Bioinformatics, 27, 2987–2993.

Li H, Durbin R (2009) Fast and accurate short read alignment with burrows–wheeler transform. Bioinformatics, 25, 1754–1760.

Li H, Durbin R (2011) Inference of human population history from individual whole-genome sequences. Nature, 475, 493–496.

Li H, Handsaker B, Wysoker A, et al. (2009) The Sequence Alignment/Map format and SAMtools. Bioinformatics, 25, 2078–2079.

Marques DA, Jones FC, Di Palma F, Kingsley DM, Reimchen TE (2018) Experimental evidence for rapid genomic adaptation to a new niche in an adaptive radiation. Nature Ecology & Evolution, 2, 1128–1138.

McKenna A, Hanna M, Banks E, et al. (2010) The Genome Analysis Toolkit: A MapReduce framework for analyzing next-generation DNA sequencing data. Genome Research, 20, 1297–1303.

Mergeay J, Santamaria L (2012) Evolution and biodiversity: the evolutionary basis of biodiversity and its potential for adaptation to global change. Evolutionary Applications, 5, 103.

Messer PW, Ellner SP, Hairston NG (2016) Can population genetics adapt to rapid evolution? Trends in Genetics, 32, 408–418.

Mills LS, Bragina EV, Kumar AV, et al. (2018) Winter color polymorphisms identify global hot spots for evolutionary rescue from climate change. Science, 359, 1033–1036.

Morjan CL, Rieseberg LH (2004) How species evolve collectively: implications of gene flow and selection for the spread of advantageous alleles. Molecular Ecology, 13, 1341–1356.

Mueller LD, Barr LG, Ayala FJ (1985) Natural selection vs. random drift: Evidence from temporal variation in allele frequencies in nature. Genetics, 111, 517–554.

Nabokov V (1943) The nearctic forms of *Lycaeides* Hub. (Lycaenidae, Lepidoptera). Psyche, 50, 87–99.

Nabokov V (1949) The nearctic members of *Lycaeides* Hübner (Lycaenidae, Lepidoptera). Bulletin of the Museum of Comparative Zoology, 101, 479–541.

Nei M, Graur D (1984) Extent of protein polymosphism and the neutral mutation theory. Evolutionary Biology, 17, 73–118.

Nei M, Tajima F (1981) Genetic drift and estimation of effective population size. Genetics, 98, 625–640.

Nei M, Takahata N (1993) Effective population size, genetic diversity, and coalescence time in subdivided populations. Journal of Molecular Evolution, 37, 240–244.

Nicholson G, Smith AV, Jonsson F, Gustafsson O, Stefansson K, Donnelly P (2002) Assessing population differentiation and isolation from single-nucleotide polymorphism data. Journal of the Royal Statistical Society Series B-Methodological, 64, 695–715.

Nielsen R, Mountain JL, Huelsenbeck JP, Slatkin M (1998) Maximum-likelihood estimation of population divergence times and population phylogeny in models without mutation. Evolution, 52, 669–677.

Nordborg M, Krone S (2002) Separation of time scales and convergence to the coalescent in structured populations. In: Modern developments in theoretical population genetics (eds. Slatkin M, Veuille M), pp. 194–232.

Nunney L (1993) The influence of mating system and overlapping generations on effective population size. Evolution, 47, 1329–1341.

Nunziata SO, Weisrock DW (2018) Estimation of contemporary effective population size and population declines using RAD sequence data. Heredity, 120, 196–207.

Oksanen J, Blanchet FG, Friendly M, et al. (2020) vegan: Community Ecology Package. R package version 2.5–7.

Oziolor EM, Reid NM, Yair S, et al. (2019) Adaptive introgression enables evolutionary rescue from extreme environmental pollution. Science, 364, 455–457.

Pääbo S, Poinar H, Serre D, et al. (2004) Genetic analyses from ancient DNA. Annual Review of Genetics, 38, 645–679.

Palstra FP, Ruzzante DE (2008) Genetic estimates of contemporary effective population size: what can they tell us about the importance of genetic stochasticity for wild population persistence? Molecular Ecology, 17, 3428–3447.

Parchman TL, Gompert Z, Mudge J, Schilkey F, Benkman CW, Buerkle CA (2012) Genomewide association genetics of an adaptive trait in lodgepole pine. Molecular Ecology, 21, 2991–3005.

Pazminõ DA, Maes GE, Simpfendorfer CA, Salinas-de León P, van Herwerden L (2017) Genome-wide snps reveal low effective population size within confined management units of the highly vagile galapagos shark (carcharhinus galapagensis). Conservation Genetics, 18, 1151–1163.

Pinsky ML, Eikeset AM, Helmerson C, et al. (2021) Genomic stability through time despite decades of exploitation in cod on both sides of the atlantic. Proceedings of the National Academy of Sciences, 118.

Rannala B, Hartigan JA (1996) Estimating gene flow in island populations. Genetical Research, 67, 147–158.

Reed DH, Frankham R (2003) Correlation between fitness and genetic diversity. Conservation Biology, 17, 230–237.

Rêgo A, Messina FJ, Gompert Z (2019) Dynamics of genomic change during evolutionary rescue in the seed beetle *Callosobruchus maculatus*. Molecular Ecology, 28, 2136–2154.

Royle JA, Dawson DK, Bates S (2004) Modeling abundance effects in distance sampling. Ecology, 85, 1591–1597.

Ryan SF, Deines JM, Scriber JM, et al. (2018) Climate-mediated hybrid zone movement revealed with genomics, museum collection, and simulation modeling. Proceedings of the National Academy of Sciences, 115, E2284–E2291.

Ryman N, Laikre L, Hössjer O (2019) Do estimates of contemporary effective population size tell us what we want to know? Molecular Ecology, 28, 1904–1918.

Scott J (1986) The Butterflies of North America: A Natural History and Field Guide. Stanford University Press.

Scott PA, Allison LJ, Field KJ, Averill-Murray RC, Shaffer HB (2020) Individual heterozygosity predicts translocation success in threatened desert tortoises. Science, 370, 1086– 1089.

Serbezov D, Jorde PE, Bernatchez L, Olsen EM, Vøllestad LA (2012) Short-term genetic changes: evaluating effective population size estimates in a comprehensively described brown trout (*Salmo trutta*) population. Genetics, 191, 579–592.

Shastry V, Adams PE, Lindtke D, et al. (2021) Model-based genotype and ancestry estimation for potential hybrids with mixed-ploidy. Molecular Ecology Resources, n/a.

Sjodin P, Kaj I, Krone S, Lascoux M, Nordborg M (2005) On the meaning and existence of an effective population size. Genetics, 169, 1061–1070.

Slatkin M (1985) Rare alleles as indicators of gene flow. Evolution, 39, pp. 53–65.

Slatkin M, Racimo F (2016) Ancient DNA and human history. Proceedings of the National Academy of Sciences, 113, 6380–6387.

Soria-Carrasco V, Gompert Z, Comeault AA, et al. (2014) Stick insect genomes reveal natural selection’s role in parallel speciation. Science, 344, 738–742.

Sousa VC, Grelaud A, Hey J (2011) On the non-identifiability of migration time estimates in isolation with migration models. Molecular Ecology, 20, 3956.

Speidel L, Forest M, Shi S, Myers SR (2019) A method for genome-wide genealogy estimation for thousands of samples. Nature Genetics, 51, 1321–1329.

Stan Development Team (2019) RStan: the R interface to Stan. R package version 2.19.2. Stan Development Team (2021) Stan Modeling Language Users Guide and Reference Manual .

Stern AJ, Speidel L, Zaitlen NA, Nielsen R (2021) Disentangling selection on genetically correlated polygenic traits via whole-genome genealogies. The American Journal of Human Genetics, 108, 219–239.

Teixeira JC, Huber CD (2021) The inflated significance of neutral genetic diversity in conservation genetics. Proceedings of the National Academy of Sciences, 118.

Turner TL, Stewart AD, Fields AT, Rice WR, Tarone AM (2011) Population-based resequencing of experimentally evolved populations reveals the genetic basis of body size variation in drosophila melanogaster. PLoS Genetics, 7, e1001336.

U.S. Fish and Wildlife Service (2003) Karner Blue Butterfly (Lycaeides melissa samuelis) Recovery Plan. Tech. rep., Region 3, U.S. Fish and Wildlife Service, Fort Snelling, Min-nesota.

Wakeley J, Sargsyan O (2009) Extensions of the coalescent effective population size. Genetics, 181, 341–345.

Walsh B, Lynch M (2018) Evolution and Selection of Quantitative Traits. Oxford University Press.

Wang J, Santiago E, Caballero A (2016) Prediction and estimation of effective population size. Heredity, 117, 193–206.

Wang J, Whitlock MC (2003) Estimating effective population size and migration rates from genetic samples over space and time. Genetics, 163, 429–446.

Wang X, Bernhardsson C, Ingvarsson PK (2020) Demography and natural selection have shaped genetic variation in the widely distributed conifer Norway spruce (*Picea abies*). Genome Biology and Evolution, 12, 3803–3817.

Waples RS (1989) A generalized approach for estimating effective population size from temporal changes in allele frequency. Genetics, 121, 379–391.

Wernberg T, Coleman MA, Bennett S, Thomsen MS, Tuya F, Kelaher BP (2018) Genetic diversity and kelp forest vulnerability to climatic stress. Scientific Reports, 8, 1–8.

Whitlock M, McCauley D (1999) Indirect measures of gene flow and migration: F-ST not equal 1/(4Nm+1). Heredity, 82, 117–125.

Whitlock MC, Barton NH (1997) The effective size of a subdivided population. Genetics, 146, 427–441.

Wright S (1931) Evolution in Mendelian populations. Genetics, 16, 0097–0159.

Wright S (1948) On the roles of directed and random changes in gene frequency in the genetics of populations. Evolution, pp. 279–294.

Yang M, He Z, Shi S, Wu CI (2017) Can genomic data alone tell us whether speciation happened with gene flow?

