## Supplementary Material for "Genomic time-series data show that gene flow maintains high genetic diversity despite substantial genetic drift in a butterfly species"

### Supplemental Text

#### Preparing the GBS libraries

We created reduced complexity, double-digest restriction fragment-based DNA libraries for each of 1536 *Lycaeides* butterflies using laboratory methods initially described by Gompert *et al.* (2012) and Parchman *et al.* (2012), with modifications introduced by Gompert *et al.* (2014b). This is a genotyping-by-sequencing (GBS) approach. Specifically, double-stranded DNA oligos with sequences overlapping the enzyme-cut sites and including a 8-100bp sequence barcode and the Illumina adaptor sequences (1 $\mu$L each at 10 $\mu$M) were then ligated to the fragmented DNA with T4 DNA ligase (0.17 $\mu$L at 20,000 units per mL). We then PCR-amplified the fragment libraries with standard Illumina PCR primers (30 cycles with annealing for 30 s at 60 °C, and extension for 30 s at 72°C followed by a final amplification with 2 min for annealing and 10 min for extension). Amplified libraries were then pooled, purified, and size-selected (300-450 bp) with a BluePippin at the Utah State University genomics core lab.

#### Variant calling and filtering

We called SNPs separately using both `samtools` (version 1.5) combined with `bcftools` (version 1.6) and `GAKT`'s `HaplotypeCaller` and the `GenotypeGVCFs` module (version 3.5) (McKenna *et al.*, 2010). Variant calling with `samtools` involved the `samtools mpileup` and `bcftools call` functions. Specifically, we used the recommended mapping-quality adjustment for Illumina data (`-C` 50), skipped alignments with mapping quality <20, skipped bases with base quality <30, and ignored insertion–deletion polymorphisms. We set the prior on SNPs to 0.001 (`-P`) and called SNPs when the posterior probability that the nucleotide was invariant was $\leq$0.01 (`-p`). For `GATK`, we first used the `HaplotypeCaller` to generate gvcf files for each butterfly. We set the prior expectation for heterozygosity to 0.001, skipped alignments with mapping qualities less than 30, and applied the "AGRESSIVE" PCR error model. Joint variant calling was then accomplished using the `GenotypeGVCFs` module.

We then filtered each set of SNPs to retain only those with data for at least 80% of the butterflies, a mean coverage per butterfly of at least 2×, $\geq$20 reads supporting the non-reference allele, no more than one alternative allele (i.e., we retained only bi-allelic SNPs), a minimum (overall) minor allele frequency of at least 0.005, and no more than 1% of the reads in the reverse orientation (this is an expectation for our GBS method). For the `samtools` SNP data set, we additionally removed SNPs failing the base quality rank sum test, mapping quality rank sum test, or read position rank sum test with $P < 0.01$. Similarly, for the `GATK` SNP set, with corresponding test statistics output by `GATK` greater than 3 (base quality rank sum test), 2.5 (mapping quality rank sum test), or 2 (read position rank sum test). With the `GATK` SNP set, we also excluded SNPs with a ratio of variant confidence to non-reference read depth less than 2. These minor differences in filtering between the two SNP sets result from differences in the variant quality information calculated by the different variant callers. We further removed SNPs with excessive coverage (3 standard deviations above the mean) or that were tightly clustered (within 3 bp of each other), as these could reflect poor alignments (e.g., reads from multiple paralogs mapping to the same region of the genome).

Next, to reduce bias associated with batch effects (see Table 1), we removed SNPs with substantial differences in sequence coverage between the two sets. Specifically, we dropped any SNP with a difference in sequence coverage between the two sequencing batches that was more than half the mean coverage for the two data sets combined. Lastly, to minimize inclusion of spurious SNP calls, we retained only the set of SNPs that were called by both variant callers and passed all of these filters in each variant caller SNP set. This left us with 12,886 SNPs for downstream analysis.

#### Estimating census population sizes

We used a distance sampling protocol to estimate *L. idas* adult population sizes at each of our focal sites. Distance sampling involves counting individuals and recording their distance from a transect line or point (Buckland *et al.*, 2001). This distance information is used to estimate a detection function that accounts for imperfect detection away from the transect line. For each population we randomly chose ten or fewer random points within a defined area of suitable habitat (we identified suitable habitat from ground surveys and satellite images). At each of these points, we walked an approximately 100-meter transect and: 1) counted the *L. idas* we saw along the way, recorded the sex and measured their distance on and from the transect line, and 2) quantified the abundance of butterfly host plant. We recorded a 0, 1 or 2 to denote whether there were no butterfly host-plants, less than 50% of the ground cover was host-plants, or more than 50% of the ground cover was host-plants within a meter of each transect line, respectively. The host-plant species recorded depended on the population: *Astragalus miser* (BCR, BTB, MRF, HNV, BNP, GNP, SKI, USL), *Astragalus bisulcatus* (USL), *Lupinus sp.* (PSP), and *Hedysarum sp.* (RNV, SKI). We only performed distance sampling between 10:00 am and 3:00 pm under sunny or partly sunny skies. Our aim was to obtain estimates during the peak of the adult flight. We missed the peak flight for some populations in some years (i.e., only or mostly males flying or only or mostly older, worn butterflies), and we either returned for a second count (which was used) or did not use the data.

On July 22, 2018, we obtained a mark-release-recapture estimate of population size at BTB to compare with the estimate from the distance sampling method (conducted July 18, 2018). We captured 64 adult *L. idas* on July 18. We made a mark on the hind-wing with a permanent marker, and then released each butterfly (as in Auckland *et al.*, 2004). We returned the following day and captured 50 adults to check for markings. This short time between release and recapture minimizes birth, mortality and movement of butterflies into and out of the site. We then obtained a Bayesian estimate of the census population size by fitting a beta-binomial model for the number of recaptures with the shape parameters $\alpha$ and $\beta$ of the beta prior set to 0 (this is an improper prior but yields a proper posterior). The posterior is a known beta distribution, and was thus solved for analytically. With three out of 50 recaptures on the second day, our estimate of the census size on July 18th was 1187.3 (95% equal-tail probability intervals = 462.0–5075.7), and thus our estimate of the total adult flight was $3 \times 1187.3 = 3561.9$ (95% ETPI = 1386.0–15227.1). This is larger than the corresponding distance-sampling estimate ($N = 1884.8$), but the ETPI of this estiamte overlaps with the distance-sampling estimate.

#### Demographic inference with $\delta$a$\delta$i

We fit isolation with migration (IM) models for each pair of populations using $\delta$a$\delta$i. Specifically, we estimated an ancestral effective population size ($\theta = 4N_{ref}\mu$), split proportion ($s$), split time ($T_{split} = 2N_{ref}$ generations), population growth parameters ($\nu_1$ and $\nu_2$, size of each population relative to the ancestral population), and migration rates $M_{12}$ and $M_{21}$ (number of migrants assuming the ancestral effective population size, $N_{ref}$) based on the join site frequency spectrum for each pair of populations (see Fig. S1). We based our inferences only on the 2017 samples (which were processed in a single batch for all populations) and only used the `samtools` method for calculating genotype likelihoods. The `python` specification of the IM model was as follows:

```
def IM(params, ns, pts):
        s, nu1, nu2, T, m12, m21 = params
        xx = Numerics.default_grid(pts)
        phi = PhiManip.phi_1D(xx)
        phi = PhiManip.phi_1D_to_2D(xx, phi)
        nu1_func = lambda t : s * (nu1/s) ** (t/T)
        nu2_func = lambda t : (1-s) * (nu2/(1-s)) ** (t/T)
        phi = Integration.two_pops(phi, xx, T, nu1_func,
              nu2_func, m12 = m12, m21 = m21)
        fs = Spectrum.from_phi(phi, ns, (xx, xx))
        return fs
```

Grid points for extrapolation were set to 60, 70 and 80. We set the following bounds on the model parameters: 0.1 to 0.9 for $s$, 0.1 to 20 for $\nu_1$ and $\nu_2$, 0 to 4 for $T_{split}$, and 0 to 10 for $M_{12}$ and $M_{21}$. We obtained initial values by applying a small perturbation to $s = 0.5$, $\nu_1 = 1$, $\nu_2 = 1.5$, $T_{split} = 0.004$, $M_{12} = 0.4$, and $M_{21} = 0.4$. We set $\theta$ to the optimal value conditional on the other model parameters. We then repeatedly optimized the model using $\delta$a$\delta$i. Fifty iterations were performed for each optimization. We then perturbed the optimal solution and re-optimized the model. We repeated this procedure 15 times to obtain the maximum likelihood estimate of the parameters (the most likely solution across all 15 optimizations). We defined overall convergence of the algorithm as a coefficient of variation for the five best fits of less than 0.01 (in terms of log-likelihoods) and a standard deviation for the number of migrants across these same best fits of less than 1. We obtained convergence for 35 of the population pairs (78%), and our inferences are based on this. Estimates of migration were converted to number of migrants per generation by multiplying each inferred migration rate by the relevant population growth parameter and dividing by two, e.g., $Nm_{12} = \nu_1 M_{12}/2$.

### Quantitative basis for assuming drift-migration equilibrium

Our second approach to inferring gene flow requires assuming an equilibrium has been reached between the opposing processes of drift and gene flow, or similarly, that the historical effects of the divergence from a single ancestral source population at some point in the past have been erased (Wright, 1931; Slatkin, 1985; Whitlock & McCauley, 1999). This can be problematic, but that is not likely the case here, at least not to such an extent to introduce substantial errors. And a benefit of this approach is that it allows us to treat many populations in a single, computationally efficient analysis. Our reasoning is as follows.

First, the Bayesian F-model we fit can be parameterized to correspond with demographic models of an island model at drift-migration equilibrium or a non-equilibrium model of divergence from a common ancestor with no gene flow; these two choices simply result in different parameterizations of the same model (or equivalently different interpretations of the parameters) (Nielsen *et al.*, 1998; Falush *et al.*, 2003; Gaggiotti & Foll, 2010). Thus, we can use the same model to calculate divergence times that would give rise to the observed levels of genetic differentiation in the absence of gene flow. In particular, if we use our average estimate of contemporary $N_e$ from the time-series analysis of 173, we obtain an estimate of $\sim$11 generations. That is, about 11 generations would be required in the absence of gene flow to explain levels of differentiation seen in the data set. This is unlikely. We have been visiting these populations for more than 11 years (11 generations) and thus can be certain that they did not diverge from a common ancestral population only 11 generations ago. Moreover, many of these populations were sampled by others in the mid 1900s, further pushing back the possible time at which they could have descended from a common ancestor (Nabokov, 1943, 1949). Thus, some level of gene flow is required to explain the low levels of genetic differentiation observed in this system.

Second, if we assume the inferred level of gene flow is correct, we can determine how long the system would take to reach drift-migration equilibrium, and thus assess whether, at least approximately, such an equilibrium is likely to hold. Specifically, by iteratively calculating $F'_{ST} = \left[\frac{1}{2Ne} + \left(1 - \frac{1}{2Ne}\right) F_{ST}\right] (1-m)^2$ with $N_e = 173$ and $m = \frac{N_e}{Nm} = \frac{173}{7.6} = 0.044$, and starting from $F_{ST} = 0$, we can show that it takes only $\sim$40 generations (i.e., 40 years) to approach drift-migration equilibrium (Fig. S4). This is a relatively short amount of time, and there is no evidence for major changes in the distribution of *L. idas* in the geographic region of interest during this period. Thus, strict isolation with no gene flow is very unlikely, and the system is likely not far from drift-migration equilibrium.

### Simulations with SLiM3

As described in the main text, we simulated evolution forward in time under a Wright-Fisher model with `SLiM3` (Haller & Messer, 2019). We considered a spatial matrix of 36 populations arranged in a 6×6 grid, each with a variance $N_e$ of 173 (the mean for *L. idas*). We assumed that these were descended from a single large, panmictic population of 6228 individuals (i.e., 173×36). Migration occurred between neighboring demes with $m = 0.001$ (low migration) or $m = 0.01$ (high migration); we also simulated the case of no migration ($m = 0$). Our high migration rate ($m = 0.01$) corresponds approximately with our migration rate estimate from the Bayesian F-model, and we consider the low migration rate ($m = 0.001$) as a reasonable lower bound on the minimum rate of gene flow in this system. We assumed a single, chromosome (or genomic segment) of length 1 megabase (Mb) with a per base mutation rate of $1 \times 10^{-7}$. We set the recombination rate between adjacent bases to $1 \times 10^{-8}$. Simulations were run for an initial 100,000 generations with the large panmictic population to allow genetic variation to accumulate (i.e., to reach mutation-drift equilibrium). This population was then divided into the 36 (sub)populations of size $N_e = 173$. We allowed the population to evolve for an additional 200,000 generations. We recorded nucleotide diversity ($\pi$) every 500 generations in the original population and one of the non-edge (sub)populations after the split. Genotypes were extracted from each population at generations 150,000 and 150,002 (50,000 generations after the split) to obtain estimates of the variance $N_e$ for each (sub)population. This was done using `varne` (version 0.1) as described for the actual data in the main text (Gompert & Messina, 2016b). Ten replicate simulations were conducted under each set of conditions.

### Supplemental Tables and Figures

Table S1: Census population size estimates. S.E. denotes standard error. Density estimates were converted to census sizes by multiplying by the geographic area of each population and then by three to account for the length of the flight season.

| Population | Year | Density (per km$^2$) | S.E. | Census estimate |
|---|---|---|---|---|
| BCR | 2013 | 8963 | 1956 | 2382 |
| | 2014 | 4672 | 1442 | 1241.7 |
| BNP | 2013 | 3084 | 1034 | 633.9 |
| | 2014 | 6196 | 1727 | 1273.2 |
| | 2018 | 5835 | 1782 | 1199.1 |
| BTB | 2013 | 4275 | 1542 | 1838.7 |
| | 2014 | 4600 | 1483 | 1978.5 |
| | 2016 | 6425 | 2006 | 2763.3 |
| | 2017 | 2686 | 944 | 1155.3 |
| | 2018 | 4289 | 1244 | 1844.8 |
| GNP | 2013 | 5281 | 1527 | 1119.9 |
| | 2014 | 4813 | 1511 | 1024.5 |
| | 2017 | 1611 | 773 | 343 |
| | 2018 | 2171 | 580 | 462.2 |
| HNV | 2014 | 5144 | 1668 | 5291.4 |
| MRF | 2014 | 5010 | 2023 | 977.7 |
| PSP | 2014 | 3868 | 1626 | 366.6 |
| | 2017 | 3842 | 1979 | 354.2 |
| SKI | 2014 | 4033 | 1207 | 1348.8 |
| | 2016 | 3714 | 1471 | 1242.2 |
| | 2017 | 2076 | 814 | 694.4 |
| | 2018 | 9780 | 2262 | 3271.1 |
| USL | 2014 | 4285 | 1234 | 1708.2 |
| | 2016 | 7342 | 2894 | 2927.1 |
| | 2018 | 2496 | 1066 | 995.1 |

Table S2: Maximum likelihood demographic parameter estimates from $\delta$a$\delta$i. $\Theta$ denotes the ancestral mutation-scaled long-term effective population size ($\theta = 4N_{ref}\mu$, where $\mu$ is the total mutation rate across the GBS loci), $s$ denotes the proportion of the ancestral population that founded derived population 1 (1-$s$ gives the proportion founding derived population 2), $\nu_1$ and $\nu_2$ denote the size of populations 1 and 2 at the present relative to the ancestral population, $T_{split}$ is the split time in $2 \times N_{ref}$ generations, $M_{12}$ is number of migrants per generation from population 2 to 1 ($M_{12}$ is defined similarly). Contemporary migration rates in terms of numbers of migrants can be obtained as $Nm_{12} = \nu_1 M_{12}/2$ and $Nm_{21} = \nu_2 M_{21}/2$.

| Populations | $\Theta$ | $s$ | $\nu_1$ | $\nu_2$ | $T_{split}$ | $M_{12}$ | $M_{21}$ |
|---|---|---|---|---|---|---|---|
| BCR × BNP | 1744.58 | 0.50 | 1.77 | 1.54 | 0.010 | 0.30 | 0.31 |
| BCR × BTB | 1788.12 | 0.50 | 0.87 | 4.07 | 0.003 | 0.50 | 0.50 |
| BCR × HNV | 1767.78 | 0.50 | 1.39 | 1.29 | 0.009 | 0.30 | 0.30 |
| BCR × PSP | 1716.44 | 0.50 | 1.89 | 1.41 | 0.006 | 0.34 | 0.35 |
| BCR × RNV | 1637.53 | 0.50 | 4.77 | 1.94 | 0.010 | 0.26 | 0.26 |
| BCR × SKI | 1789.67 | 0.50 | 0.80 | 2.89 | 0.003 | 0.31 | 0.45 |
| BCR × USL | 1810.69 | 0.50 | 0.59 | 1.56 | 0.003 | 0.00 | 0.46 |
| BNP × BTB | 1814.08 | 0.50 | 0.91 | 2.43 | 0.008 | 0.36 | 0.36 |
| BNP × GNP | 1647.20 | 0.50 | 7.41 | 0.73 | 0.006 | 0.44 | 0.44 |
| BNP × HNV | 1711.07 | 0.50 | 2.58 | 2.57 | 0.007 | 0.29 | 0.29 |
| BNP × PSP | 1721.93 | 0.50 | 1.70 | 1.55 | 0.012 | 0.24 | 0.24 |
| BNP × RNV | 1640.15 | 0.53 | 3.21 | 1.97 | 0.016 | 0.21 | 0.21 |
| BNP × SKI | 1824.79 | 0.50 | 0.78 | 1.68 | 0.008 | 0.39 | 0.39 |
| BNP × USL | 1855.01 | 0.50 | 0.57 | 1.25 | 0.006 | 0.45 | 0.42 |
| BTB × GNP | 1797.87 | 0.50 | 2.95 | 0.43 | 0.010 | 0.44 | 0.50 |
| BTB × HNV | 1828.96 | 0.50 | 1.91 | 0.79 | 0.007 | 0.36 | 0.38 |
| BTB × MRF | 1694.04 | 0.50 | 11.46 | 7.09 | 0.008 | 0.27 | 0.26 |
| BTB × PSP | 1776.44 | 0.50 | 4.14 | 0.76 | 0.004 | 0.35 | 0.47 |
| BTB × RNV | 1709.59 | 0.50 | 13.34 | 0.96 | 0.007 | 0.34 | 0.35 |
| BTB × SKI | 1836.53 | 0.50 | 1.24 | 1.07 | 0.002 | 0.13 | 0.48 |
| BTB × USL | 1842.66 | 0.48 | 1.09 | 0.95 | 0.002 | 0.24 | 0.53 |
| GNP × HNV | 1685.73 | 0.50 | 0.60 | 2.99 | 0.007 | 0.45 | 0.45 |
| GNP × SKI | 1812.59 | 0.50 | 0.38 | 2.11 | 0.010 | 0.50 | 0.50 |
| HNV × PSP | 1737.72 | 0.50 | 1.61 | 1.46 | 0.011 | 0.30 | 0.30 |
| HNV × SKI | 1840.04 | 0.50 | 0.70 | 1.53 | 0.007 | 0.40 | 0.41 |
| HNV × USL | 1857.83 | 0.50 | 0.54 | 1.22 | 0.006 | 0.48 | 0.48 |
| MRF × PSP | 1591.74 | 0.44 | 18.72 | 4.26 | 0.013 | 0.21 | 0.22 |
| MRF × SKI | 1699.74 | 0.50 | 6.41 | 8.07 | 0.008 | 0.22 | 0.21 |
| MRF × USL | 1730.33 | 0.50 | 4.85 | 4.68 | 0.007 | 0.09 | 0.26 |
| PSP × RNV | 1619.40 | 0.50 | 3.72 | 2.05 | 0.011 | 0.28 | 0.28 |
| PSP × SKI | 1769.21 | 0.50 | 0.75 | 3.31 | 0.004 | 0.00 | 0.44 |
| PSP × USL | 1804.00 | 0.50 | 0.42 | 1.76 | 0.003 | 0.48 | 0.38 |
| RNV × SKI | 1718.86 | 0.50 | 0.86 | 7.73 | 0.007 | 0.37 | 0.37 |
| RNV × USL | 1745.82 | 0.50 | 0.66 | 4.81 | 0.006 | 0.45 | 0.44 |
| SKI × USL | 1847.88 | 0.63 | 0.78 | 0.69 | 0.001 | 0.11 | 0.65 |

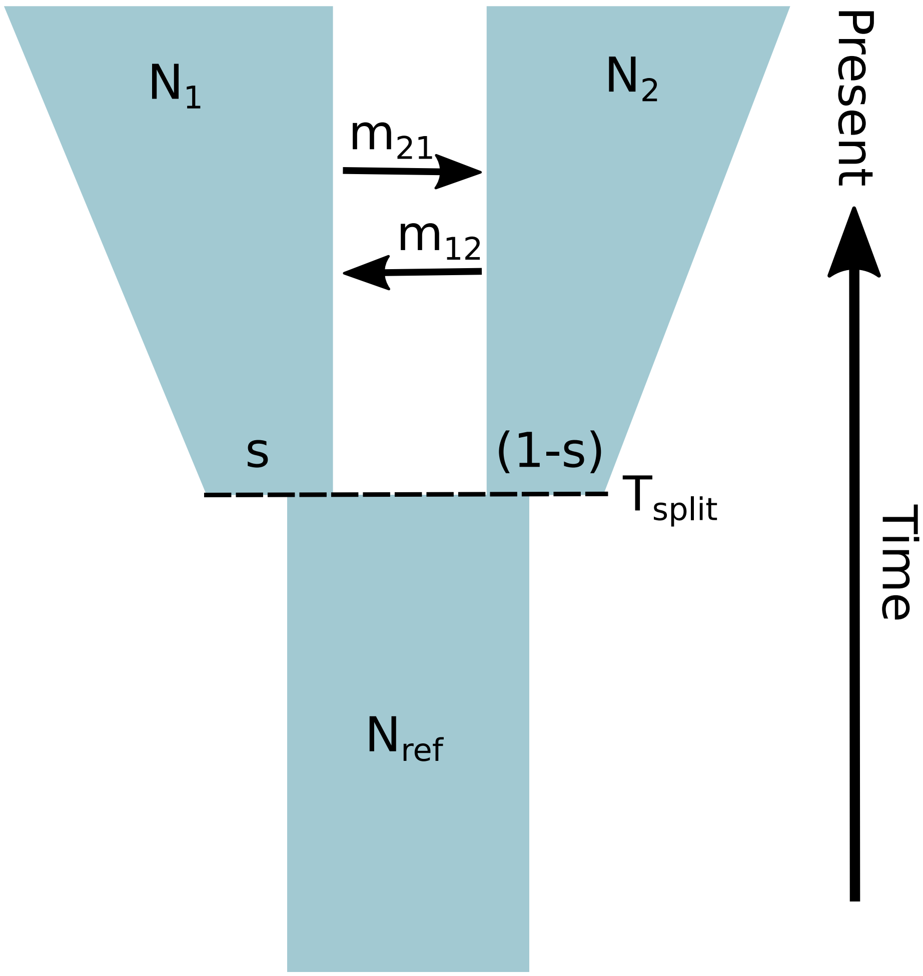


Figure S1: Diagram representing the isolation with migration (IM) model. At time $T_{split}$ in the past (measured in $2N_{ref}$ generations), a population of size $N_{ref}$ splits into two populations, with $s \times N_{ref}$ going to population 1 and $(1-s) \times N_{ref}$ going to population 2. The populations then grow to sizes $N_1$ and $N_2$ with migration rates (proportions) $m_{12}$ and $m_{21}$ ($m_{12}$ is the migration rate from population 2 to population 1). Actual inferences are made of the following composite parameters: $\nu_1 = N_1/N_{ref}$, $\nu_2 = N_2/N_{ref}$, $M_{12} = 2N_{ref}m_{12}$, $M_{21} = 2N_{ref}m_{21}$, and $\theta = 4N_{ref}\mu$.

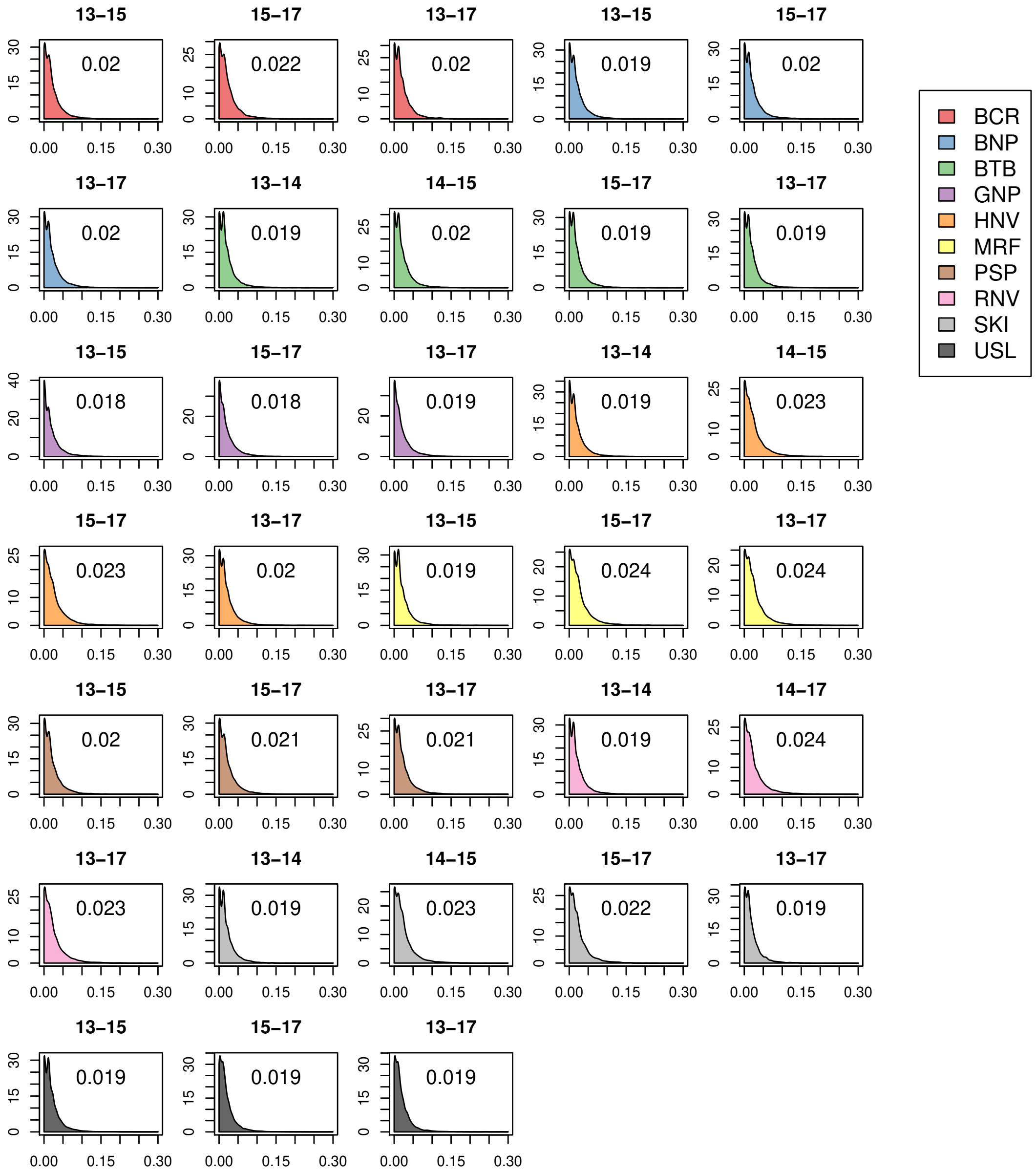


Figure S2: Density plots show the distribution of allele frequency change (absolute value of the difference in allele frequency) between different generations. Colors denote different populations. The mean (absolute) change is given within each plot, and the last two digits of the years are provided above each plot.

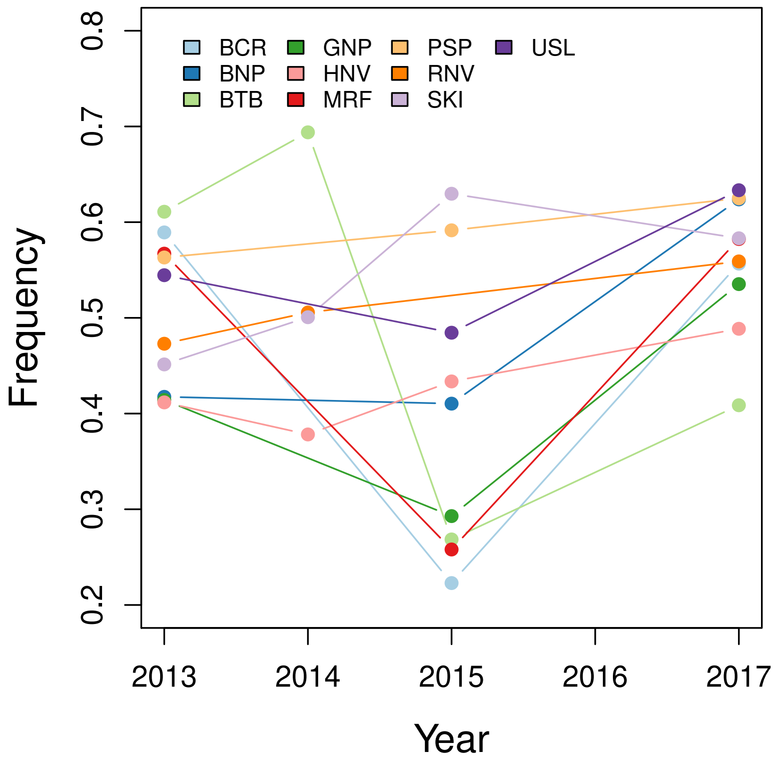


Figure S3: Scatterplot show the allele-frequency time series for a SNP on chromosome 8 (position = 1,042,561, G-C polymorphism), which exhibited the greatest change in BTB between 2013 and 2015.The nearest gene to this SNP (∼100 kilobases away) is *Chitriosidase*.

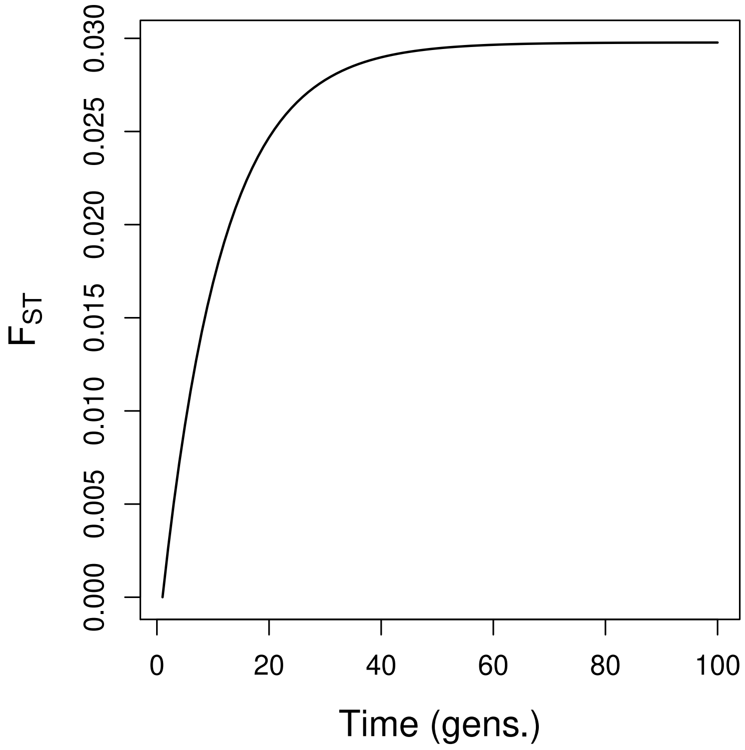


Figure S4: This plot shows the expected accumulation of $F_{ST}$ over time and progress towards drift-migration equilibrium under an island model with rates of gene flow ($Nm = 7.6$) and drift ($N_e = 173$) meant to approximate evolutionary dynamics in *L. idas*.

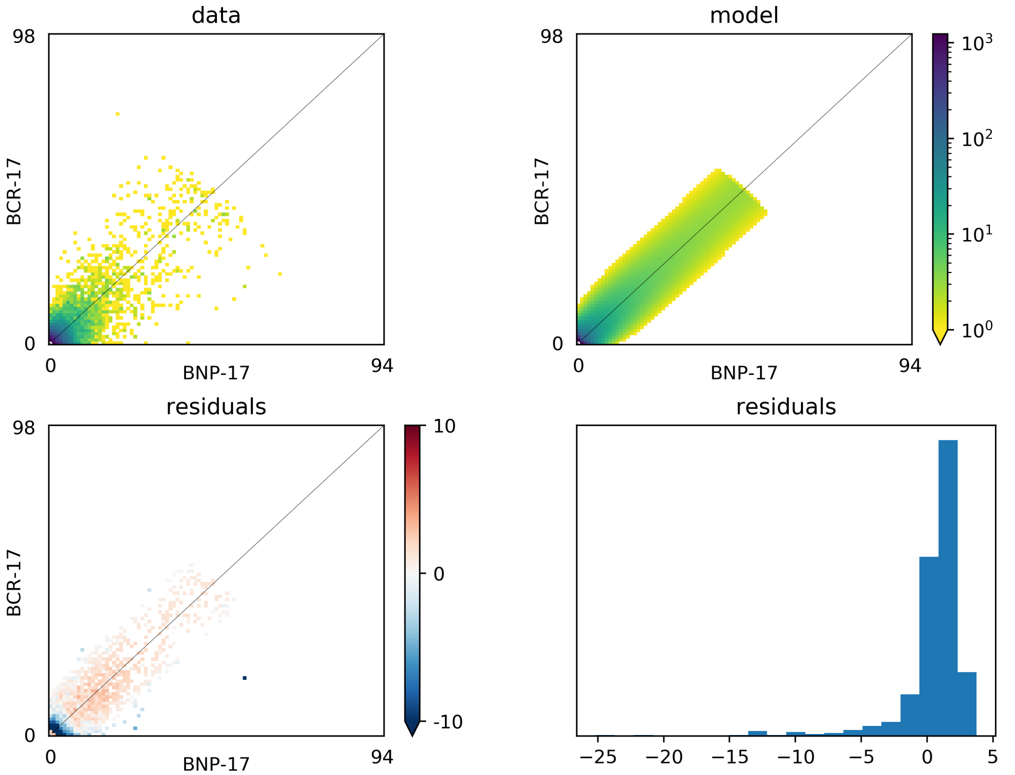


Figure S5: Summary of demographic model fit for BCR and BNP. The top panels show the allele frequency spectra based on the observed data and best-fit model. The bottom panels show the associated residuals.

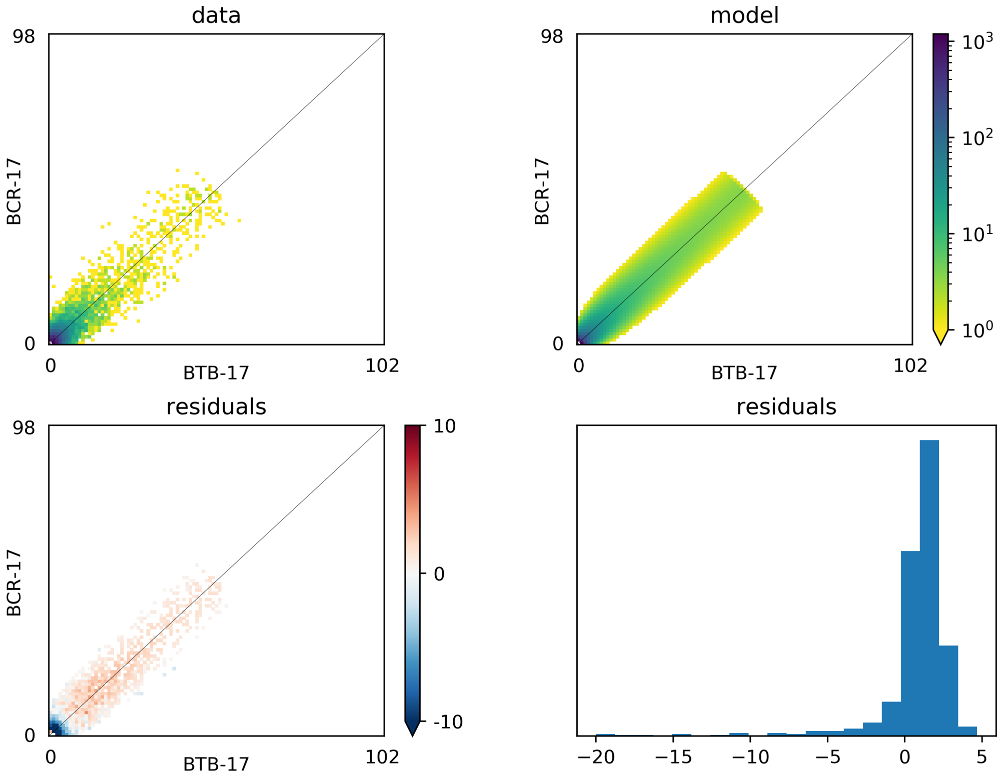


Figure S6: Summary of demographic model fit for BCR and BTB. The top panels show the allele frequency spectra based on the observed data and best-fit model. The bottom panels show the associated residuals.

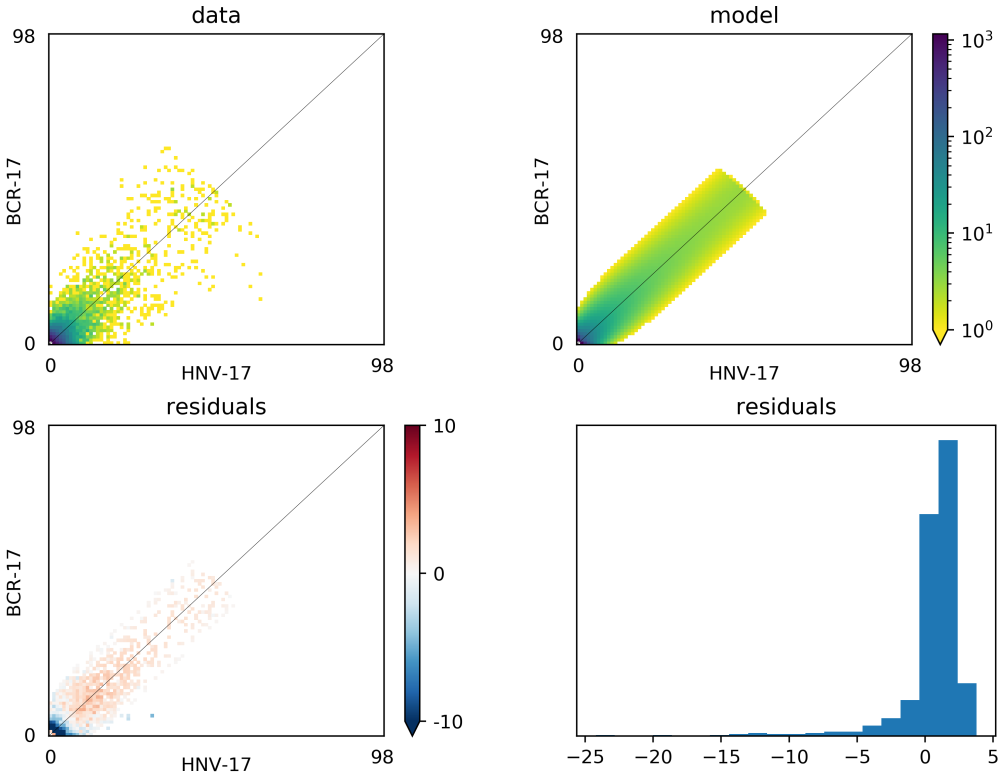


Figure S7: Summary of demographic model fit for BCR and HNV. The top panels show the allele frequency spectra based on the observed data and best-fit model. The bottom panels show the associated residuals.

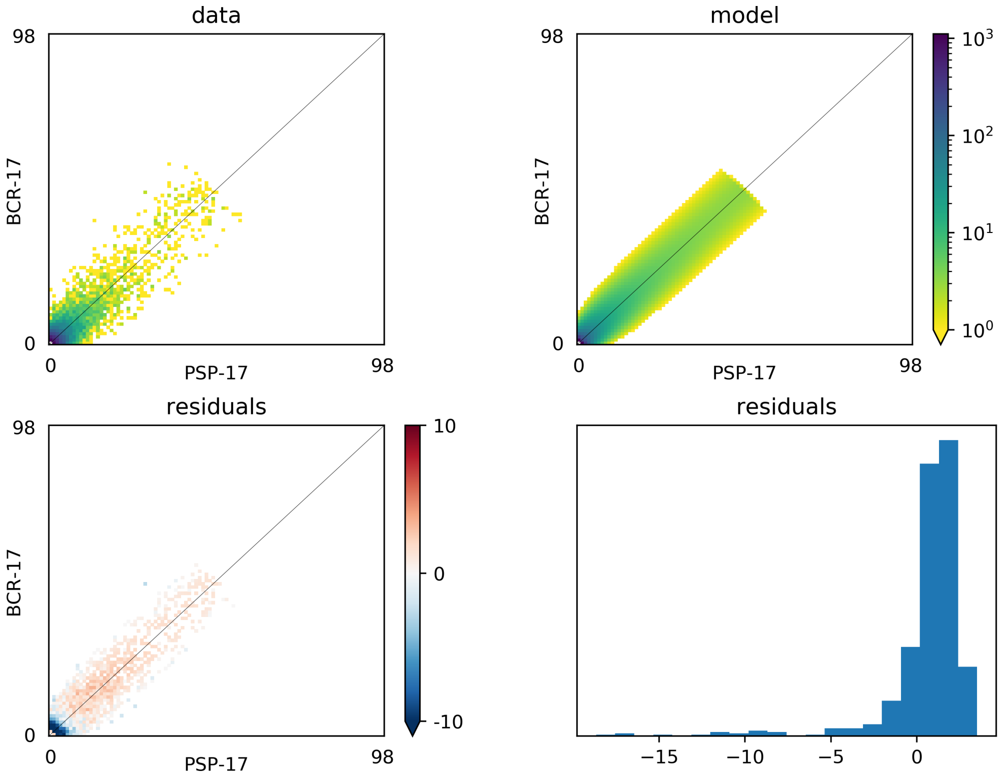


Figure S8: Summary of demographic model fit for BCR and PSP. The top panels show the allele frequency spectra based on the observed data and best-fit model. The bottom panels show the associated residuals.

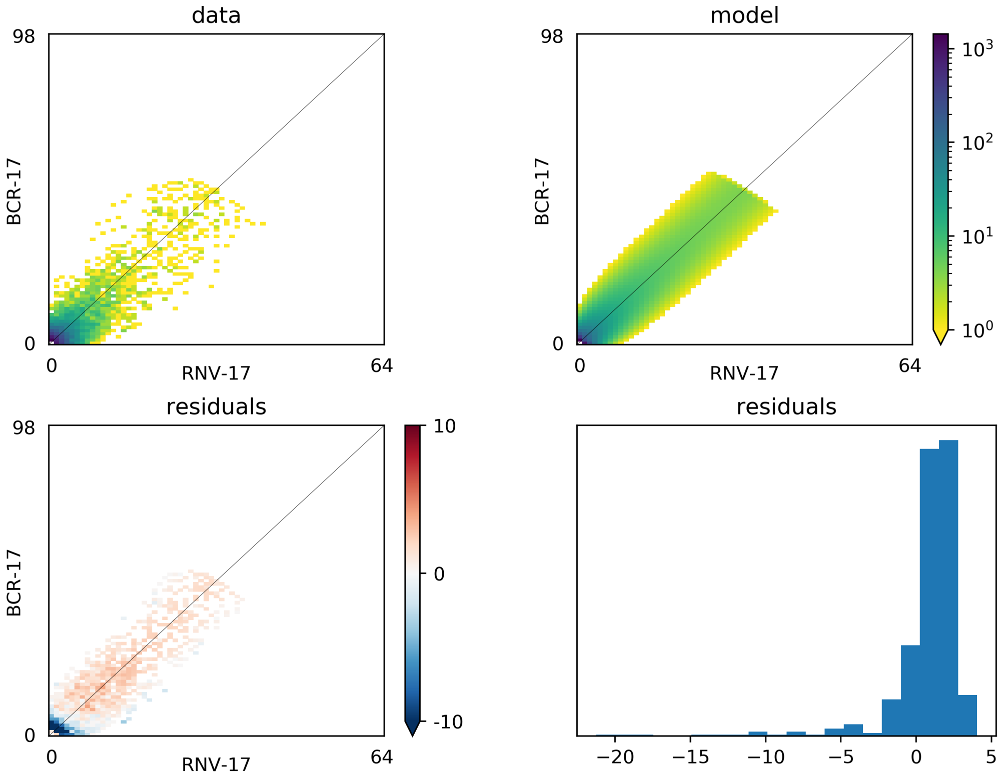


Figure S9: Summary of demographic model fit for BCR and RNV. The top panels show the allele frequency spectra based on the observed data and best-fit model. The bottom panels show the associated residuals.

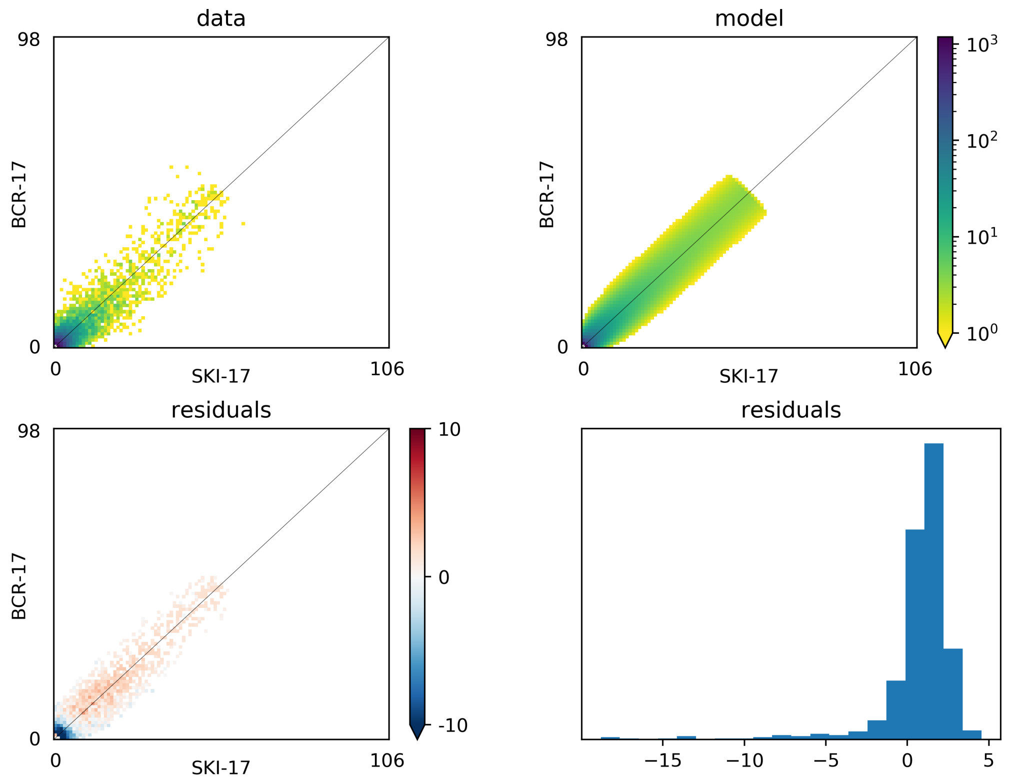


Figure S10: Summary of demographic model fit for BCR and SKI. The top panels show the allele frequency spectra based on the observed data and best-fit model. The bottom panels show the associated residuals.

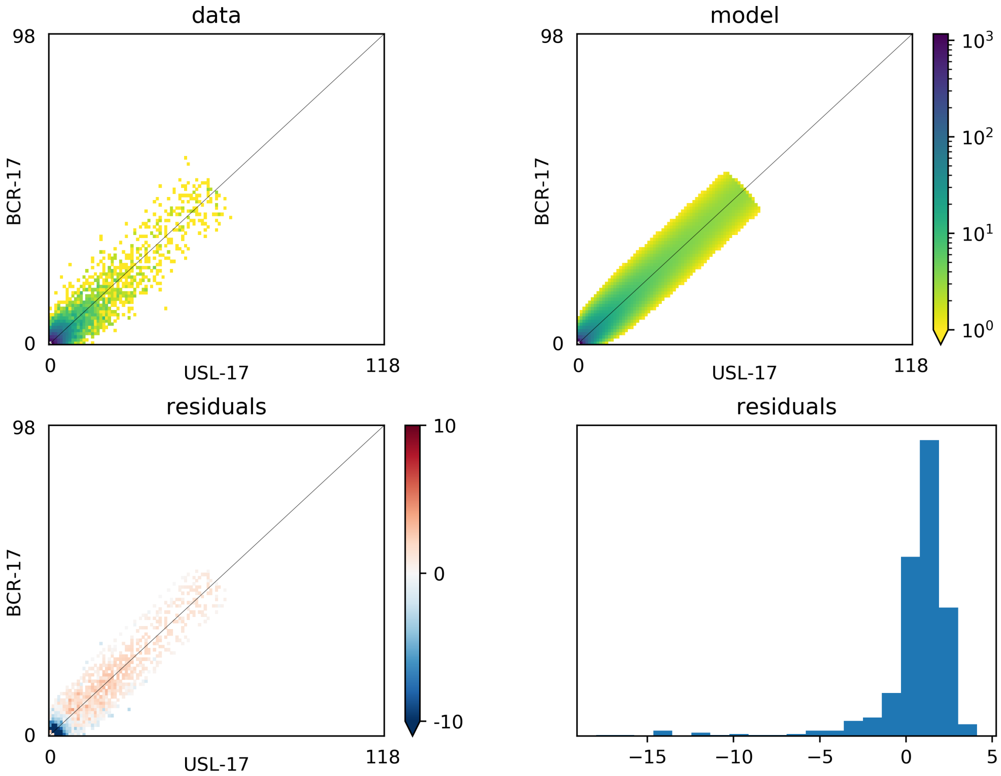


Figure S11: Summary of demographic model fit for BCR and USL. The top panels show the allele frequency spectra based on the observed data and best-fit model. The bottom panels show the associated residuals.

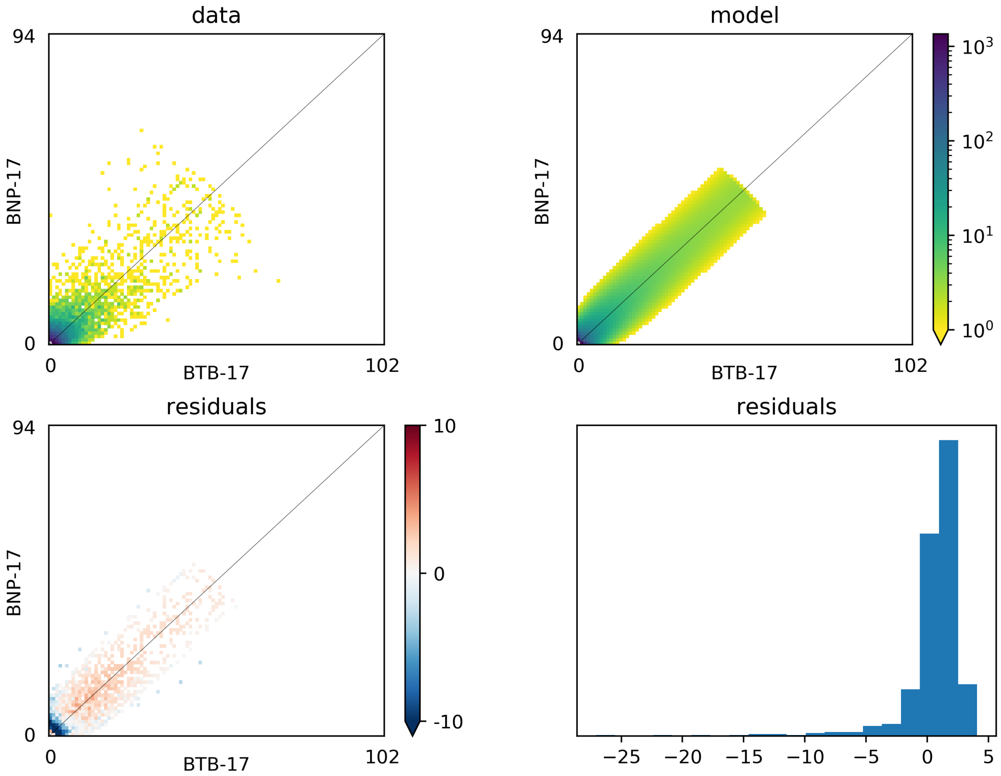


Figure S12: Summary of demographic model fit for BNP and BTB. The top panels show the allele frequency spectra based on the observed data and best-fit model. The bottom panels show the associated residuals.

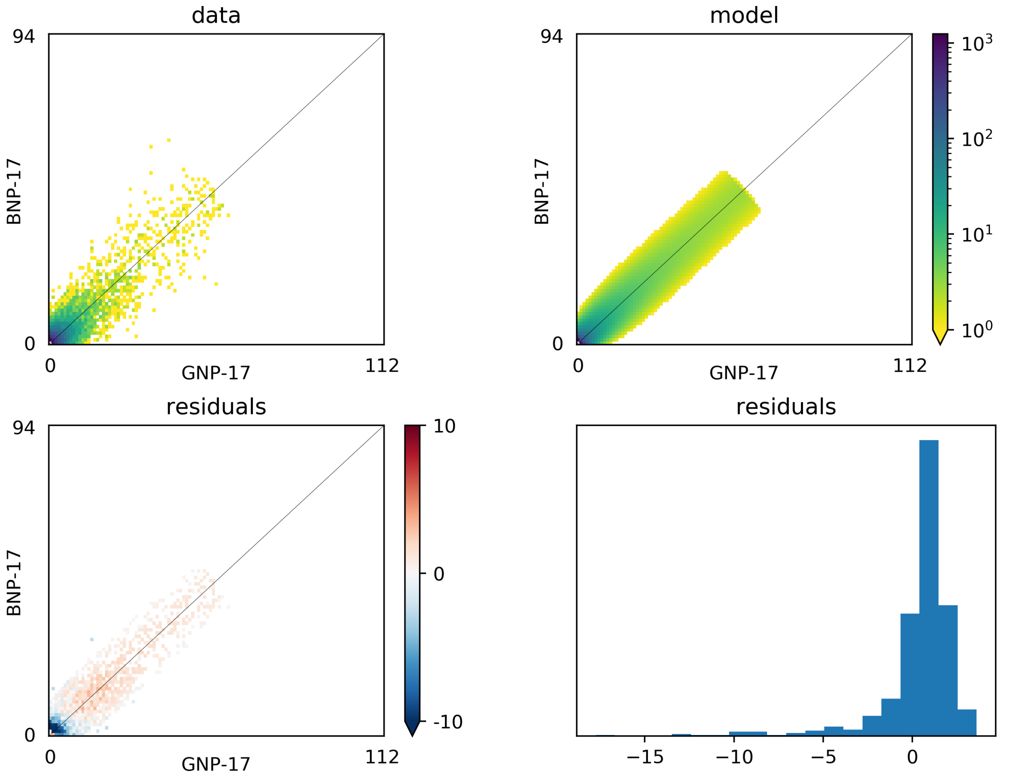


Figure S13: Summary of demographic model fit for BNP and GNP. The top panels show the allele frequency spectra based on the observed data and best-fit model. The bottom panels show the associated residuals.

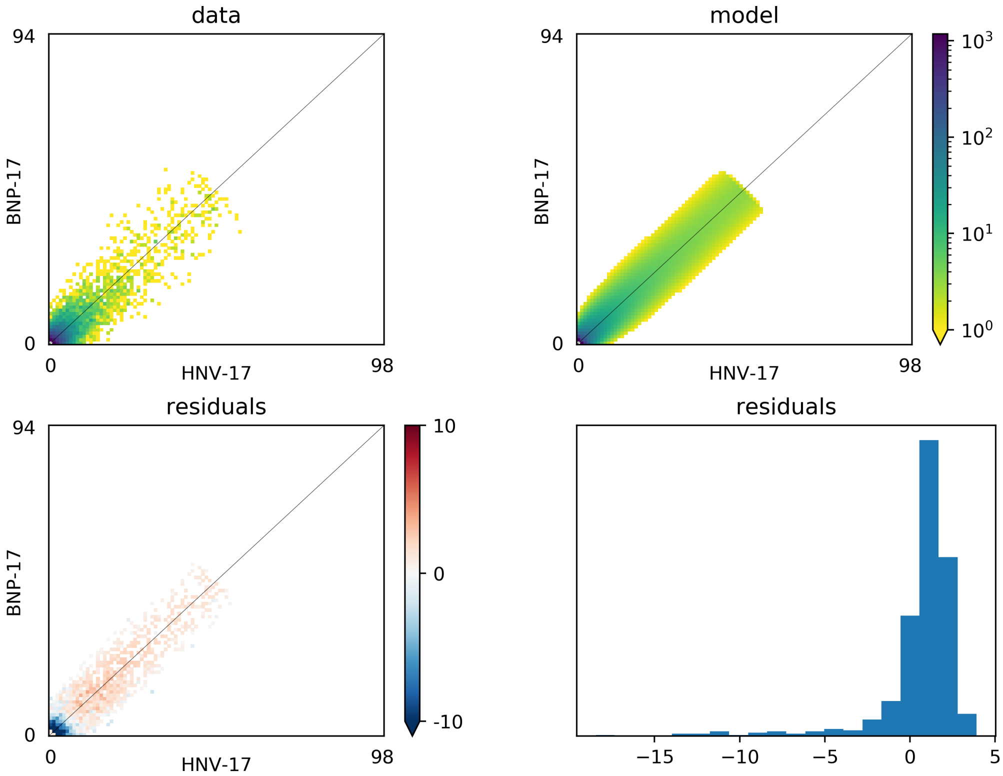


Figure S14: Summary of demographic model fit for BNP and HNV. The top panels show the allele frequency spectra based on the observed data and best-fit model. The bottom panels show the associated residuals.

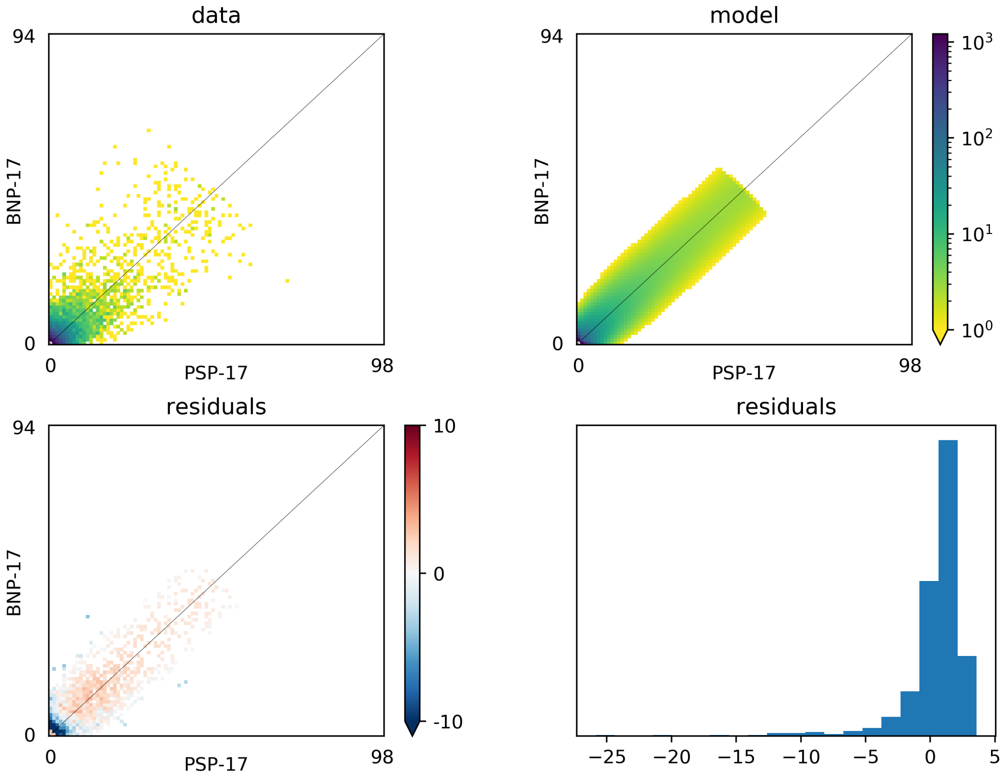


Figure S15: Summary of demographic model fit for BNP and PSP. The top panels show the allele frequency spectra based on the observed data and best-fit model. The bottom panels show the associated residuals.

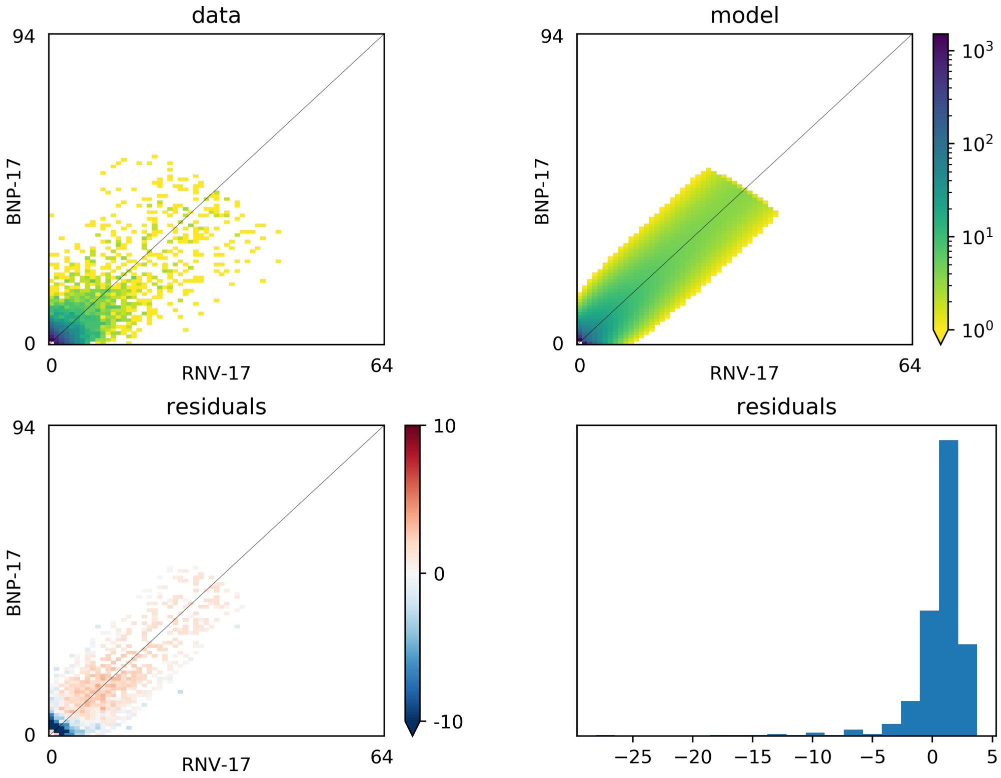


Figure S16: Summary of demographic model fit for BNP and RNV. The top panels show the allele frequency spectra based on the observed data and best-fit model. The bottom panels show the associated residuals.

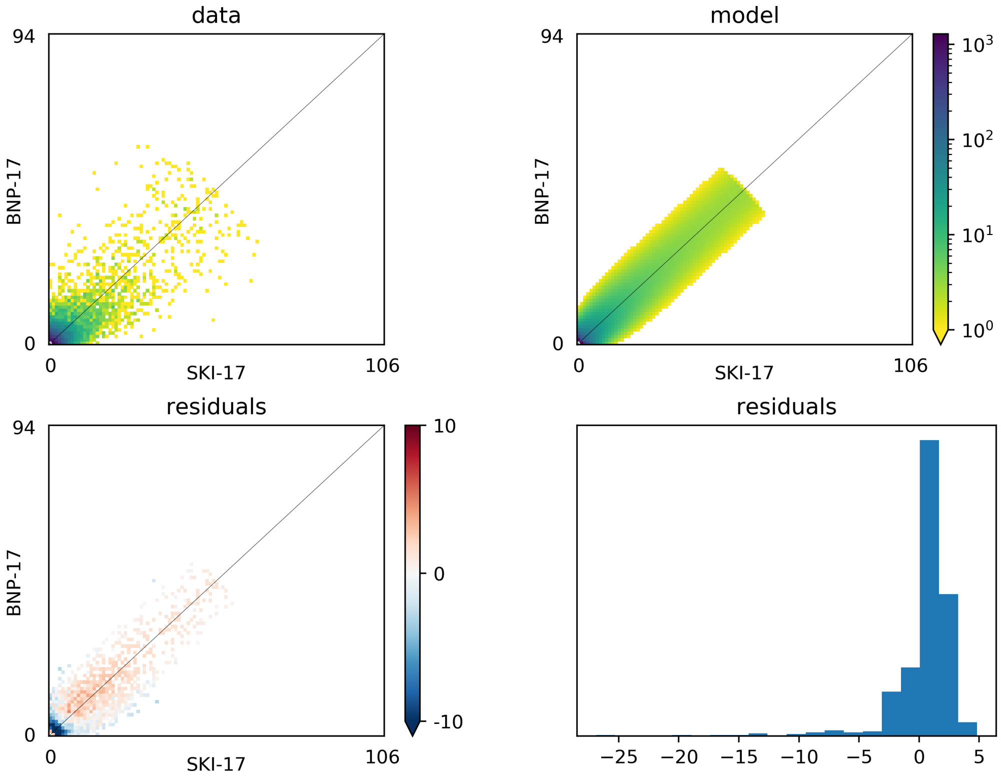


Figure S17: Summary of demographic model fit for BNP and SKI. The top panels show the allele frequency spectra based on the observed data and best-fit model. The bottom panels show the associated residuals.

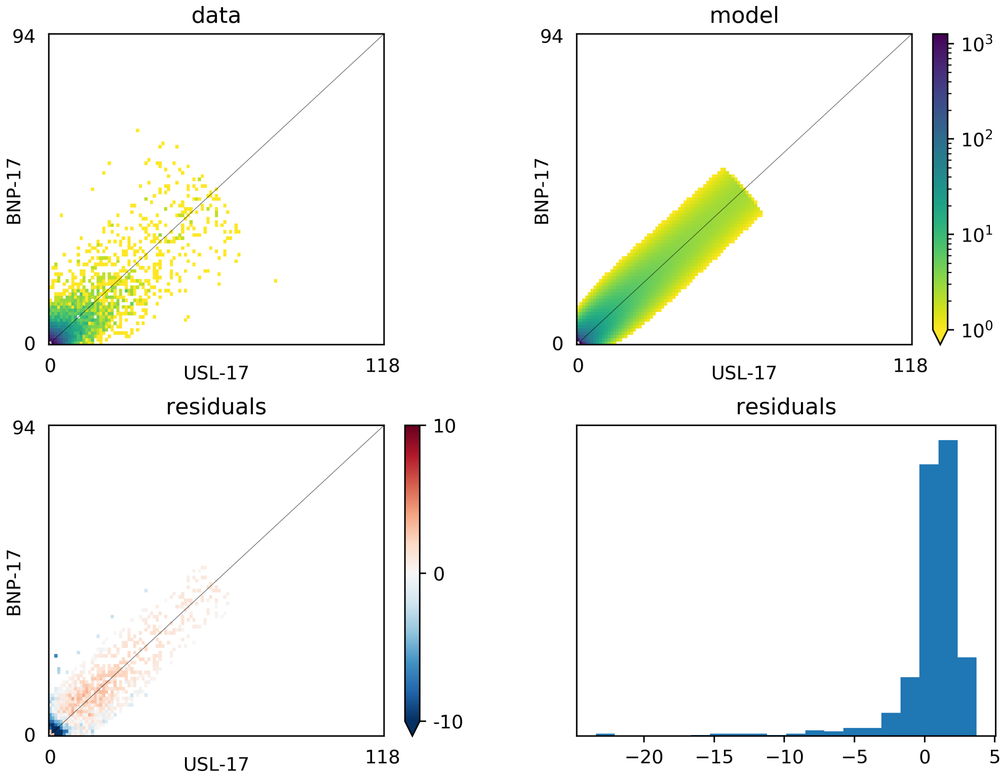


Figure S18: Summary of demographic model fit for BNP and USL. The top panels show the allele frequency spectra based on the observed data and best-fit model. The bottom panels show the associated residuals.

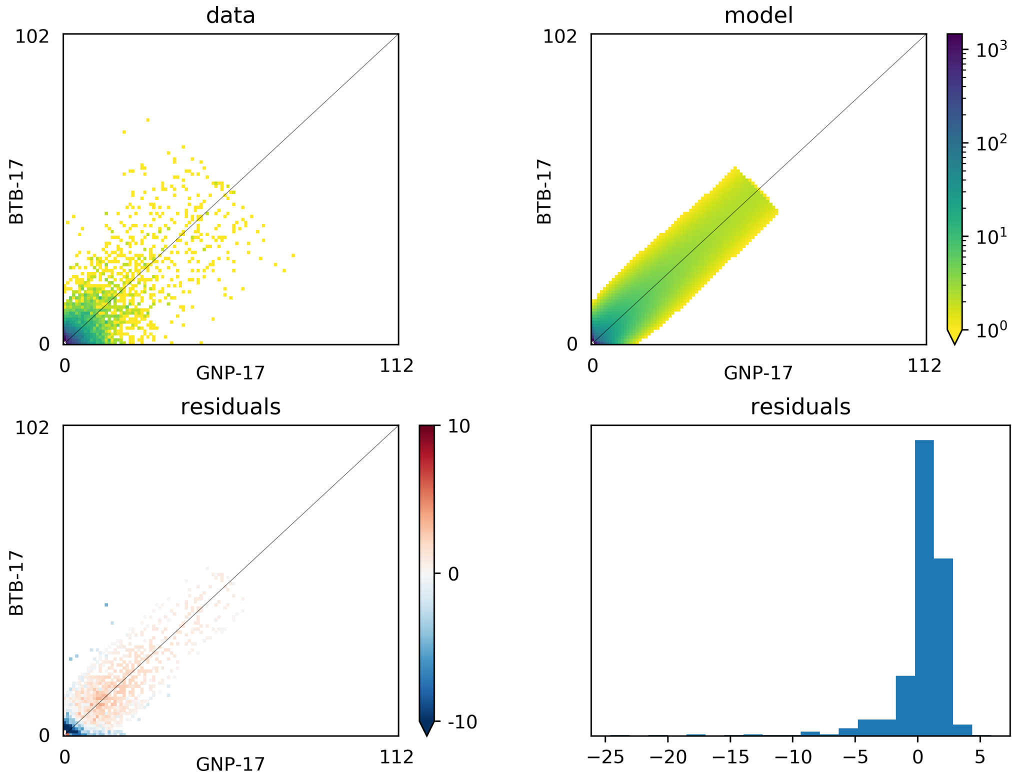


Figure S19: Summary of demographic model fit for BTB and GNP. The top panels show the allele frequency spectra based on the observed data and best-fit model. The bottom panels show the associated residuals.

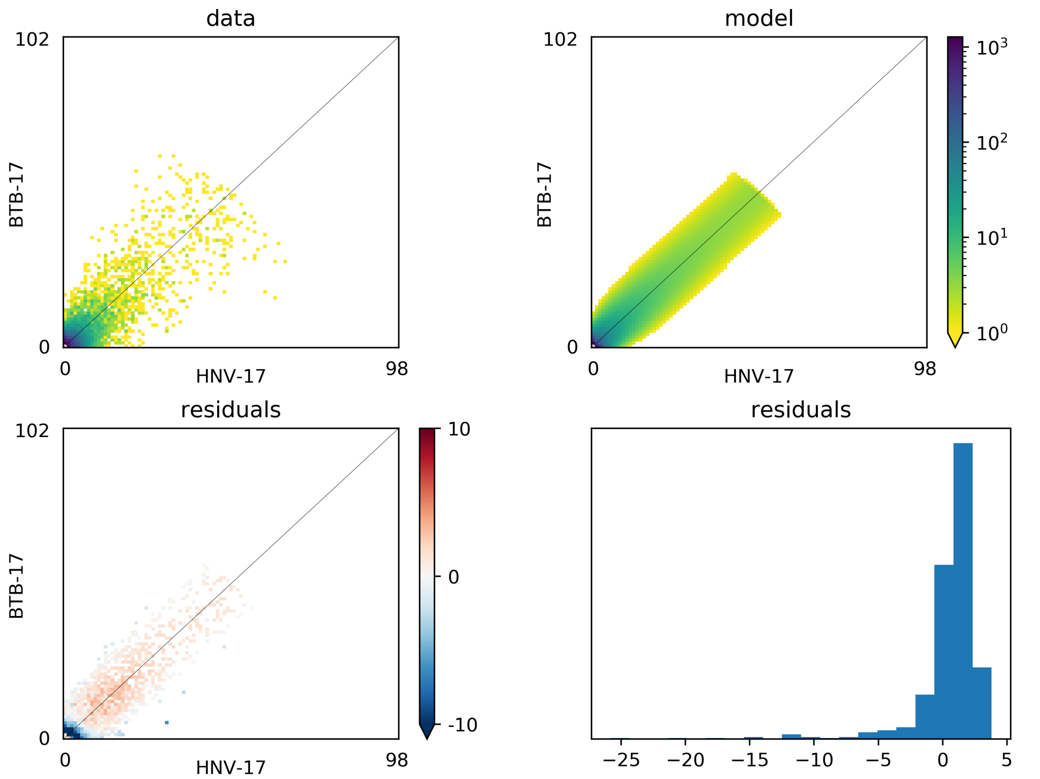


Figure S20: Summary of demographic model fit for BTB and HNV. The top panels show the allele frequency spectra based on the observed data and best-fit model. The bottom panels show the associated residuals.

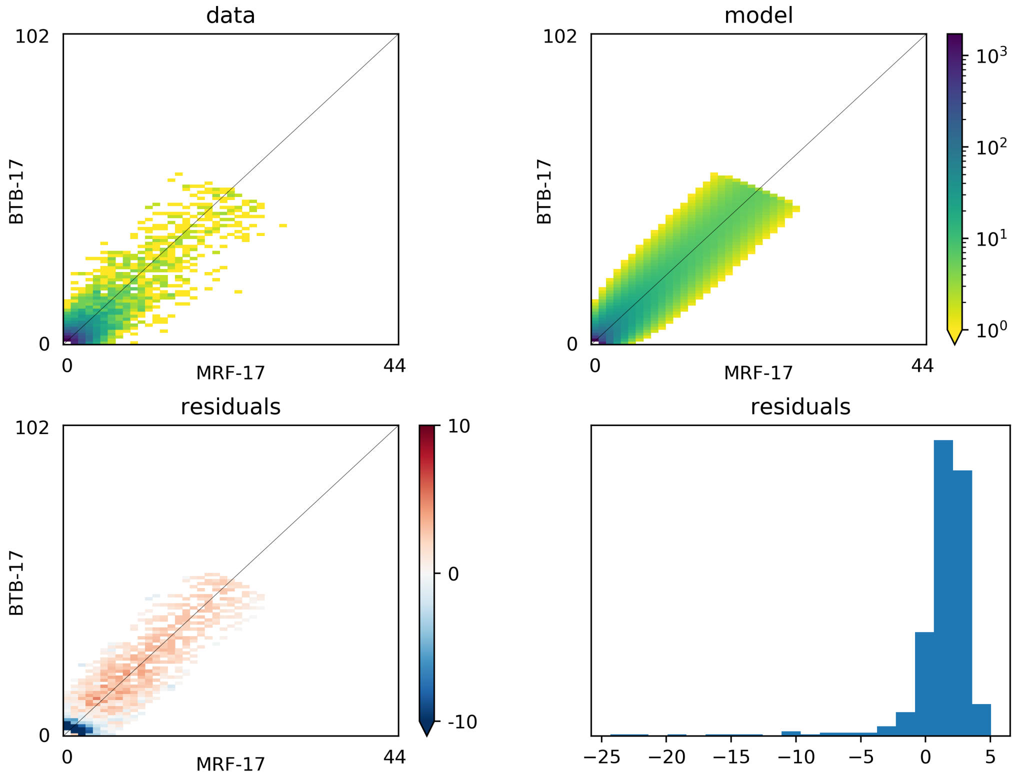


Figure S21: Summary of demographic model fit for BTB and MRF. The top panels show the allele frequency spectra based on the observed data and best-fit model. The bottom panels show the associated residuals.

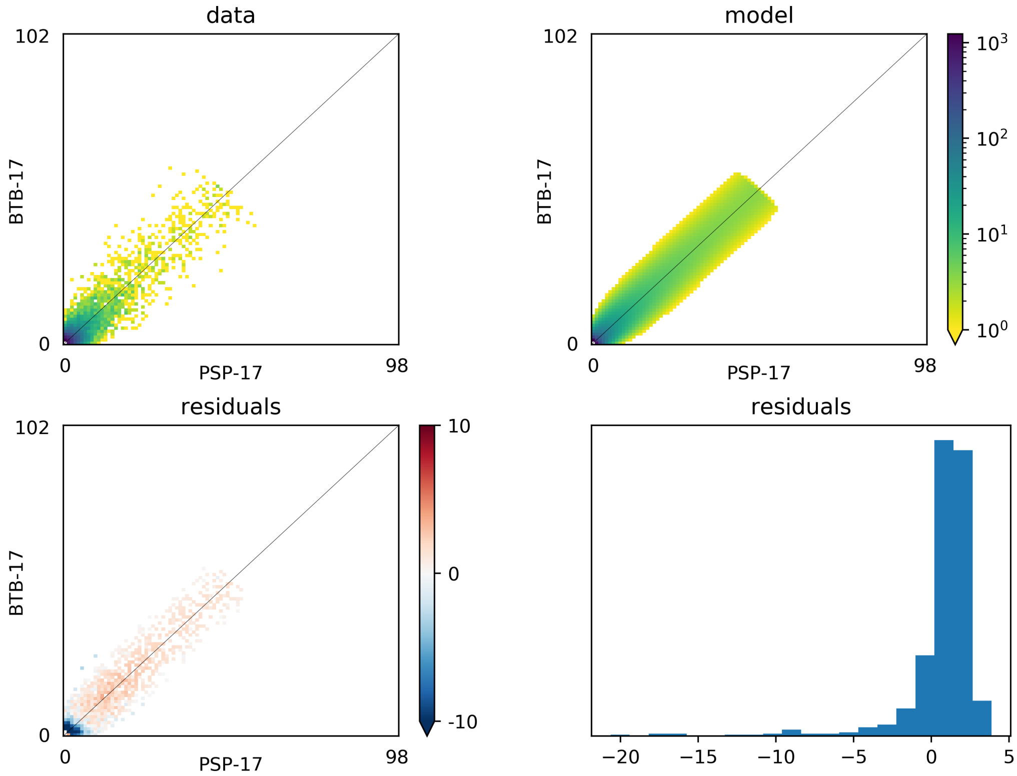


Figure S22: Summary of demographic model fit for BTB and PSP. The top panels show the allele frequency spectra based on the observed data and best-fit model. The bottom panels show the associated residuals.

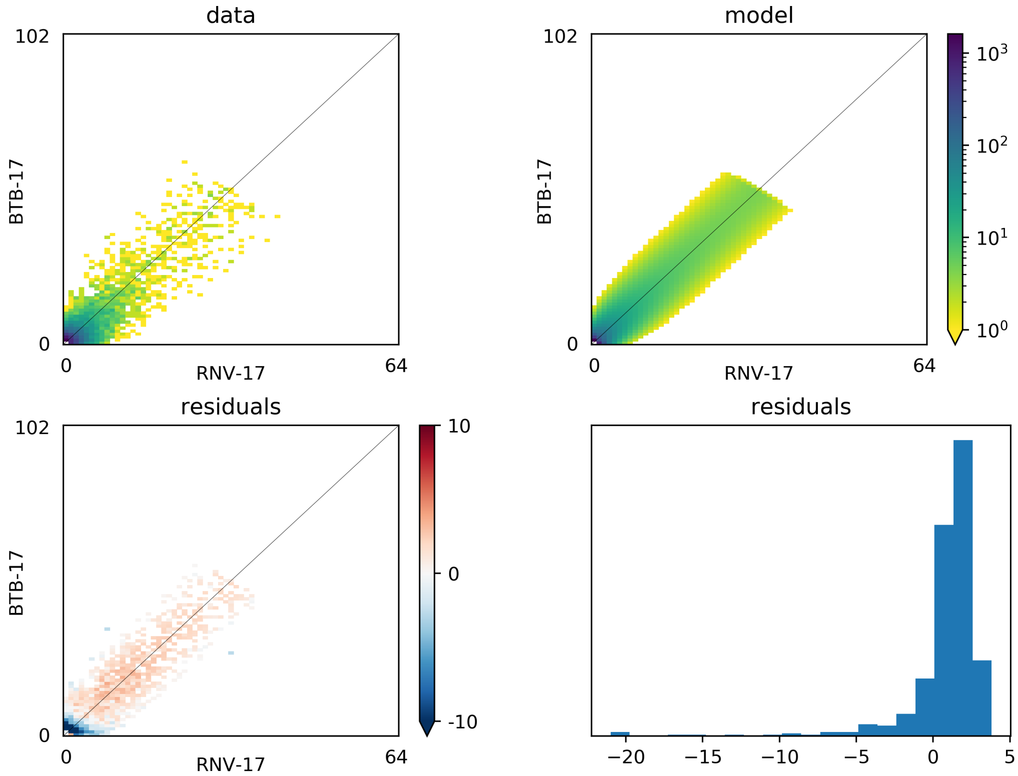


Figure S23: Summary of demographic model fit for BTB and RNV. The top panels show the allele frequency spectra based on the observed data and best-fit model. The bottom panels show the associated residuals.

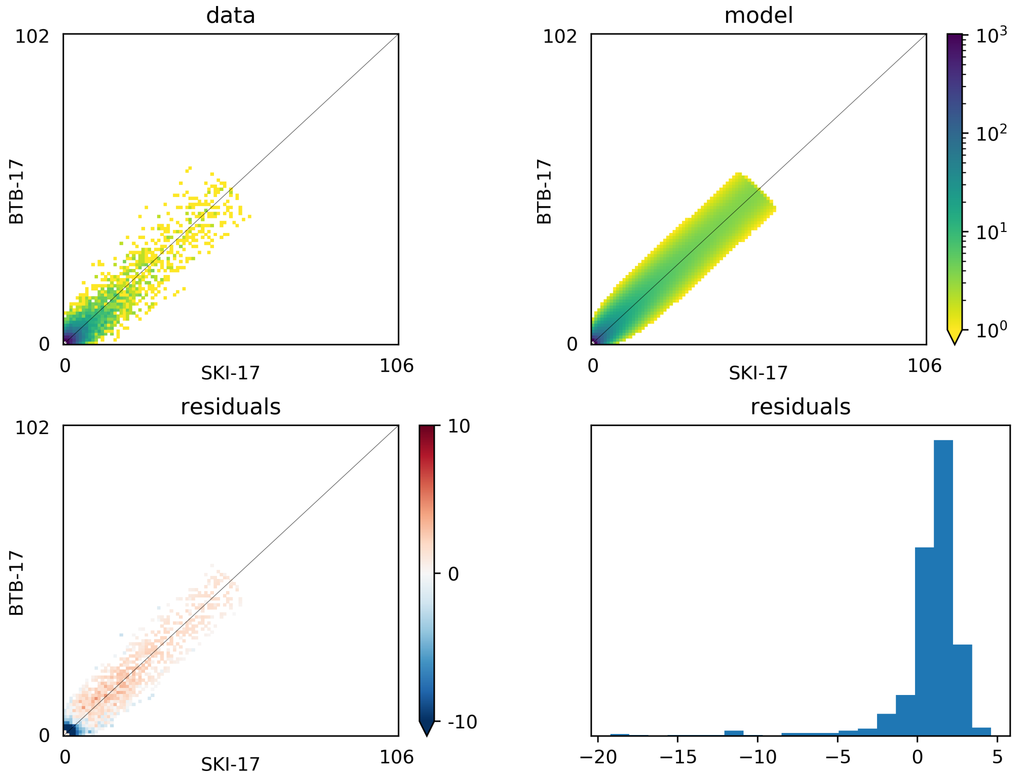


Figure S24: Summary of demographic model fit for BTB and SKI. The top panels show the allele frequency spectra based on the observed data and best-fit model. The bottom panels show the associated residuals.

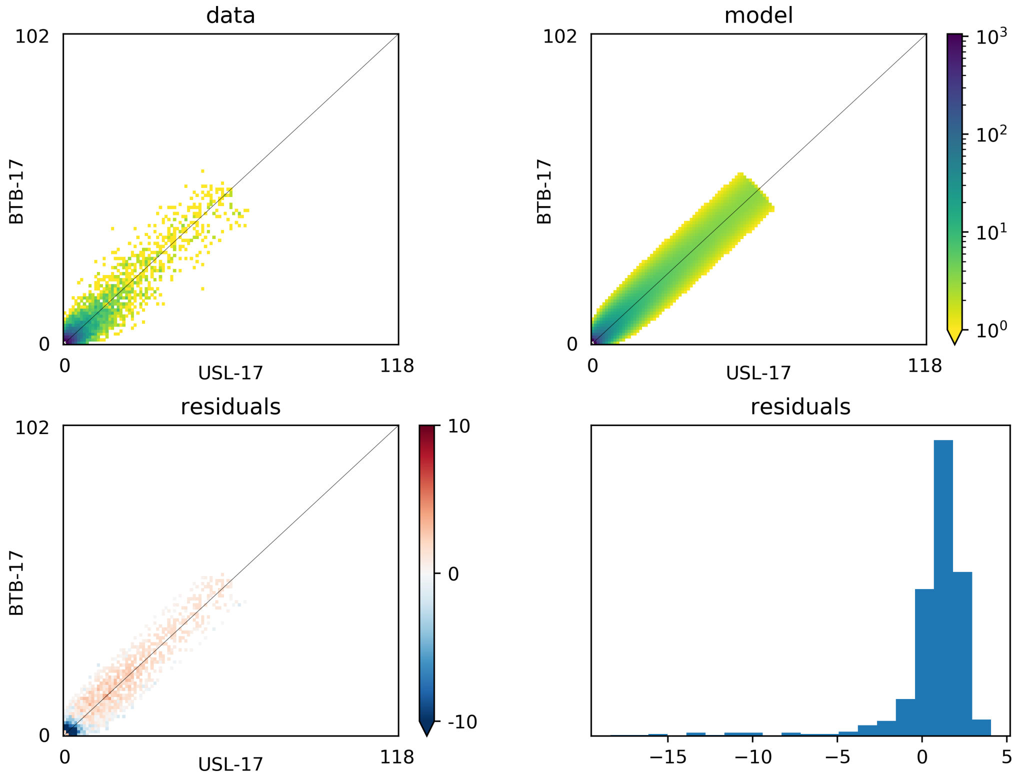


Figure S25: Summary of demographic model fit for BTB and USL. The top panels show the allele frequency spectra based on the observed data and best-fit model. The bottom panels show the associated residuals.

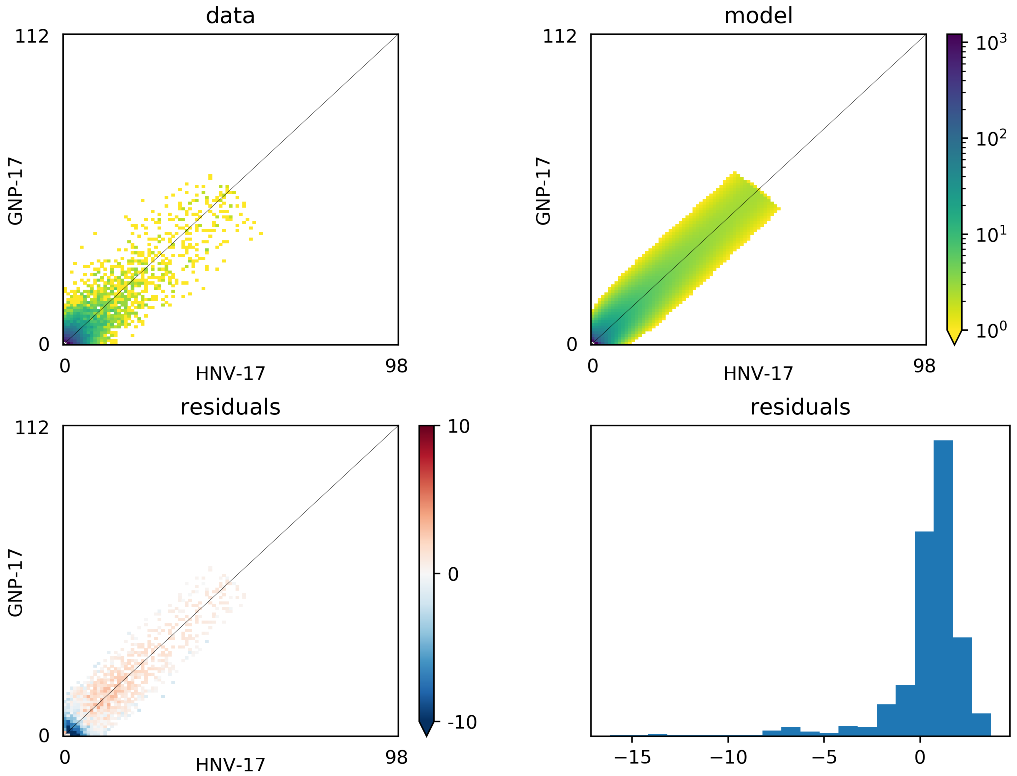


Figure S26: Summary of demographic model fit for GNP and HNV. The top panels show the allele frequency spectra based on the observed data and best-fit model. The bottom panels show the associated residuals.

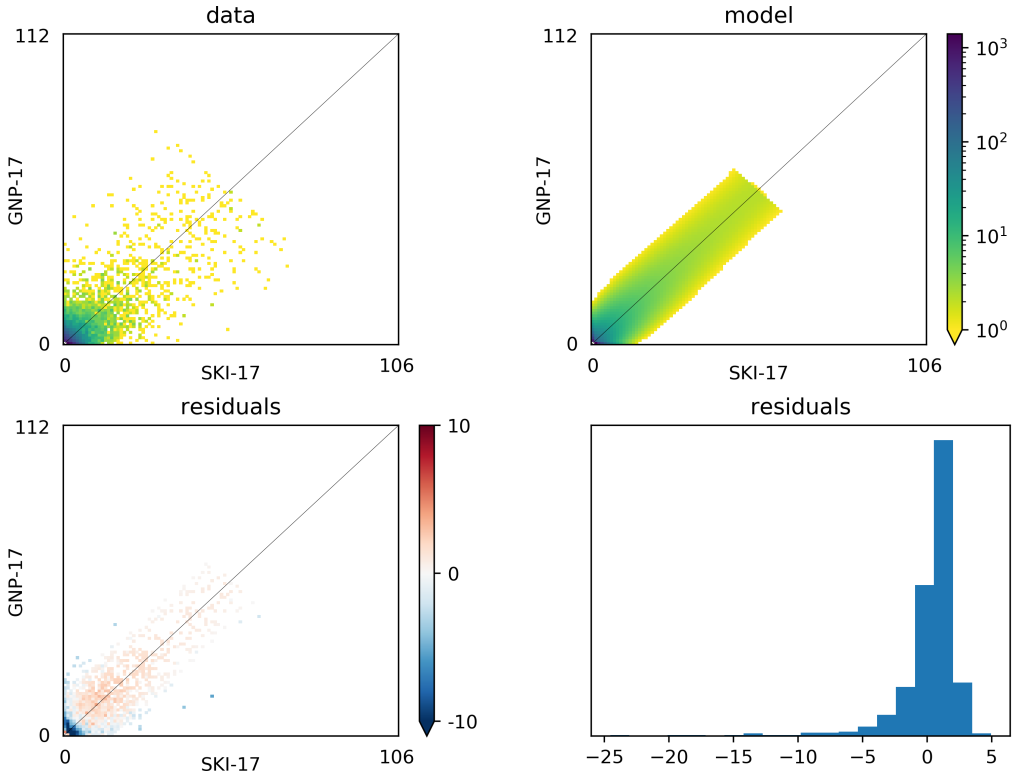


Figure S27: Summary of demographic model fit for GNP and SKI. The top panels show the allele frequency spectra based on the observed data and best-fit model. The bottom panels show the associated residuals.

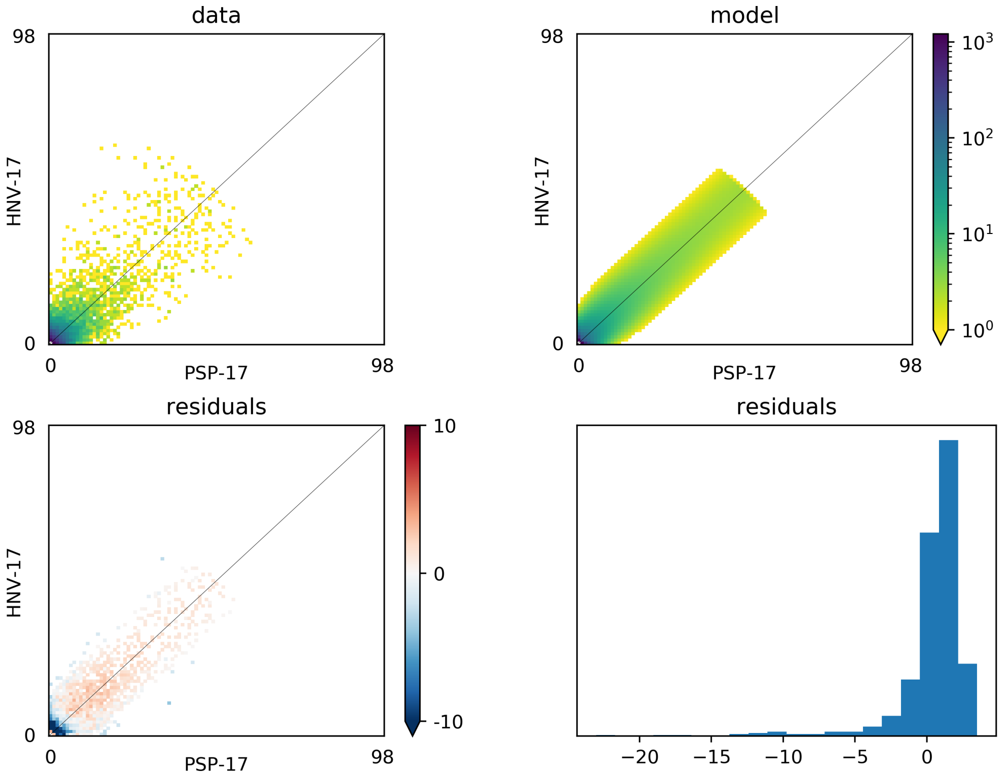


Figure S28: Summary of demographic model fit for HNV and PSP. The top panels show the allele frequency spectra based on the observed data and best-fit model. The bottom panels show the associated residuals.

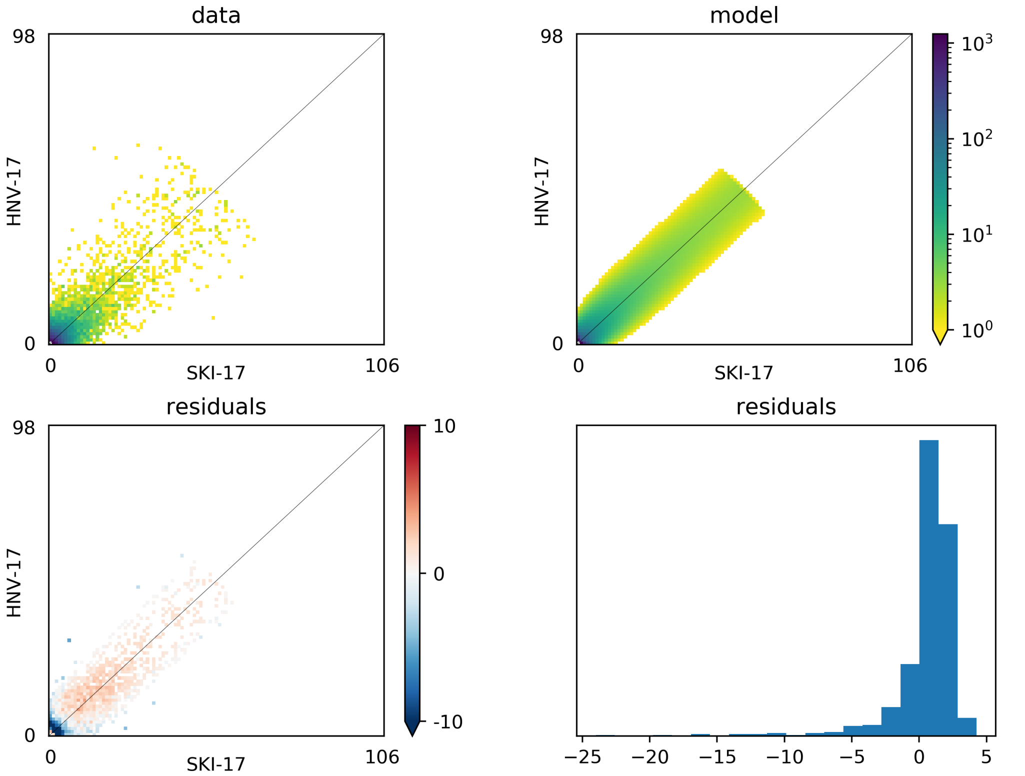


Figure S29: Summary of demographic model fit for HNV and SKI. The top panels show the allele frequency spectra based on the observed data and best-fit model. The bottom panels show the associated residuals.

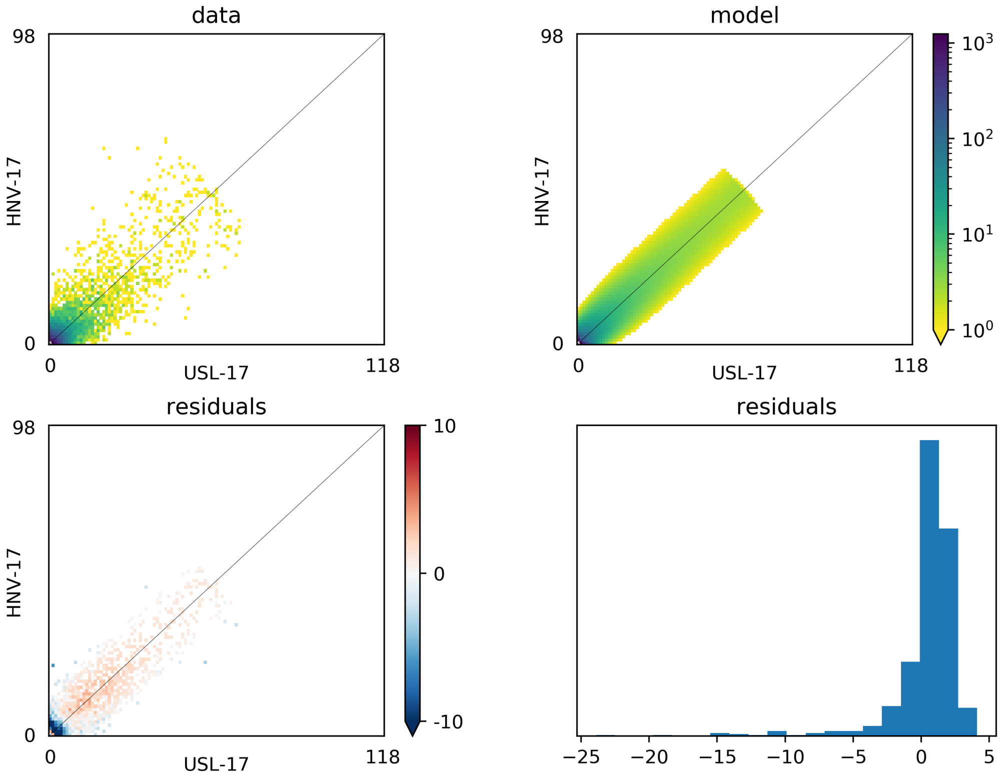


Figure S30: Summary of demographic model fit for HNV and USL. The top panels show the allele frequency spectra based on the observed data and best-fit model. The bottom panels show the associated residuals.

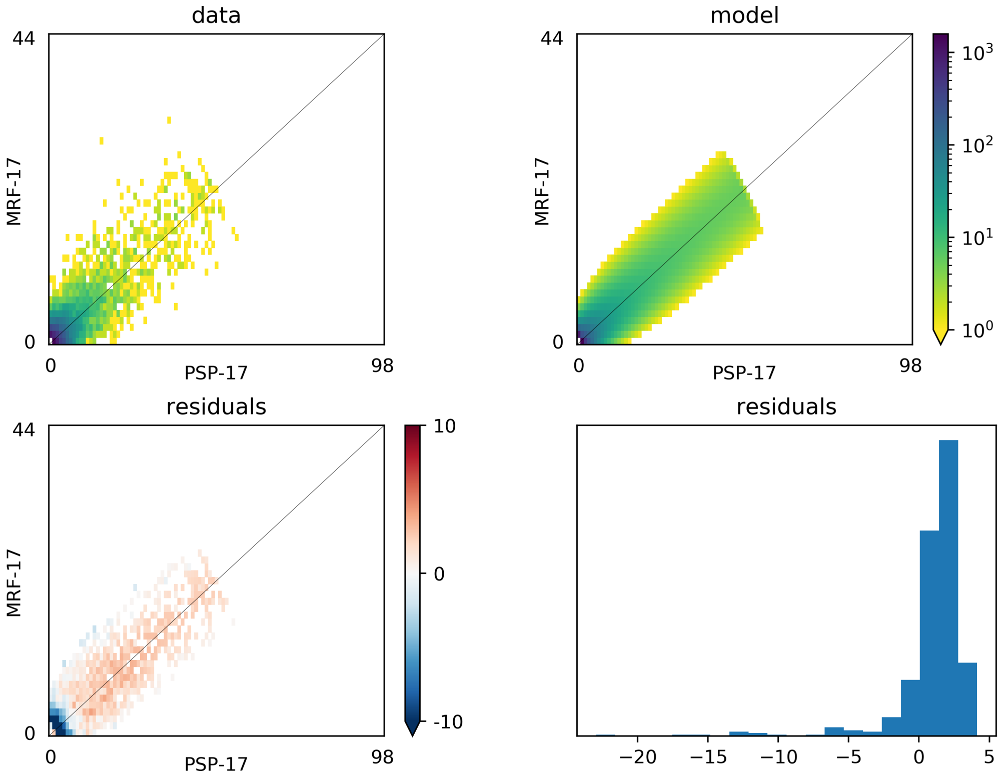


Figure S31: Summary of demographic model fit for MRF and PSP. The top panels show the allele frequency spectra based on the observed data and best-fit model. The bottom panels show the associated residuals.

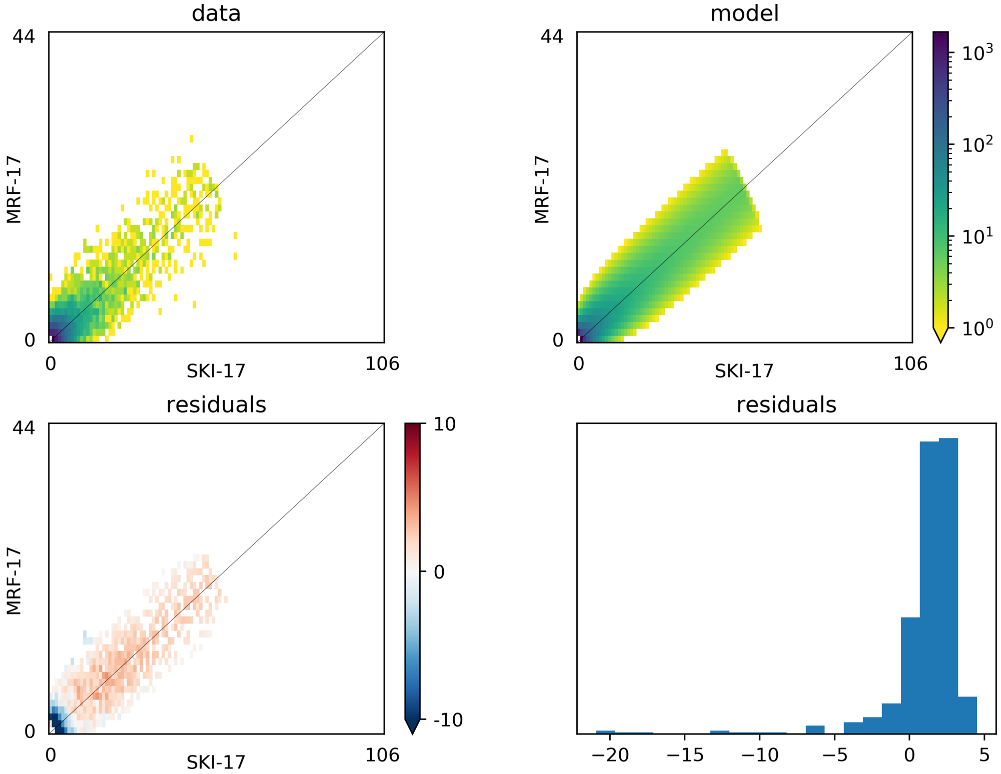


Figure S32: Summary of demographic model fit for MRF and SKI. The top panels show the allele frequency spectra based on the observed data and best-fit model. The bottom panels show the associated residuals.

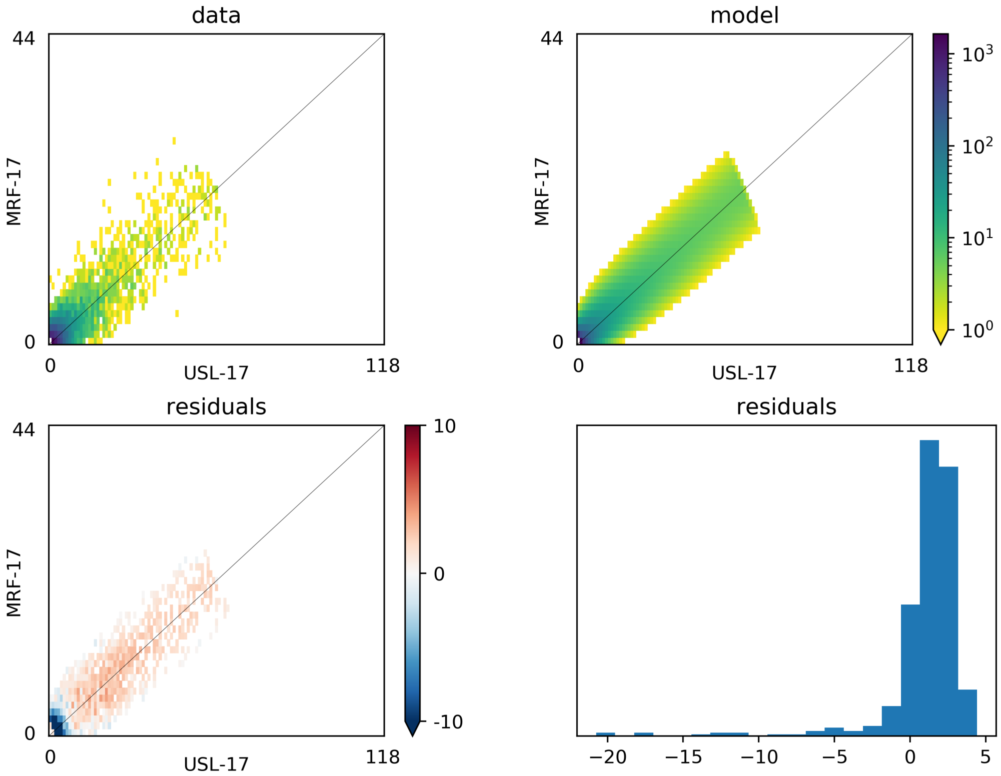


Figure S33: Summary of demographic model fit for MRF and USL. The top panels show the allele frequency spectra based on the observed data and best-fit model. The bottom panels show the associated residuals.

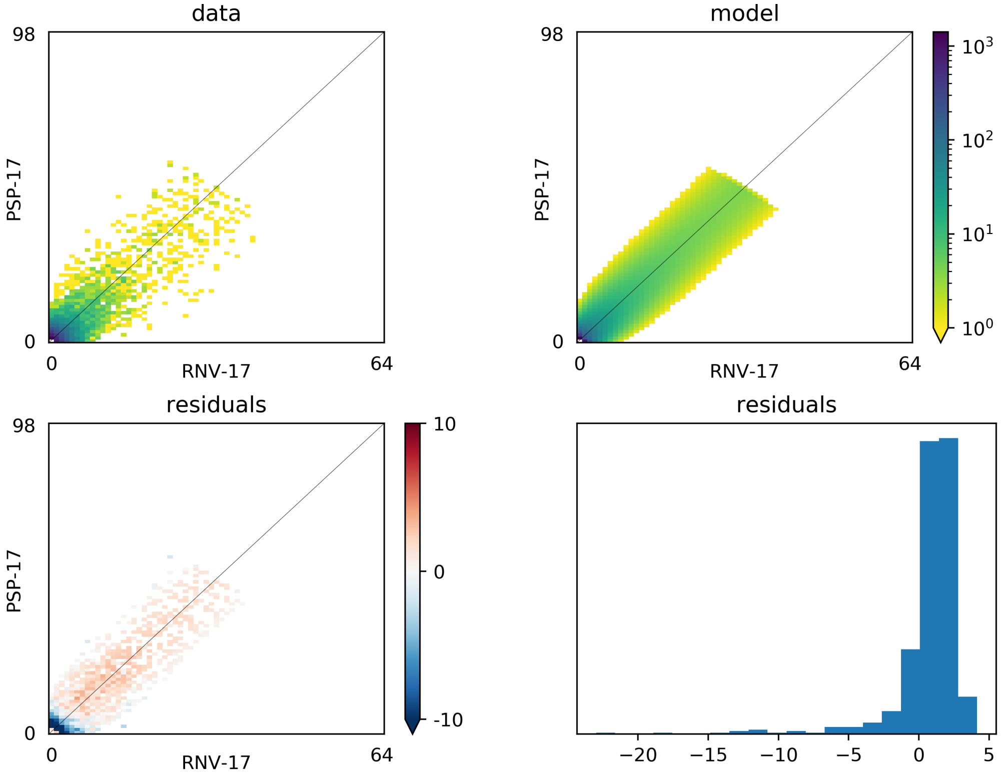


Figure S34: Summary of demographic model fit for PSP and RNV. The top panels show the allele frequency spectra based on the observed data and best-fit model. The bottom panels show the associated residuals.

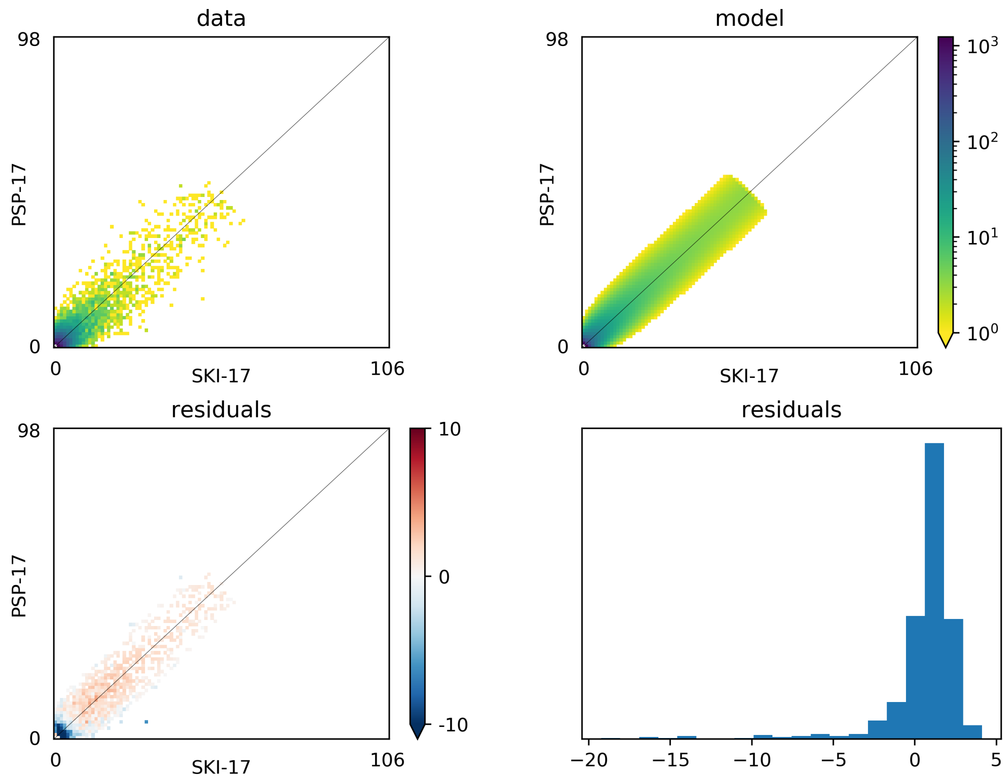


Figure S35: Summary of demographic model fit for PSP and SKI. The top panels show the allele frequency spectra based on the observed data and best-fit model. The bottom panels show the associated residuals.

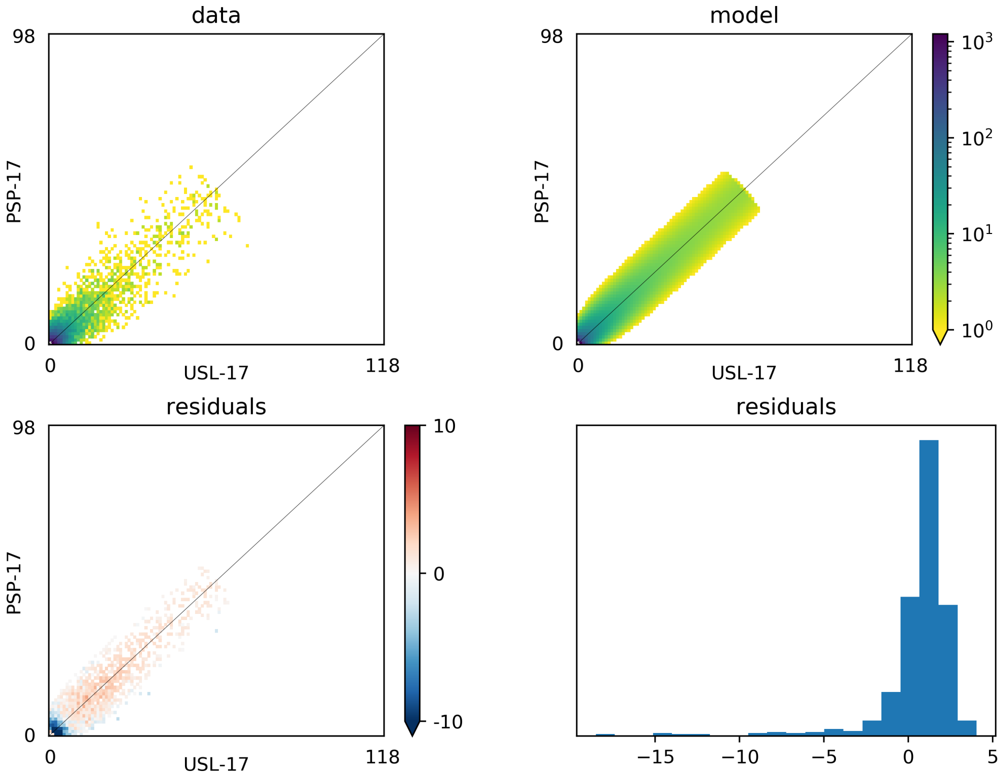


Figure S36: Summary of demographic model fit for PSP and USL. The top panels show the allele frequency spectra based on the observed data and best-fit model. The bottom panels show the associated residuals.

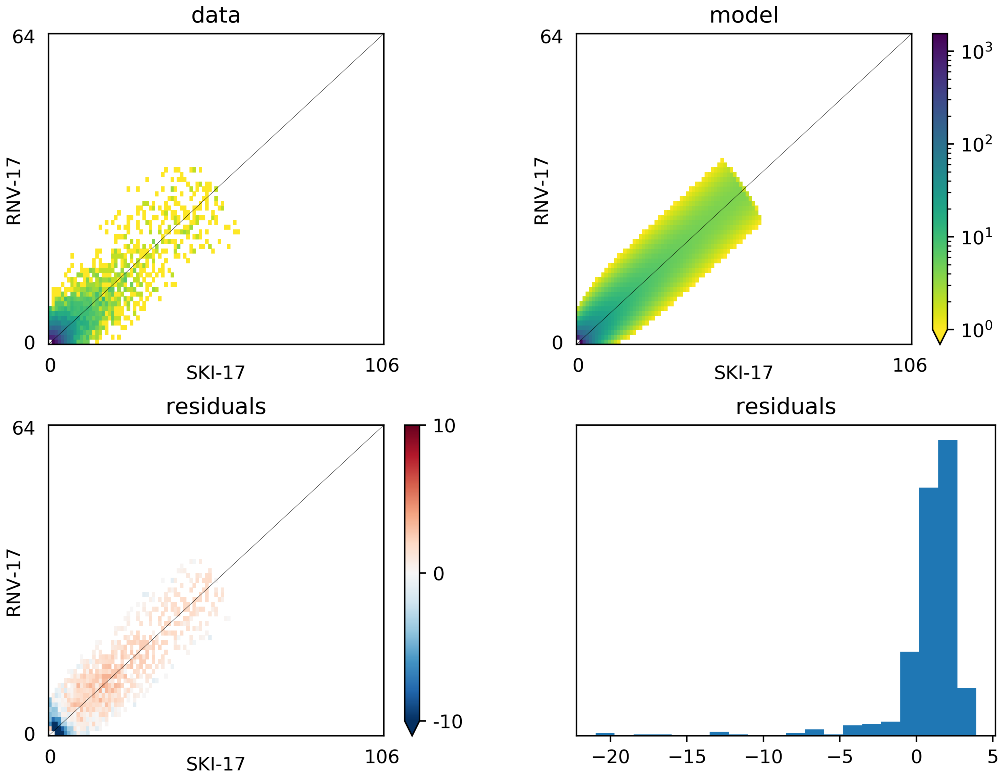


Figure S37: Summary of demographic model fit for RNV and SKI. The top panels show the allele frequency spectra based on the observed data and best-fit model. The bottom panels show the associated residuals.

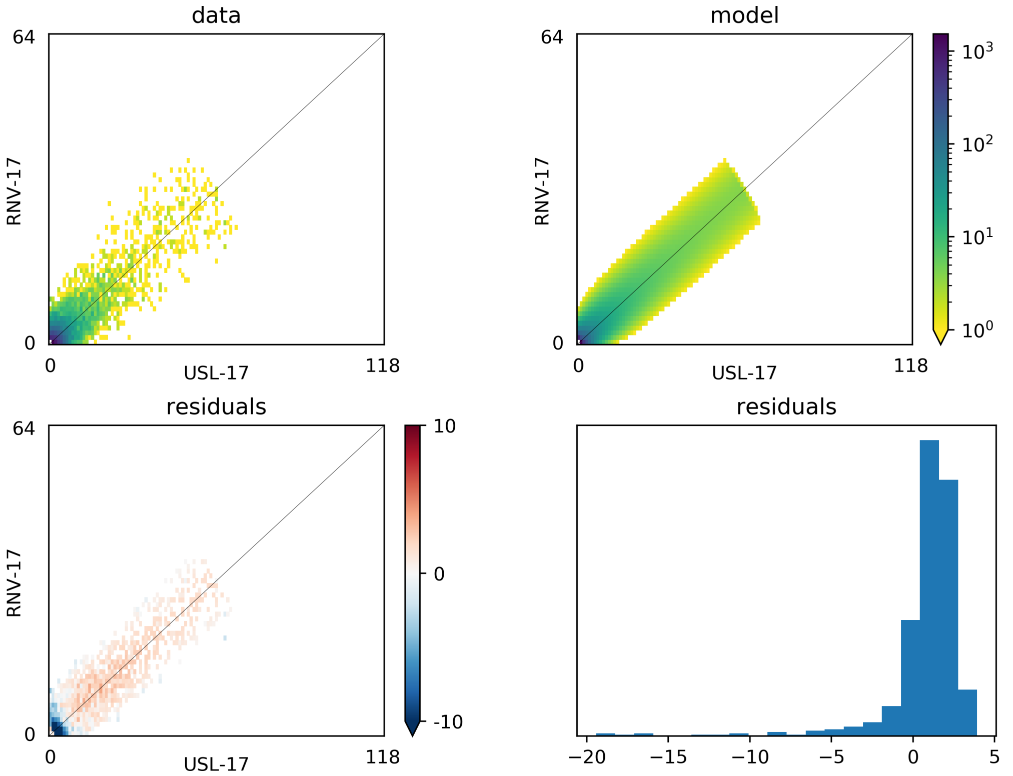


Figure S38: Summary of demographic model fit for RNV and USL. The top panels show the allele frequency spectra based on the observed data and best-fit model. The bottom panels show the associated residuals.

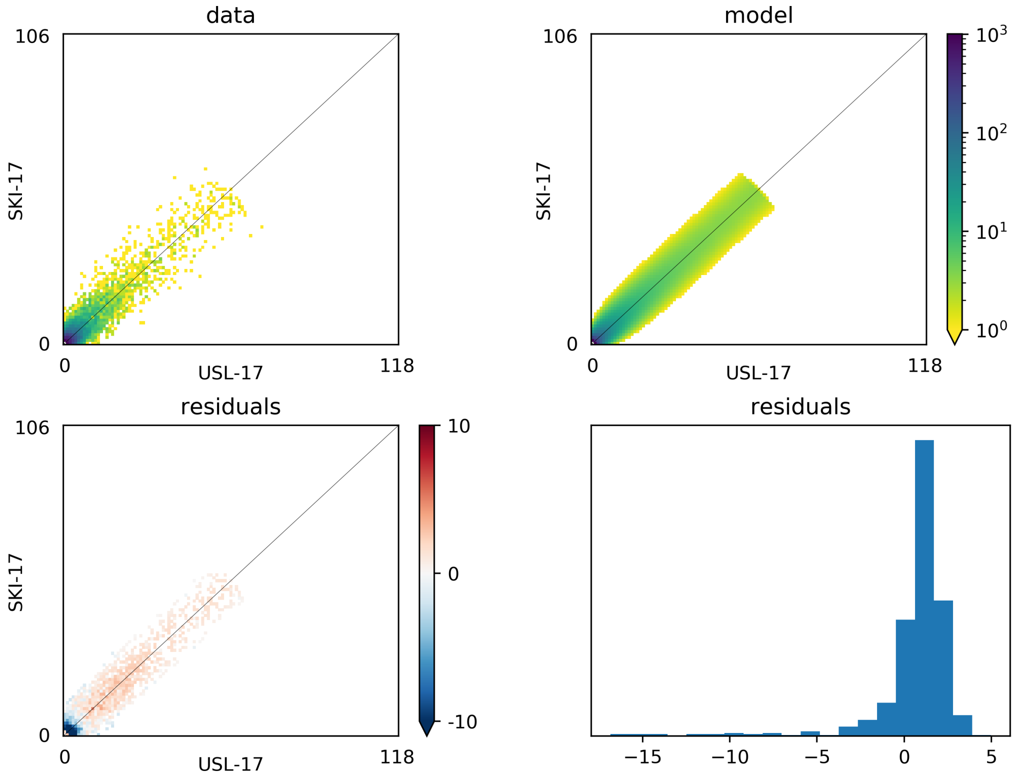


Figure S39: Summary of demographic model fit for SKI and USL. The top panels show the allele frequency spectra based on the observed data and best-fit model. The bottom panels show the associated residuals.
